# EMC is required for biogenesis and membrane insertion of Xport-A, an essential chaperone of rhodopsin-1 and the TRP channel

**DOI:** 10.1101/2021.02.01.429115

**Authors:** Catarina J. Gaspar, Lígia C. Vieira, John C. Christianson, David Jakubec, Kvido Strisovsky, Colin Adrain, Pedro M. Domingos

## Abstract

Insertion of hydrophobic transmembrane domains (TMDs) into the endoplasmic reticulum (ER) lipid bilayer is an essential step during eukaryotic membrane protein biogenesis. The ER membrane complex (EMC) functions as an insertase for TMDs of low hydrophobicity and is required for the biogenesis of a subset of tail-anchored (TA) and polytopic membrane proteins, including rhodopsin-1 (Rh1) and the TRP channel. To better understand the physiological implications of membrane protein biogenesis dependent on the EMC, we performed a bioinformatic analysis to predict TA proteins present in the *Drosophila* proteome. From 254 predicted TA proteins, subsequent genetic screening in *Drosophila* larval eye discs led to the identification of 2 proteins that require EMC for their biogenesis: farinelli (fan) and Xport-A. Fan is required for sperm individualization and male fertility in *Drosophila* and we now show that EMC is also required for these important biological processes. Interestingly, Xport-A is essential for the biogenesis of both Rh1 and TRP, raising the possibility that disruption of Rh1 and TRP biogenesis in EMC loss of function mutations is secondary to the Xport-A defect. We show that EMC is required for Xport-A TMD membrane insertion and increasing the hydrophobicity of Xport-A TMD rendered its membrane insertion to become EMC-independent. Moreover, these EMC-independent Xport-A mutants rescued Rh1 and TRP biogenesis in EMC mutants. Our data establish that EMC can impact the biogenesis of polytopic membrane proteins indirectly, by controlling the biogenesis and membrane insertion of an essential protein co-factor.

## INTRODUCTION

Membrane proteins comprise approximately 30% of the eukaryotic proteome and confer many essential functions to biological membranes^1,2^. These proteins contain hydrophobic transmembrane domains (TMDs) that must be inserted into the membrane of the endoplasmic reticulum (ER) through evolutionarily conserved molecular machineries. The vast majority of membrane proteins are inserted into the ER membrane through the co-translational pathway, in which proteins containing signal peptides or TMDs are recognized by the SRP (signal recognition particle) as they emerge from the ribosomes. Eventually, the ribosome-nascent chain complex is delivered to the Sec61 translocon, where membrane insertion takes place^3^.

The co-translational insertion pathway is the predominant mechanism for membrane protein insertion, but it is unable to deal with the biogenesis of a specific class of membrane proteins called tail anchored (TA) proteins, which lack a signal peptide and contain a single TMD at their C-terminus^4^. Consequently, their TMD remains shielded inside the ribosomal tunnel until the termination codon is reached, and TMD recognition can only occur post-translationally^5^. Accordingly, TA proteins utilize a dedicated conserved TMD recognition complex (TRC) pathway, which facilitates the targeted release of these proteins into the ER in eukaryotic cells, in a post-translational manner^6^. The crucial component of this pathway is the ATPase TRC40 (or in yeast Get3), which captures and shields TMDs of TA proteins^6–8^, until they are released to a receptor complex composed by WRB-CAML (yeast Get1-Get2), which mediates TMD insertion into the ER membrane^9,10^. Structural and biochemical studies have shown that this pathway displays a preference for TMDs of high hydrophobicity^11–13^.

Recently, the ER membrane protein complex (EMC) was identified as an insertase for TA proteins with TMDs of moderate to low hydrophobicity^13–15^. The EMC is a highly conserved, multi-subunit protein complex, with 9 subunits in mammals^16^. The EMC was initially described in yeast in a high throughput genetic screen for genes required for protein folding^17^ and its mammalian counterpart was later identified as part of the interaction network of the ER-associated protein degradation (ERAD) machinery^18^.

The EMC was subsequently shown to serve as an insertase for specific polytopic membrane proteins that contain a signal anchor sequence (SAS), including a subset of G-protein coupled receptors (GPCR)^19^. A variety of experiments suggested that the EMC coordinates the insertion of the first TMD of the GPCRs, after which subsequent TMD insertions are EMC-independent and require the Sec61 translocon, via a “handing-off” mechanism that remains to be fully elucidated^19^.

The structure of both the human and yeast EMC have recently been determined using cryo-electron microscopy (cryo-EM)^20–23^ and a model has been proposed for EMC-mediated co- and post-translational insertion of client proteins^20^. According to this model, a captured client protein is directed towards the membrane by the flexible cytosolic loop of EMC3, the insertase subunit of EMC; the EMC then presumably reduces the energetic cost of insertion by inducing a local thinning of the membrane and by arranging polar and positively charged residues within the bilayer. The client would then dissociate from EMC3, encountering EMC1’s β propellers, which could act as a scaffold for co-factor recruitment^25,22^.

The requirement for the EMC in the biogenesis of some TA proteins and the first TMD of some polytopic membrane proteins containing a SAS has been dissected *in vitro,* leading to a rationalization of how EMC can act mechanistically. However, the EMC has also been shown to be required for the biogenesis of multi-pass membrane proteins enriched with “challenging” TMDs^24^. Indeed, the majority of proteins found to date to be affected by loss of EMC are neither TA nor SAS-containing membrane proteins, suggesting that much remains to be understood about EMC specificity for its client proteins^15,24^ These include the ABC transporter Yor1 in yeast^25,26^, acetylcholine receptor in *C. elegans*^27^, rhodopsins^28–31^, the transient receptor potential (TRP) channel in *Drosophila*^29^, ABCA1 in mice^32^ and mutant connexin32^33^. Furthermore, EMC loss of function has also been associated with defects in phospholipid trafficking^34,35^, cholesterol homeostasis^14^, autophagosome formation^36,37^, viral pathogenesis^38–42^, neurological defects^43^ and male fertility^44^. Although this diversity in phenotypes is still an area of investigation, several of these examples impact unrelated membrane proteins with multiple TMDs^45,46^ and many of these candidate EMC clients are neither TA nor SAS-containing proteins. An alternative possibility to EMC acting as an insertase for a broader range of TMD protein topologies is that some of these proteins may be “indirect” or “secondary clients” of the EMC, for example, proteins whose biogenesis or stability depends on a direct EMC client protein.

In this study, we interrogated the *Drosophila* proteome in an effort to bioinformatically identify TA proteins that could have a dependency on the EMC for their biogenesis. Using a *Drosophila* larval eye imaginal disc assay, we identified two EMC clients: fan (farinelli), which controls sperm individualization^47^ and Xport-A (e*x*it *p*rotein of *r*hodopsin and *T*RP-A)^48,49^, an essential chaperone for the biogenesis of both Rhodopsin-1 (Rh1) and the TRP (Transient Receptor Potential) channel, and their targeting to the rhabdomere, the light sensitive compartment of the photoreceptors. Interestingly, the biogenesis of Rh1 and TRP has been shown to be deficient in EMC mutant clones^29–31^. We generated a mutant of Xport-A (Xport-A4L) whose biogenesis proceeds independently of the EMC. Xport-A4L is able to rescue the expression of Rh1 and TRP in EMC mutant tissue, suggesting that the latter proteins are not direct clients of the EMC but rather, depend on EMC indirectly via Xport-A. Overall, our results suggest that EMC is required for sperm and photoreceptor differentiation in *Drosophila*, due to EMC’s role in the biogenesis of fan and Xport-A, respectively. Crucially, our data establish that EMC impacts the biogenesis of some multi-pass membrane proteins indirectly, by governing insertion of the cofactors they require for assembly and deployment. This paradigm expands the potential governance of the EMC to a greater portion of the membrane proteome.

## RESULTS

### EMC is required for the biogenesis of a subset of tail-anchored proteins

The EMC has been shown to function as an insertase for TA proteins with TMDs of moderate to low hydrophobicity^13^, but the identities of its clients have not been fully determined. To ascertain the range of EMC-dependent TA proteins, we began by screening the *Drosophila melanogaster* proteome for all possible TA proteins. Tail anchored proteins are defined by their cytosolic N-terminal domain that is anchored to the lipid bilayer by a single hydrophobic TMD proximal to the C-terminus^50^. We bioinformatically interrogated the proteome using prediction algorithms for: signal peptide, TMD and topology^51^ as well as subcellular localisation^52^ (Figure 1A). This analysis yielded a total of 254 candidate TA proteins in the *Drosophila* proteome, with predicted membrane localizations (Figure 1A, S1-A, supplementary file1). We subsequently characterized the distribution of different biophysical features such as TMD length, TMD hydrophobicity, tail length and charge (Figure S1-C-F). These analyses show that within the list of predicted TA proteins, those that are ER-localized tend to have higher TMD hydrophobicity and length than their mitochondrial and peroxisomal counterparts (Figure S1-C-F), which is consistent with published predictions for both human^53^ and *Arabidopsis^54^* proteomes.

**Figure 1.**
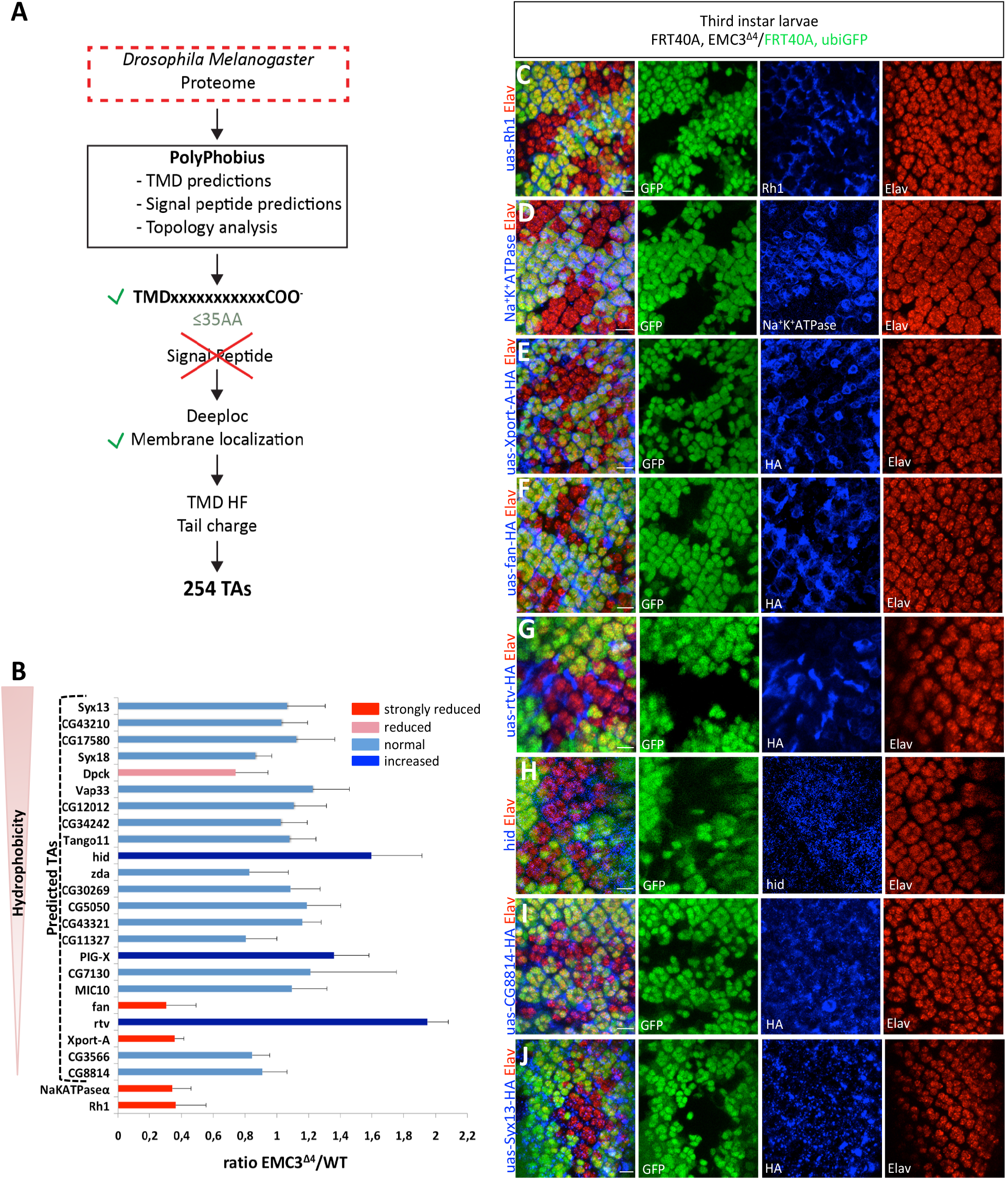
EMC is required for the biogenesis of a subset of tail-anchored proteins. (A) The *Drosophila* proteome was screened to identify TA proteins using the PolyPhobius^51^ algorithm within the TOPCONS^61^ web server. Proteins with a single TMD within 35 AAs or less upstream of the C terminus and lacking a signal peptide or mitochondrial targeting peptide were selected for further interrogation. The DeepLoc^52^ predictor of protein sub-cellular localization was used to eliminate soluble proteins from the list. These procedures yielded 254 proteins, which were scored fortheirtail charge and hydrophobicity using the “transmembrane tendency” score of Zhao & London^55^. (B) Asubset of the predicted TAproteins was screened by overexpression in *Drosophila* larval eye imaginal discs containing clones of a previously isolated EMC3 mutant allele, EMC3^Δ4^^29^. The ratio of fluorescence intensity for the tested TA proteins in EMC3 homozygous mutant cells over WT cells (EMC3^Δ4^/WT) was measured and plotted. Proteins were classified into four groups according to the ratio measured: normal (ratio 0.8-1.2; bars in blue), increased (ratio>1.2; bars in dark blue), reduced (ratio 0.8-0.5; bar in pink) and strongly reduced (ratio<0.5; bars in red). Error bars correspond to standard deviation (stdev). (C-J) All panels show *Drosophila* 3^rd^ instar larval eye imaginal discs with eyeless-Flippase-induced clones of cells homozygous for EMC3^Δ4 29^, labelled by the absence of ubiGFP (green). The red channel shows ELAV, which labels the nuclei of photoreceptor cells. All UAS constructs were expressed under the control of GMR-GAL4. (C) Expression of UAS-Rh1 (4C5, in blue) and (D) Na^+^K^+^ATPase (A5-C, in blue) is strongly reduced in EMC3^Δ4^ homozygous mutant cells. (E) Expression of UAS-Xport-A-HA (anti-HA, in blue) and (F) UAS-fan-HA (anti-HA, in blue) is strongly reduced in EMC3^Δ4^ homozygous mutant cells. (G) Expression of UAS-rtv-HA (anti-HA, in blue) and (H) hid (anti-hid, in blue) is increased in EMC3^Δ4^ homozygous mutant cells. (I-J) Normal expression of two TA proteins in EMC3^Δ4^ homozygous mutant cells. (I) UAS-CG8814-HA (anti-HA, in blue) is a TA protein containing a TMD of low hydrophobicity (J) UAS-Syx13-HA is a TA protein with high hydrophobicity TMD. Scale bars represent 10 μm.

The TMDs of most predicted TA proteins (~67%) exhibited low hydrophobicity^13^ (hydrophobicity value below 22 in the Zhao & London hydrophobicity scale^55^). This ratio was maintained in most sub-cellular localizations including the ER but with the exception of the Golgi membrane (75% proteins with high hydrophobicity) and peroxisome and mitochondrion membrane (100% of proteins with low hydrophobicity).

Next, we selected 23 candidate TA proteins to test for EMC dependency by monitoring their expression in EMC3 mutant clones in *Drosophila* larval eye imaginal discs (Figure 1B, 1C-J). This model enables candidate TA protein expression levels in WT and adjacent EMC3 mutant territories to be compared side-by-side, within the same tissue. Of the 23 proteins tested, approximately 87% had low predicted TMD hydrophobicity, were from within all predicted sub-cellular membrane localizations and had similar distribution of localization to the overall pool of predicted TA proteins (Figure S1-A-B). Candidate TA proteins were overexpressed under the control of GMR-GAL4 in larval eye imaginal discs containing EMC3 mutant (EMC3^Δ429^) clones. EMC3 is considered a “core” subunit of the complex, with mutations in EMC3 and other core subunits (EMC1, 2, 5 and 6) causing significant impairment of EMC functionality^13,14,46^. Candidate TA protein expression was assayed by immunofluorescence using antibodies raised against the respective proteins or, when no suitable antibodies existed, against an appended HA (Hemagglutinin) tag. Rh1 and the Na^+^K^+^ATPase α subunit were also evaluated (Figure 1C, D), as their expression has been reported previously to be EMC dependent^29–31^.

We evaluated TA candidate expression by determining the ratio of fluorescence intensity between the signal associated with the candidate TA protein in EMC3 mutant clones versus nonmutant (WT) clones (EMC3^Δ4^/WT) (Figure 1B). TA candidate expression was classified into four groups according to the EMC3^Δ4^/WT fluorescence intensity ratio: normal (ratio 0.8-1.2, bars in blue), increased (ratio>1.2, bars in dark blue), reduced (ratio 0.8-0.5, bar in pink) and strongly reduced (ratio<0.5, bars in red). Rh1 (Figure 1C) and the Na^+^K^+^ATPase α subunit (Figure 1D) exhibited strongly reduced expression in EMC3^Δ4^ mutant clones, consistent with previously published results in the adult/pupal eye^29–31^. Expression of Dpck was reduced in EMC3^Δ4^ mutant clones while rtv (Figure 1G), hid (Figure 1H) and PIG-X showed increased expression. Notably, two TA candidates, Xport-A (Figure 1E) and fan (Figure 1F), showed strongly reduced expression in EMC3^Δ4^ mutant clones compared to WT. Both proteins are predicted to localize to the ER membrane and contain TMDs of low hydrophobicity.

Of the 23 TA proteins tested, 17 exhibited normal ratios of expression, demonstrating that the GMR-GAL4 driven transcription of the candidate proteins is not affected by the loss of EMC function. Examples of TA proteins with TMDs of low hydrophobicity (CG8814 – Figure 1I) and high hydrophobicity (Syx13 – Figure 1J) showing normal EMC3^Δ4^/WT fluorescence intensity ratios, could also be found.

### EMC is required for sperm differentiation in *Drosophila*

Our screen identified the predicted TA protein fan, whose expression was defective in EMC3^Δ4^ homozygous mutant clones (Figure 1-F). As fan reportedly plays an important role in sperm individualization and male fertility^47^, we asked whether the EMC was also required for male fertility in *Drosophila*. We selected RNAi lines targeting different EMC subunits (EMC1, EMC3, EMC4, EMC5, EMC6, EMC7, EMC8/9) and mated adult males in which individual EMC subunits were knocked down, with wild-type virgin females (Figure S2A-B). We counted the mean number of progeny obtained from these crosses and observed a statistically significant reduction for all RNAi lines of EMC5 and EMC6, and for one RNAi line of EMC1 (Figure S2-A). We analysed sperm vesicle size and observed that, in accordance with the reduced male fertility, all EMC5 and EMC6 RNAi lines tested exhibited a statistically significant reduction in sperm vesicles (Figure S2-C), which correlates with a reduction in sperm production^47,56^. Furthermore, the seminal vesicles of flies expressing EMC5_2 RNAi line appeared devoid of mature sperm, as the needle shaped nuclei of mature sperm were absent (Figure S2-D). Finally, we tried to address the cause of male fertility in an EMC5 RNAi line, observing that while staining with an antibody against active Caspase-3 was present in multiple cystic bulges (cb) in wild-type testis, this staining was deficient in EMC5 RNAi testis (Figure S2-E), demonstrating sperm differentiation defects, as previously shown^57^.

### EMC3 is required for the expression of Xport-A, but not Xport-A4L

Although the specificity of EMC for its clients remains to be fully delineated, the EMC exhibits a preference for TA proteins with low TMD hydrophobicity and/or containing polar/charged amino acid residues^13–15^. As the Xport-A TMD has a reduced hydrophobicity due to the presence of one polar and multiple charged residues, an Xport-A TMD with increased hydrophobicity would be expected to relieve its EMC dependency. To that end, we performed site-directed mutagenesis on the Xport-A TMD to substitute the four most hydrophilic amino acids with leucine, creating the mutant Xport-A4L (Figure 2A). As hypothesized, whereas WT HA-Xport-A was dependent on the EMC (Figure 2B), expression of HA-Xport-A4L (Figure 2C) or Xport-A4L-HA (Figure 2D) no longer required EMC3 in larval eye imaginal discs. Importantly, the levels of the Na^+^K^+^ATPase control protein remained defective in EMC3 mutant clones (Figure 2D), confirming the presence of “bona fide” EMC3 mutant clones in the eye disc. A similar impact of the Xport-A4L mutant was observed in experiments carried out in adult retinas (Figure 2 E,F). Altogether, these results indicate that increasing the hydrophobicity of the Xport-ATMD renders its expression independent of the EMC and that tagging Xport-A and Xport-A4L at either the N- or C-terminus yields identical results.

**Figure 2.**
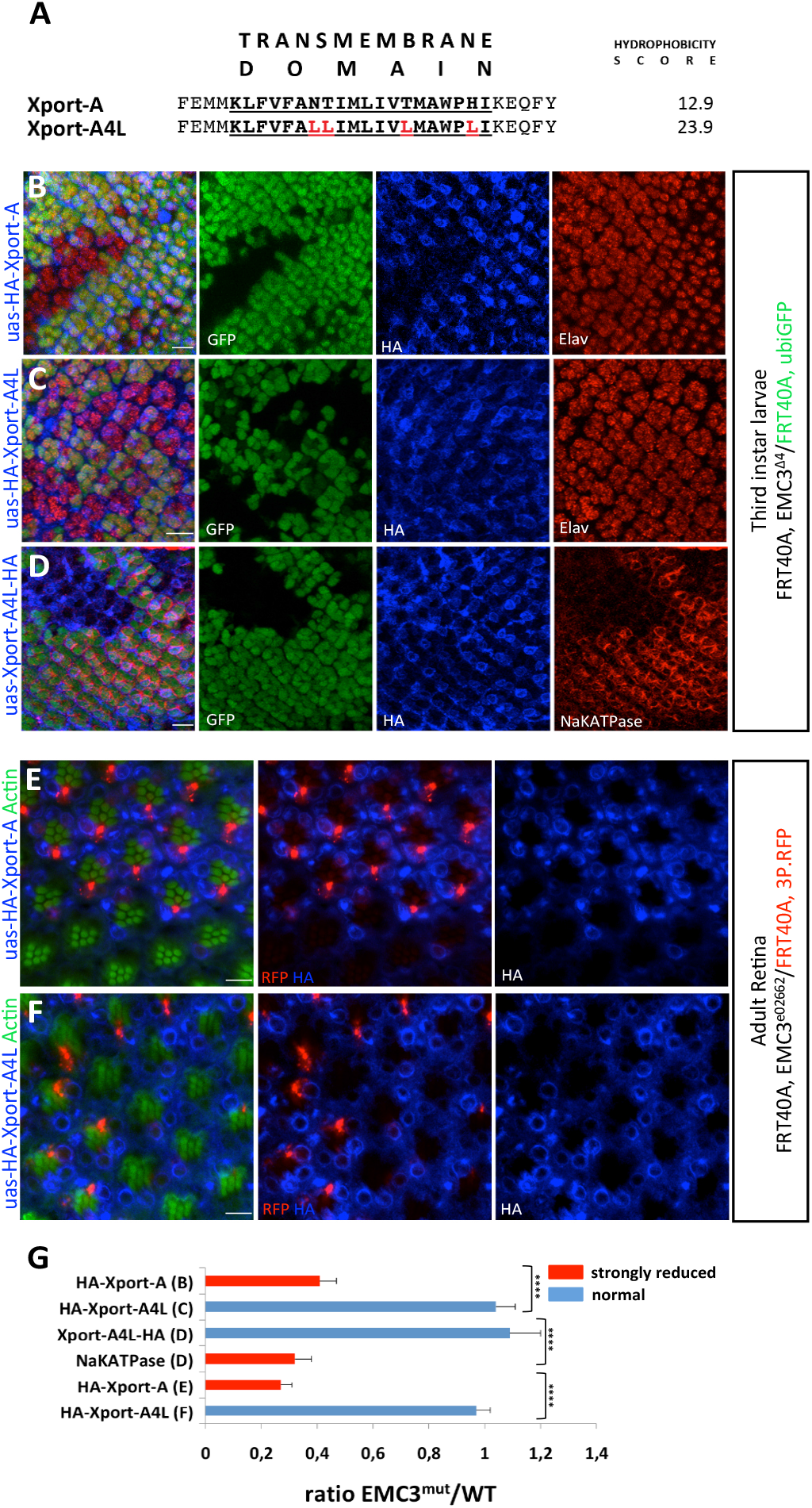
EMC3 is required for the expression of Xport-A, but not Xport-A4L. (A) The TMD of Xport-A was identified using TOPCONS^61^ and is in bold/underlined, together with the immediate flanking residues. Red residues indicate mutations made in the Xport-ATMD to increase its hydrophobicity, creating the Xport-A4L mutant. To the right of each sequence is the “transmembrane tendency” score, calculated according to Zhao & London^55^. (B-D) – Immunostaining of 3^rd^ instar larval eye imaginal discs with eyeless-Flippase-induced clones of cells homozygous for EMC3^Δ4^, labelled by the absence of ubiGFP (green) shows reduced expression of (B) UAS-HA-Xport-A (anti-HA, in blue) but normal expression of (C) UAS-HA-Xport-A4L (anti-HA, in blue) or (D) UAS-Xport-A4L-HA (anti-HA, in blue) in EMC3 mutant homozygous cells. ELAV (red in B and C) marks photoreceptor cells and Na^+^K^+^ATPase (red in D) acts as a positive control for the presence of EMC3^Δ4^ homozygous mutant clones. The UAS constructs were expressed under the control of GMR-GAL4. (E-F) Immunostaining of mosaic adult retina with eyeless-Flippase-induced clones of cells homozygous for EMC3^e02662^, labelled by the absence of RFP (red), show loss of (E) UAS-HA-Xport-A (anti-HA, in blue) in EMC3 mutant homozygous cells. (F) Expression of UAS-HA-Xport-A4L (anti-HA, in blue) is normal in EMC3^e02662^ homozygous mutant clones. The rhabdomeres are stained for actin (Phalloidin, in green). Scale bars represent 10 μm. (G) The ratio of fluorescence intensity in EMC3 homozygous mutant cells over that of WT cells (EMC3^mut^/WT) was measured and plotted. Proteins were classified into two groups according to the ratio measured: normal (ratio 0.8-1.2; bars in blue) and strongly reduced (ratio<0.5; bars in red). Error bars correspond to standard deviation (stdev). Significance was determined by Welch’s t-test: ****p ≤ 0.0001.

### EMC is required for the biogenesis of Xport-A in mammalian cells

We next sought to confirm the dependence of Xport-A biogenesis on the EMC using an assay independent of *Drosophila.* We cloned the TMDs of Xport-A, Xport-A4L and the validated EMC client squalene synthase (SQS)^13,14^, individually into the dual fluorescent reporter GFP-P2A-RFP (Figure 3A). When translated in mammalian cells, this mRNAwill generate two different products due to peptide bond skipping induced by the P2A site: GFP and RFP-SQS, RFP-Xport-A or RFP-Xport-A-4L. A stable GFP serves as a reporter for translation of the construct that enables quantitative read-out for protein stability through fluorescence ratios^19^. When the RFP:GFP ratio is reduced, it reflects post-translational degradation of the RFP-client protein fusion, presumably due a failure to integrate into the ER membrane. In human U2OS cells lacking the core subunit EMC5 (ΔEMC5)^13,14^, transiently expressed GFP-P2A-RFP-Xport-A exhibited a reduced RFP:GFP ratio when compared to WT U2OS cells (Figure 3C). Induced expression of EMC5 in ΔEMC5 cells restored the EMC and resulted in a RFP:GFP ratio that was comparable to WT cells (Figure 3C). Similar results were obtained from cells lacking the core subunit EMC6 (ΔEMC6, Figure 3F). In fact, Xport-A behaved comparably to SQS (Figure 3B and E), a TA protein involved in cholesterol homeostasis whose biogenesis has been established as EMC dependent^13,14^. Importantly, there was no difference in the measured RFP:GFP ratios following expression of GFP-P2A-RFP-Xport-A4L in ΔEMC5 or ΔEMC6 cells, when compared to either WT or EMC rescue cell lines (Figure 3D-G). These results indicate that the EMC is important for post-translational stability of Xport-A and that this effect relies on the low hydrophobicity of its TMD.

**Figure 3.**
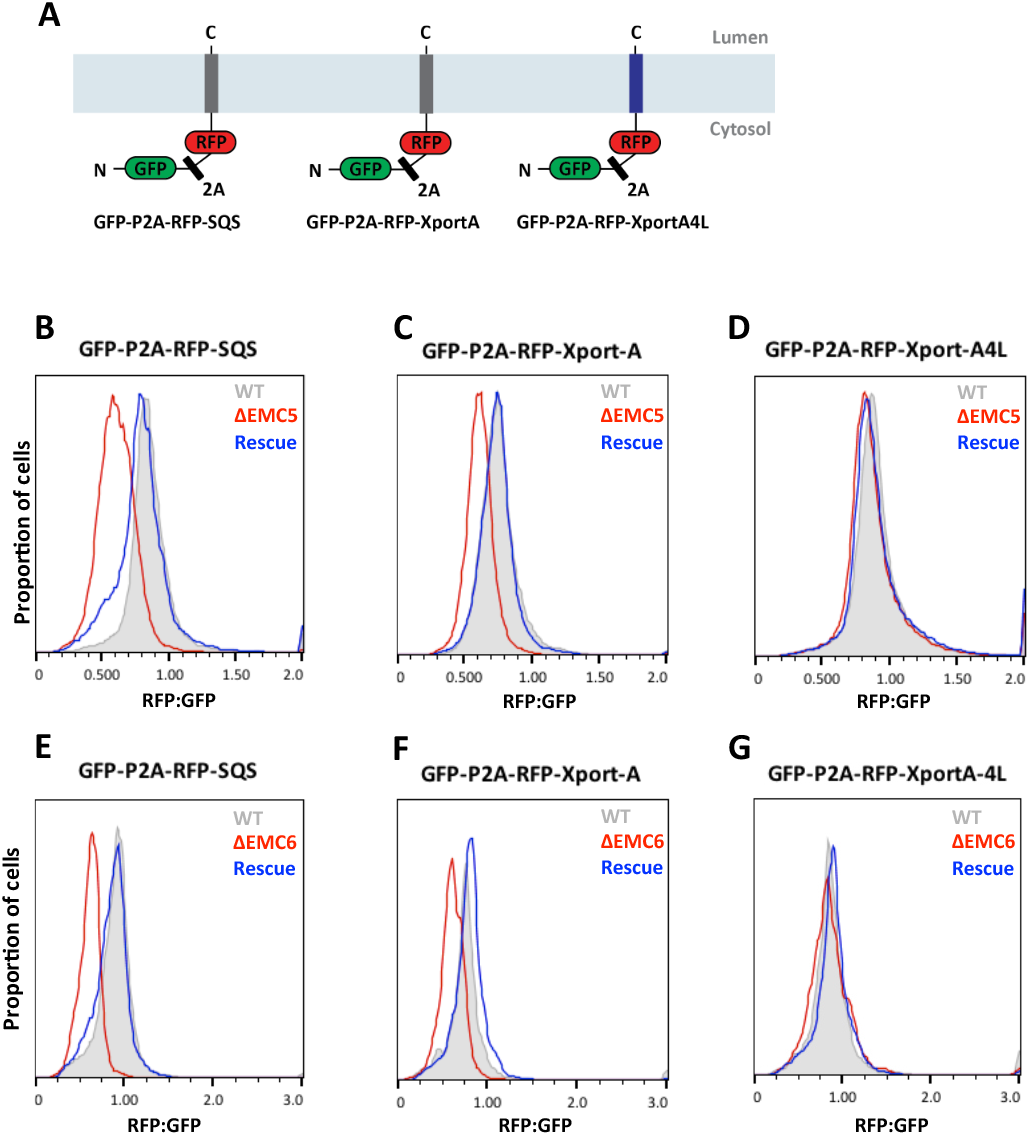
EMC is required for the biogenesis of Xport-A in mammalian cells. (A) Schematic representation of the reporter constructs used for analysis of protein biogenesis by flow cytometry. All constructs contain an N-terminal GFP and a C-terminal RFP, separated by a viral 2A peptide that mediates peptide bond skipping. Changes in the stability of SQS, Xport-A and Xport-A4L fused to RFP change the RFP:GFP fluorescence ratio. (B-D) Histograms of flow cytometry data monitoring the fluorescence protein ratio in the indicated U2OS cell lines for each construct. “EMC5KO+EV” indicates a knockout of EMC5 stably harbouring empty vector, while ‘‘EMC5KO+EMC5” indicates EMC5KO cells rescued by inducible re-expression of a stably integrated EMC5. (E-G) Histograms of flow cytometry data monitoring the fluorescence protein ratio in the indicated U2OS cell lines for each construct. “EMC6KO+EV” indicates a knockout of EMC6 stably harbouring empty vector, while ‘‘EMC6KO+EMC6” indicates EMC6KO cells rescued by re-expression of a stably integrated EMC6.

### EMC6 is required for membrane insertion of Xport-A TMD

Xport-A is a tail-anchored protein and its TMD must be inserted post-translationally into the ER membrane. To monitor insertion efficiency in the ER membrane, we cloned the TMDs from Sec61β, SQS, and Xport-A into a common cassette containing a C-terminal opsin tag (Figure 4A). The opsin tag, if exposed to the ER lumen, is able to accept N-linked oligosaccharides, serving as an indicator of successful TMD insertion into the membrane^58^. As predicted by our *Drosophila* results (Figures 1 and 2), the Xport-A TMD reporter is glycosylated in WT cells, but fails to be glycosylated in ΔEMC6 cells (Figure 4D), behaving similarly to HA-SQS-opsin (Figure 4C), as previously reported^13,14^. In contrast, the Sec61β reporter remains fully glycosylated in ΔEMC6 cells (Figure 4C), consistent with its EMC-independent insertion pathway^13^.

**Figure 4.**
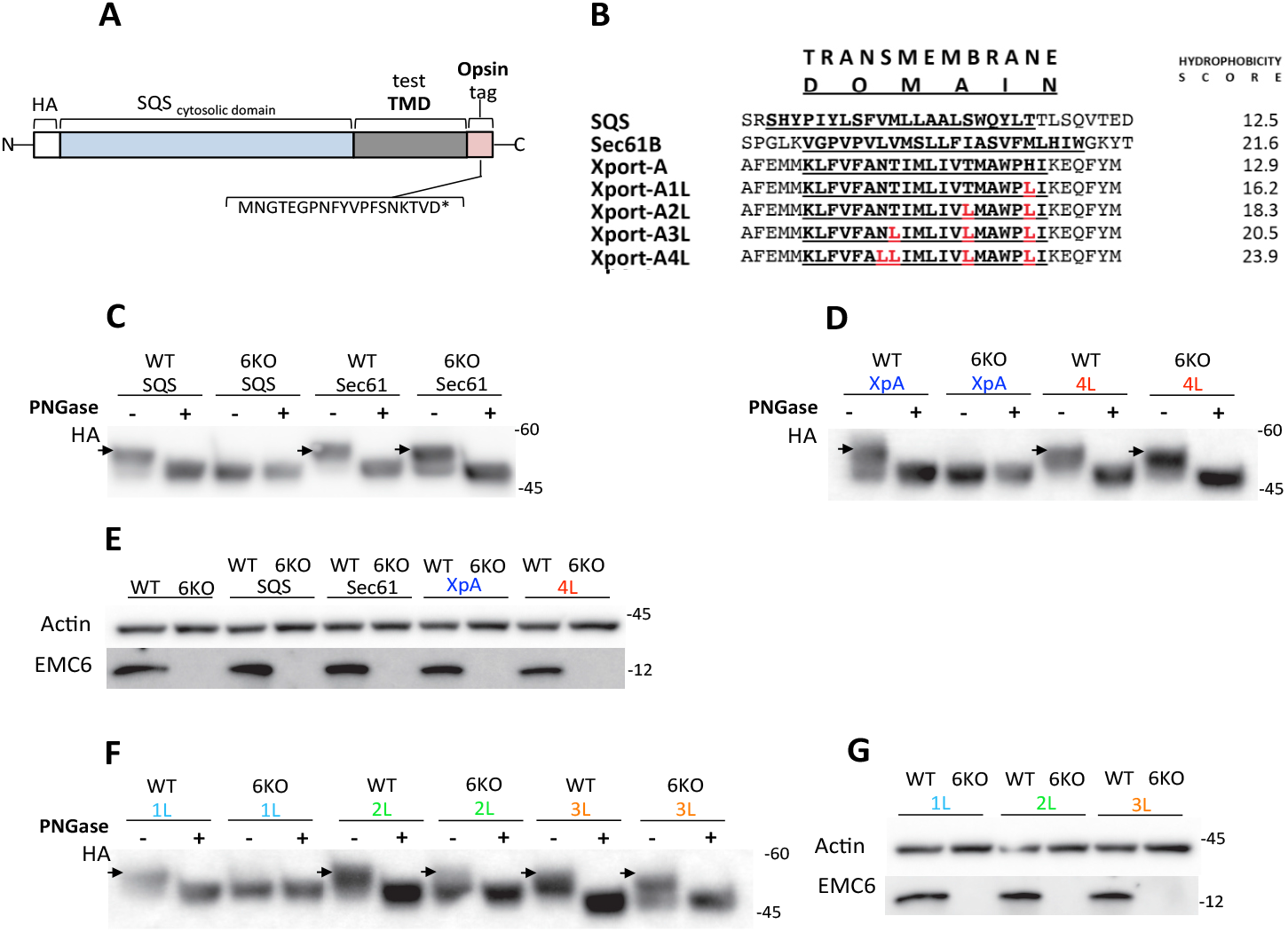
EMC6 is required for membrane insertion of Xport-A TMD. (A) Schematic representation of the constructs used to monitor insertion efficiency in the ER membrane. All constructs contain a HA-tagged SQS cytosolic domain fused to a varying TMD sequence, which is C-terminally fused to an opsin tag (sequence indicated). Glycosylation of the C-terminal opsin tag acts as an insertion readout. (B) Sequences of the TMD regions used for cloning and the immediate flanking residues are represented. The regions defined as TMDs were determined using TOPCONS61 and are in bold/underlined. Red residues indicate mutations made in the Xport-A TMD to increase its hydrophobicity. To the right of each sequence is the “transmembrane tendency” score of Zhao & London^55^. (C-D, F) Western-Blot of WT and EMC6KO (6KO) U2OS cell lines transiently transfected with the opsin-tagged constructs probed for HA. When indicated, denatured samples were treated with PNGase to confirm glycosylation. Glycosylated proteins shift upwards in the gel and are indicated by arrows (□). (E-G) Western-Blot of WT and EMC6KO (6KO) U2OS cell lines transiently transfected with the opsin-tagged constructs probed for Actin and EMC6.

Next we sequentially mutagenized the TMD of Xport-A to substitute the four most hydrophilic AAs with leucine (Figure 4B). By progressively increasing the hydrophobicity of Xport-A TMD, we attempted to identify a TMD hydrophobicity threshold for EMC recognition. A single point mutant of the Xport-A TMD (Xport-A1 L) retained its dependence on the EMC for membrane insertion (Figure 4F), but subsequent mutations (Xport-A2L, -A3L) were increasingly glycosylated in ΔEMC6 cells (Figure 4F). With four leucine mutations in the Xport-A TMD (Xport-A4L), insertion became entirely EMC independent, as indicated by the glycosylation pattern equivalent to what is observed in WT cells (Figure 4D). This finding supports the existence of a TMD “hydrophobicity threshold” that directs clients to insert in the ER membrane via the EMC.

### Expression of Xport-A4L can rescue Rhodopsin-1 and TRP biogenesis defects in EMC3 mutant clones

Xport-A serves as an essential chaperone during the biogenesis and maturation of both Rh1 and TRP; proteins that are essential for light sensing within the rhabdomere^48,49^. The protein levels of both Rh1 and TRP are significantly reduced in Xport-A loss of function mutations^48,49^, as well as in EMC mutations^29–31^. Our demonstration that Xport-A is a client protein of the EMC, raised the possibility that loss of TRP and Rh1 observed in EMC mutants could instead be attributable to failed membrane insertion and biogenesis of Xport-A. To test this hypothesis, we compared the capacity of Xport-A and Xport-A4L to rescue expression of Rh1 and TRP in EMC mutant clones. While Xport-A overexpression in EMC3 mutant clones was unable to rescue Rh1 (Figure 4-A), we did find that Xport-A4L expression was sufficient to restore Rh1 protein levels in EMC3^Δ4^ clones (Figure 4-B). Likewise, Xport-A4L rescued TRP expression in cells homozygous for a hypomorphic mutation of EMC3 (EMC3^e02662^, Figure 4D). Together, these data suggest that Rh1 and TRP biogenesis and maturation are not directly dependent on the EMC, but rather appear to be indirectly dependent on it, via the membrane insertion of Xport-A.

## DISCUSSION

Understanding the full scope of client proteins that depend on the EMC for biogenesis and maturation is yet to be fully appreciated. Here we describe an approach to predict, screen and validate candidate EMC client proteins which we apply to the *Drosophila* tail-anchored (TA) proteome. From a subset of TA candidate proteins with low predicted hydrophobicity scores, we identified fan and Xport-A as novel clients of the EMC. Taken together with previous studies that identified SQS, a key enzyme in sterol biogenesis^13,14^ and ZFPL1, a zinc-finger containing protein required for efficient ER to Golgi transport/integrity of the *cis-Golgi*^15,59^, our observations emphasize the importance of the EMC as an insertase for a subset of TA proteins whose insertion is independent of the GET/TRC40-complex. Taken together with previous studies^13–15^, our data also demonstrate conservation of the EMC’s ability to recognize TA proteins with reduced TMD hydrophobicity caused by the presence of charged/polar residues. Notably, the EMC appears to have additional levels of selectivity TA client proteins. Even though our predictions identified multiple candidate proteins with reduced TMD hydrophobicity, only a small fraction of these were confirmed as clients in our genetic screen in *Drosophila*. Future studies will be required to dissect the precise basis whereby the presence, nature and position of non-hydrophobic residues confers EMC dependency.

Fan is a member of VAP (VAMP associated protein) family of proteins that bind OSBPs (Oxysterol binding proteins) and play a role in sterol trafficking, including in the “individualization complex”, a cellular structure that promotes the membrane reorganization process (sperm individualization) that is crucial for the production of mature, functional sperm. Consistent with the role for fan in sperm individualization, depletion of several independent EMC subunits in testis (particularly EMC5, EMC6) render male flies infertile, with individualization defects similar to those reported for ablation of fan or OSBP family proteins^47^. Our identification of Fan as an EMC client, further strengthens the connections between the EMC and sterol homeostasis, established by the identification of SQS and SOAT1 as EMC clients^13,14^.

The identification of Xport-A as an EMC client raises some interesting questions. The Xport locus, consisting of the bicistronic operon separately encoding the Xport-A and Xport-B proteins, is required for the stable expression and localization of Rh1 (a GPCR) and TRP (a calcium ion channel) to the rhabdomeres^48,49^. Loss of Xport proteins results in depletion of Rh1 and TRP, causing photoreceptor degeneration and visual impairment. Intriguingly, as highlighted above, several studies have reported the loss of Rh1, TRP and accompanying photoreceptor degeneration in EMC-mutant flies^29–31^. As polytopic membrane proteins, Rh1 and TRP could directly engage the EMC during their biogenesis. However, our identification of Xport-A as an EMC client suggested the possibility that EMC governance over Rh1 and TRP expression could be indirect. Such a model is supported by the results obtained with the Xport-A mutant engineered to be EMC independent (Xport-A4L), which is sufficient to rescue expression of both Rh1 and TRP in mutant clones lacking functional EMC (Figure 5). Moreover, in the heterologous setting of mammalian cells where neither Rh1 nor TRP are expressed, Xport-A remains an EMC client with no collateral impact on its biogenesis. Conceptually, a model where “third party” proteins are lost as an indirect consequence of impaired biogenesis of a bona fide EMC client protein, could help to explain the pleiotropic impact of loss on multiple classes of membrane proteins when EMC is ablated.

**Figure 5.**
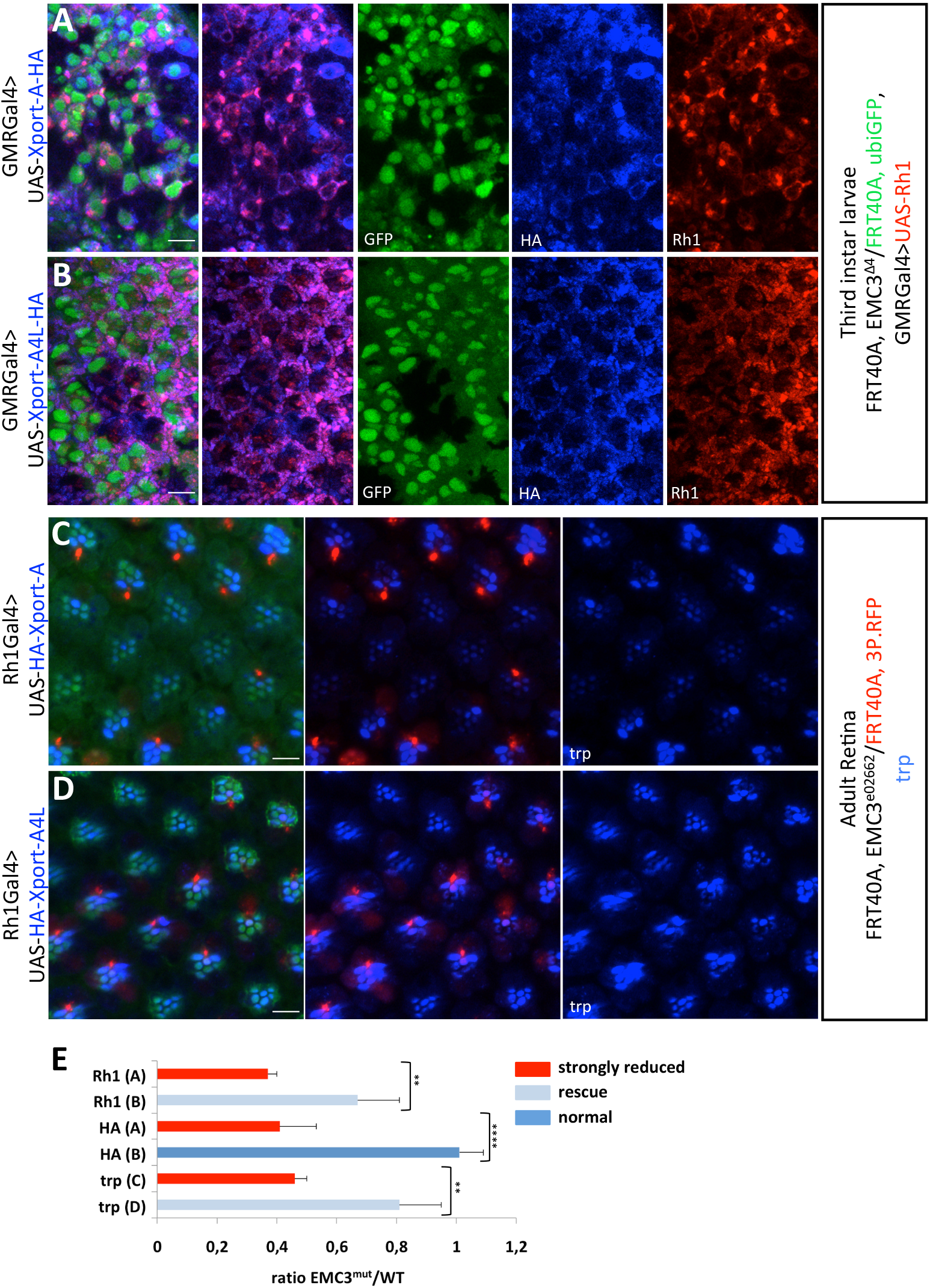
Expression of Xport-A4L can rescue Rhodopsin-1 and TRP biogenesis defects in EMC3 mutantclones. (A-B) Immunostaining of 3^rd^ instar larval eye imaginal discs with eyeless-Flippase-induced clones of cells homozygous for EMC3^Δ4^, labelled by the absence of ubiGFP (green) show loss of (A) UAS-Xport-A-HA (anti-HA, in blue) and UAS-Rh1 (4C5, in red). (B) Expression of UAS-Xport-A4L (anti-HA, in blue) and UAS-Rh1 (4C5, in red) is observed in EMC3^Δ4^ homozygous cells. The UAS constructs were expressed under the control of GMR-GAL4. Scale bars represent 10 μm. (C-D) Immunostaining of mosaic adult retinas with eyeless-Flippase-induced clones of cells homozygous for EMC3^e02662^, labelled by the absence of RFP (red), show loss of (C) TRP (Mab83F6, in blue) when UAS-HA-Xport-A is expressed. (D) Expression of TRP (Mab83F6, in blue) is observed in EMC3^e02662^ homozygous cells, when HA-Xport-A4L is expressed. The rhabdomeres are stained for actin (Phalloidin, in green). The UAS constructs were expressed under the control of Rh1-GAL4. Scale bars represent 10 μm. (E) The ratio of fluorescence intensity in EMC3 homozygous mutant cells over that of WT cells (EMC3^mut^/WT) was measured and plotted. Proteins were classified into three groups according to the ratio measured: normal (ratio 0.8-1.2; bar in dark blue), rescue (ratio 0.5-0.8, bars in light blue) and strongly reduced (ratio<0.5; bars in red). Error bars correspond to standard deviation (stdev). Significance was determined by Welch's t-test: **p≤ 0.01, ****p ≤

While an attractive hypothesis, some important caveats could apply to the model above. First, Rh1 is a GPCR, a class of proteins that contain an N-terminal SAS which have recently been proposed to be direct EMC clients^19^. Chitwood and colleagues showed that the EMC can mediate the insertion of the 1^st^ TMD of some SAS-containing GPCRs, after which insertion proceeds co-translationally in a Sec61-translocon-dependent manner^19^. However, this study did not investigate EMC requirement for *Drosophila* Rh1 membrane insertion. Moreover, expression levels of a Rh1 C-terminal truncation containing only the first TMD were uncompromised in EMC mutant *Drosophila* retinal cells^31^. In fact, only a truncation containing the first 5 of the 7 Rh1 TMDs was expressed at lower levels in EMC mutant tissue^31^. This study did, however, assessed expression levels of Rh1 mutants by immunofluorescence, only, and did not investigate either Rh1 TMD membrane insertion or topogenesis.

TRP contains 6 TMDs and a cytoplasmic N-terminus, which does not currently conform to the elucidated criteria for EMC client candidacy. An outstanding question is whether such non-canonical membrane protein clients are in fact engaged by the EMC for their biogenesis and if so, how? Several studies have reported the loss of multi-pass membrane proteins with loss of EMC functionality^15,23,24^. Often, these studies correlate EMC-dependence with abundance changes at steady-state, rather than monitoring EMC-dependent biogenesis directly. Consequently, it remains possible that loss of some of these putative EMC clients will occur indirectly, as we have demonstrated for Rh1 and TRP through Xport-A. Moreover, metabolic changes arising from disruption to EMC-dependent processes, such as sterol regulation, may also indirectly impact protein levels post-translationally through changes in intrinsic stability or homeostatic mechanisms linked to transcription. Care must be exercised when interpreting EMC-dependent biogenesis of complex multi-pass clients to ensure that, in fact, what is observed is a direct impact linked to TMD engagement.

Recently, studies from the Weissman lab identified features within the EMC structure that differentially impact TA protein clients when compared to multi-pass membrane proteins including TMEM97, a 4-TMD protein with a similar cytoplasmic N-terminal topology to TRP^23^. One interpretation is that EMC may be able to handle distinct classes of clients through different features. Miller-Vedam and colleagues proposed that the EMC may function not only as an insertase for TA proteins, but also as a “holdase” factor, that participates in the biogenesis of a wider class of polytopic membrane proteins via distinct mechanisms, which may also include the recruitment of client-specific chaperones^24^. It will be important to determine whether the impact of EMC on TRP is fully indirect, through its impact on Xport-A, or whether TRP directly engages the “holdase” function of EMC. The latter case could also involve recruitment of Xport-A to the EMC, to serve as a client-specific chaperone for Rh1/TRP, to aid in their maturation and/or release from the EMC and subsequent trafficking. Clearly, the biogenesis of membrane protein via the EMC has far reaching impact on organismal physiology and defining precisely the protein clients of EMC is indispensable to understand the selectivity of this complex.

## METHODS

### Prediction of TA proteins in *Drosophila* proteome

The *Drosophila melanogaster* proteome (UP000000803, release 2018_10_26) was downloaded from the UniProt website^60^. Canonical forms of protein sequences longer than 49 amino acids (AAs) and shorter than 6,001 AAs were selected. Topology (*i.e.*, localization of TMDs and nonmembrane regions) and presence of the signal peptide were predicted for each protein using the PolyPhobius^51^ predictor implemented within the TOPCONS consensus predictor^61^. The presence of the mitochondrial targeting peptide was predicted for each protein using TargetP 1.1^62^. Proteins without predicted signal and mitochondrial targeting peptides, lacking the keywords “SIGNAL” and “TRANSIT” in their UniProt annotation, and containing only a single predicted TMD localized close to the C-terminus (*i.e.*, followed by 35 or fewer flanking AAs) were selected. All thus selected protein sequences were analysed using the DeepLoc^52^ cellular localization predictor in order to further filter out soluble proteins. The TMD hydrophobicity was determined using the transmembrane tendency values for individual amino acids as defined by Zhao & London^55^. The C-terminal tail charge was calculated by adding +1 for arginine and lysine residues and −1 for aspartic and glutamic acid residues. The prediction of TA proteins was validated using experimentally identified proteins^13,63^. Characterization of the distribution of cell localizations and biophysical features was performed in GraphPad Prism v.9 and significance was evaluated using the Kolmogorov-Smirnov test.

### Fly stocks and crosses

Flies and crosses were raised with standard cornmeal, at 25°C under 12hr light/12 hr dark cycles.

Fly stocks used were obtained from multiple sources and are listed in Supplementary Table 1 and 2.

To determine if expression of predicted tail anchored proteins was dependent on EMC3, fly stocks carrying C-terminally HA-tagged constructs on the third chromosome (obtained from FlyOrf-Zurich ORFeome Project) were crossed with flies with an EMC3 mutant allele EMC3^Δ4^FRT40A^29^ on the second chromosome. The resulting flies (w; EMC3^Δ4^FRT40A; UAS-TA-HA) were then crossed with “eyFLP, GMRGAL4; ubiGFPFRT40A” to obtain EMC3^Δ4^ mosaic eye imaginal discs, expressing the predicted TA proteins. In the case of VAP33A, hid and Na^+^K^+^ATPase α-subunit, there was no need to express tagged fly lines, as these genes are endogenously expressed in eye imaginal discs and antibodies were available.

### Immunostaining and imaging

Larval eye imaginal discs and adult retina were dissected in 1x PBS, fixed in 4% PFA (paraformaldehyde) at room temperature (15min for eye imaginal discs, 30min for adult retina) and washed three times in PBT (1x PBS + 0.3% Triton X-100), 10 min each. For eye imaginal discs, primary antibodies were diluted in blocking buffer (1x PBS, 0.1% TX-100, 0.1% BSA, 250mM NaCl) and for adult retina, primary antibodies were diluted in 5% Fetal Calf Serum in PBT. Primary antibody incubations were performed overnight at 4°C under gentle agitation. Primary antibodies were as follow: mouse anti-ELAV (1:50) (9F8A9, Developmental Studies Hybridoma Bank (DSHB)), rat anti-ELAV (1:50) (7E8A10, DSHB), rat anti-HA (1:400) (7C9, Chromotek), mouse anti-Rh1 (1:50) (4C5, DSHB), mouse anti-Na^+^K^+^ATPase α subunit (1:25) (A5-C, DSHB), anti-VAP33A(1:50)(kind gift of Hugo Bellen), anti-hid (kind gift from Hyung Don Ryoo), rabbit anti-HA (1:40) (9110, Abcam), mouse anti-TRP (1:25) (Mab83F6, DSHB). Samples were washed three times with PBT and incubated with appropriate secondary antibodies (Jackson Immuno Research) for 2 hours at room temperature. Control staining of actin in the rhabdomeres of adult retina was performed with Phalloidin-FITC (1:500) (Abcam 235137). After rinsing three times with PBT, samples were mounted in Vectashield (Vector Laboratories). Adult eye samples were mounted in a bridge formed by two coverslips to prevent the samples from being crushed and analysed on a Zeiss LSM 880 confocal microscope. Larval eye imaginal discs were analysed on a Leica SP5Live confocal microscope.

Adult testis were dissected in 1x PBS and fixed in 4% PFA overnight at 4°C under gentle agitation. Following this, samples were washed three times in PBS, 5 min each and then treated with PBT at room temperature for 30 min. After three more washes with PBS (5 min each), testis were incubated with rabbit anti-cleaved caspase 3 (1:200) (Asp175, Cell Signaling) diluted in 0.1% PBT, overnight at 4°C. Samples were washed three times with PBS, 5 min each, and incubated with antirabbit Cy3 (1:200) (Jackson Immuno Research). After three more washes with PBS (5 min each), samples were incubated with DAPI, diluted in 0.1% PBT for 10 min. After rinsing three more times with PBS, samples were mounted in Vectashield and image acquisition was performed in Leica SP5Live.

For measurement of seminal vesicle size, testis were stained with DAPI, as described above and image acquisition was performed in Leica SP5Live. Imaging was performed at the widest portion of the seminal vesicle and area measurement was performed in Fiji. Characterization of distribution of seminal vesicle size was performed in Prism and significance was determined by Welch’s t-test.

### Quantification of protein expression in EMC3Δ4 mutant clones

Quantification was executed in Fiji^64^. Maximum intensity projections were performed on the raw confocal data to visualize EMC3^Δ4^ mutant clones. The three coloured images were split into single colour images. Channel 1 corresponded to ubiGFP, in which homozygous mutant clones were identified by the absence of fluorescence. Channel 2 corresponded to the fluorescence signal for the protein of interest, while channel 3, which showed ELAV staining, marked the photoreceptor cells. Segmentation was first performed on channel 3 to define the GMRGAL4 domain, and then on channel 1 to highlight patches with no detectable ubiGFP fluorescence, which coincided with EMC3^Δ4^ homozygous mutant cells. Both segmentations were performed using the Otsu thresholding method^65^. Following segmentation, the mathematical process “AND” was performed to intersect mutant patches delimited in the second segmentation with the GMRGAL4 domain identified in the first segmentation. EMC3^Δ4^ mutant patches in the GMRGAL4 domain were defined as ROIs. Mean grey values for the ROIs in channel 2 were measured (EMC3^Δ4^ mean) and compared to mean grey values of WT patches of comparable area in channel 2 (WT mean). A ratio of fluorescence intensity in EMC3^Δ4^ mutant cells/WT cells was calculated for each protein. The Fiji macro developed for this analysis can be found on GitHub (https://github.com/zeserrado-marques/Quantification-of-protein-expression-inside-dark-patches). For quantification, at least 6 mutant patches were quantified per eye imaginal disc, and at least three eye imaginal discs derived from distinct flies were used. Significance was evaluated using the Welch’s t-test in Graphpad Prism 9 software.

### Generation of HA tagged Xport-A transgenic flies

To generate transgenic flies expressing HA tagged Xport-A we PCR amplified the full open reading frame of Xport-A from plasmid pAC5.1-FLAG::XPORT-A^49^ (kind gift of Craig Montell). For N-terminally tagged Xport-A, we used the following primers: 5’-CCGGATTACGCCAAGCCGAAGAAATCGGCC-3’ and 5’-CGCAGATCTGTTAACGAATTCTTACTGCTGATGCTCCTCGC-3’. For C-terminally tagged Xport-A we used the following primers: 5’-GAATAGGGAATTGGGAATTCATGAAGCCGAAGAAATCGGCC-3’ and 5’-TCAGGGACGTCGTA CGGGTACTGCTGATGCTCCTCGCGAAAG-3’. For amplification of 3XHA tag, we PCR amplified 3XHA with following primers: 5’-TGAATAGGGAATTGGGAATTCATGTACCCGTACGACGTCCCTGA-3’ and 5’-TCTTCGGCTTGGC GTAATCCGGCACATCA-3’ for HA-Xport-A. For PCR amplification of 3XHA intended for Xport-A-HA, we used the following primers: 5’-GAAAGCTTTCGCGAGGAGCATCAGCAGTACCCGTACGACGTCCCTGA-3’ and 5’-CGCAGATC TGTTAACGAATTCTTAGCCGGCGTAATCCGGCACATCA-3’. After amplification of Xport-A and 3XHA inserts, we proceeded to perform a Gibson assembly^®^ (New England Biolabs) with plasmid pUASTattb (previously linearized with EcoRI).

Xport-A4L constructs were made by mutating hydrophilic AAs to Leucine, using KOD polymerase site-directed mutagenesis (Merck Millipore). Xport-A4L insert was then amplified by PCR with primers listed above and Gibson assembly was used to create N-terminal and C-terminal HA tagged pUASTattb constructs.

The resulting constructs were inserted into the *attp2* site by PhiC31 integrase mediated transgenesis at the Champalimaud Foundation transgenics facility.

### Cell culture

U2OS Flp-In^TM^ TRex™ cell lines describe previously^14^ were maintained in Dulbecco’s Modified Eagle Medium (DMEM)(Biowest) + 10% v/v fetal bovine serum (FBS) (Biowest) + L-glutamine (2mM) (GRiSP) + 1% v/v Penicillin-Streptomycin (Biowest) in 5% CO_2_ at 37°C. Stable expression cell lines, generated by flippase mediated site-specific integration, were continuously cultured in 100 μg/ml hygromycin B (Merck Millipore). Cells containing a stably expressed tetracycline-induced constructs were cultured in tetracycline-free FBS (Biowest), 100 μg/ml hygromycin, with their expression induced by addition of 1 μg/ml doxycycline (Sigma Aldrich).

### Plasmids and expression constructs

Xport-A and XportA4L dual fluorescent reporters were generated by first amplifying the TMD and flanking residues of Xport-A (AFEMM<u>KLFVFANTIMLIVTMAWPHI</u>KEQFYM) and Xport-A4L (AFEMM<u>KLFVFALLIMLIVLMAWPLI</u>KEQFYM) using the primers PFmcherry-xportTMD (GGCGGCATGGACGAGCGTTACAAGGCATTTGAAATGATGAAACTC) and PRmcherry-xportTMD (TTATGATCAGTTATCTAGATCCGGTTTACATGTAGAATTGCTCCTTGAT). Reporter plasmids were constructed from BamHI-linearized, dephosphorylated pcDNA5/FRT/TO-GFP-P2A-mcherry vector (kind gift from Manu Hegde, LMB-Cambridge) using Gibson assembly^®^ (New England Biolabs). The SQS fluorescent reporter GFP-P2A-mcherry-SQS has been reported previously^13^. TMD insertion reporters pcDNA5/ FRT/TO-HA:SQS_opsin_ and pcDNA5/FRT/TO-HA:SQS-Sec61□TMD^opsin^ have been reported previously^14^. To generate XportATMD^opsin^ (pcDNA5/FRT/TO-HA:SQS-XportATMD^opsin^) and hydrophobic TMD variants (1L/2L/3L/4L), geneblocks (IDT) containing each TMD sequence fused to the opsin tag were purchased. Each gene block contained flanking restriction sites for BamHI and XhoI that enabled subcloning into linearized pcDNA5/FRT/TO-HA:SQS.

### Flow cytometry

Analysis of protein post-translational stability was performed according to described methods^13^. Cells were transfected with fluorescent reporter constructs (1 μg total DNA/well, 6 well plate) using GenJet (SignaGen Laboratories) according to manufacturer’s instructions. The amount of fluorescent reporter for each protein of interest was titrated according to transfection efficiency and a control plasmid was cotransfected in order to maintain the total amount of DNA (1 μg/ well). The U2OS FlpIn Trex EMC5KO rescue cell lines were cultured in the presence of doxycycline (1μg/ml) for 24 hours prior to transfection, to rescue EMC5 expression. After transfection, cells were trypsinized, washed twice with PBS +2%FCS and stained with DAPI (to determine cell viability). Flow cytometry was performed on a BD LSR Fortessa X20 instrument, where 20,000 RFP-positive cells were collected. Data analysis was performed in FlowJo (v10). For quantification, cells were first gated for GFP (translation reporter) and mean RFP and GFP fluorescence intensities were determined. Then, RFP:GFP ratios were calculated and normalized to the ratio observed in WT cells.

### Transient transfection of TMD insertion reporters

1×10^6^ cells/6cm^2^ tissue culture plate were transfected with TMD insertion reporters (2.5 μg total DNA/6cm^2^ plate) using GenJet (SignaGen Laboratories) according to manufacturer’s instructions. The following day, cells were lysed.

### Immunoblotting

Cells were washed twice in ice-cold PBS and lysed in TX-100 lysis buffer (50mM Tris-HCl pH 7.4, 150mM NaCl, 1% Triton X-100, protease inhibitors) + 5mM EDTA. Following incubation on ice (10 min), lysates were centrifuged (top speed, 10min, 4°C). Protein concentrations were determined by Bradford and protein denaturation was performed by adding 3x LDS buffer + DTT and incubating the samples at 65°C (15 min). After denaturation, samples were digested with PNGaseF (New England Biolabs) for 2 hours at 37°C, according to manufacturer’s instructions.

Samples were fractioned by SDS-PAGE, transferred to PVDF membranes (Amersham Hybond) and probed with following antibodies: mouse anti-HA-HRP (1:1000) (12013819001, Roche) rabbit anti-EMC6 (1:300) (84902, Abcam) and rabbit anti-actin (1:2000) (8227, Abcam).

### Fertility assay

To examine fertility, individual adult males (3-days old) were mated to three wild-type virgin females in separate vials. All RNAi lines were expressed with bam-GAL4 driver. For each RNAi line, thirty individual males were mated to virgin females. The females were transferred after 7 days at 25°C to fresh vials. Progeny from the original vial and the first transfer vial were counted. Characterization of distribution of progeny/ female/day was performed in Prism and significance was determined by Welch’s t-test.

## Acknowledgements

We thank the flow cytometry facility at IGC, the microscopy facility at IGC (more specifically José Marques), the microscopy facility at ITQB-NOVA, the fly facility from IGC and the transgenics facility at Champalimaud Foundation – Centre for the Unknown. We thank the Developmental Studies Hybridoma Bank (DSHB) at the University of Iowa for antibodies and the Bloomington Drosophila Stock Centre for fly stocks. We thank Craig Montell for flies, plasmids and antibodies, Akiko Satoh for the EMC3 mutant flies, Manu Hegde for plasmids, Norbert Volkmar and Miguel Cavadas for discussions and suggestions, Nansi Jo Colley and Steve Britt for antibodies, Hugo Bellen for VAP33A antibody, Hyung Don Ryoo for UAS-Rh1 flies and hid antibody. We thank Sebastien Alfaiate for help with figure design. The project leading to these results has received funding from ‘la Caixa’ Foundation (ID 100010434), under the agreement <LCF/PR/HR17/ 52150018>.

## Competing interests

The authors declare no competing interests.

## Author contributions

Research design: all authors. TA protein prediction was done by DJ and KS. All experiments were performed by CJG, except the *Drosophila* male fertility experiments (LCV). Paper writing and editing: all authors.

## Data availability

The macro method developed for the quantification of protein expression in EMC3^Δ4^ mutant clones can be found on Github (https://github.com/zeserrado-marques/Quantification-of-protein-expression-inside-dark-patches).

## Abbreviations

AAs: amino acids
cb: cystic bulges
Cryo-EM: cryo-electron microscopy
EMC: ER membrane protein complex
ER: endoplasmic reticulum
ERAD: ER-associated protein degradation
fan: farinelli
GPCR: G-protein coupled receptor
HA: hemagglutinin
OSBP: oxysterol binding protein
PFA: paraformaldehyde
Rh1: Rhodopsin1
SAS: signal anchor sequence
SOAT1: sterol O-acyltransferase 1
SQS: squalene synthase
SRP: signal recognition particle
TA: tail anchored
TMD: transmembrane domain
TRC: TMD recognition complex
TRP: transient receptor potential
VAP: VAMP associated protein
Xport-A: exit protein of rhodopsin and TRP – A

---

**Figure S1.**
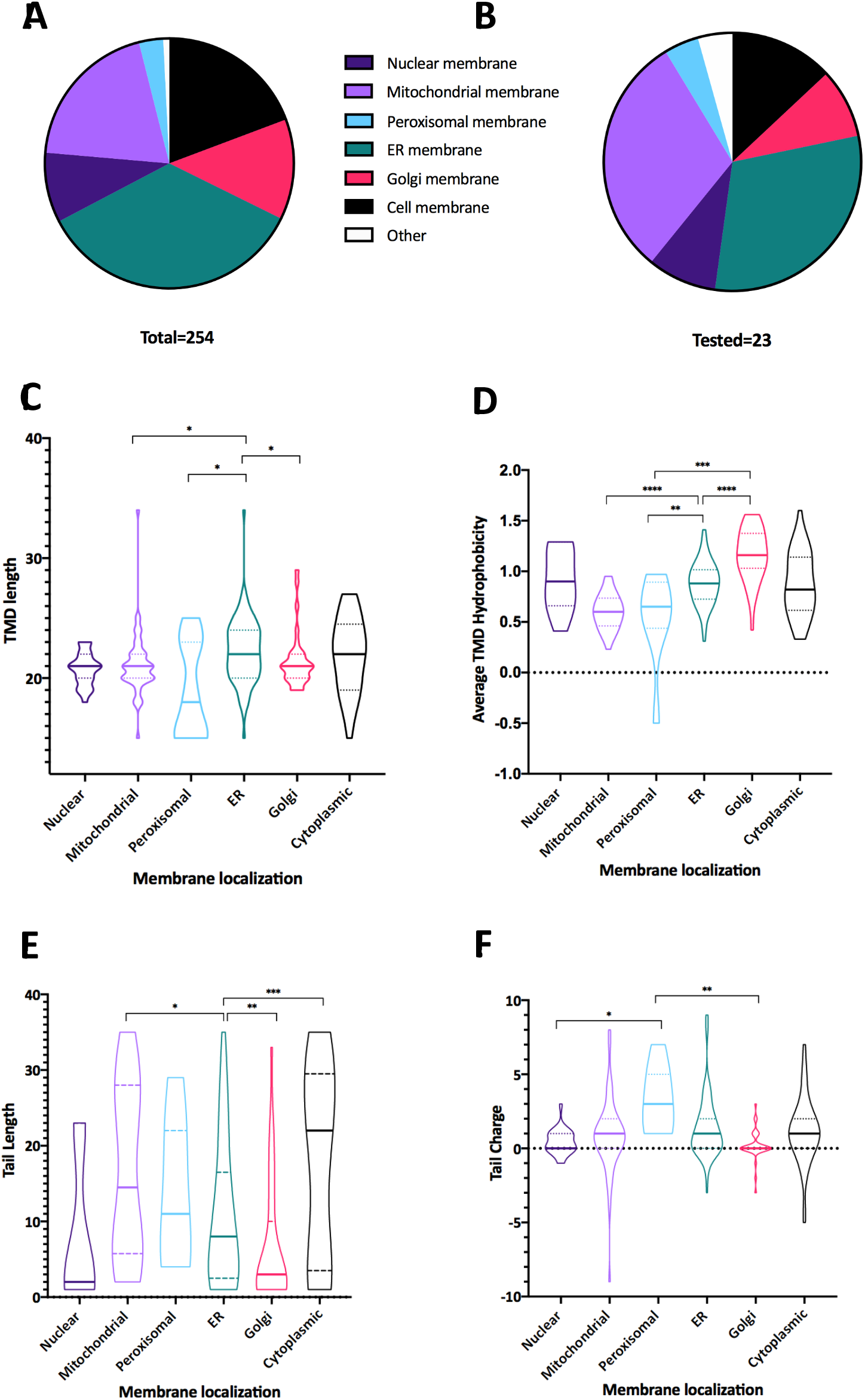
Characterization of the predicted TA proteins. (A) Pie chart showing the distribution of the cell localization in predicted TA proteins. The predicted TA proteins were predominately localized to the ER membrane (35%), mitochondrial membrane (~20%) and cell membrane (~20%). Other subcellular localizations predicted were Golgi membrane (~13%), nuclear membrane (9%) and peroxisome membrane (~3%). (B) Pie chart showing the distribution of the cell localization in tested TA proteins. TA candidates with all predicted sub-cellular membrane localizations were tested. The tested proteins had the following sub-cellular localizations: ER membrane (~30%), mitochondrion membrane (~30%), cell membrane (~13%), nuclear membrane (~8,7%), Golgi membrane (~8,7%) and peroxisome membrane (~4,3%). (C-F) Violin plots showing the distribution of biophysical features such as (C) TMD length, (D) average TMD hydrophobicity (TMD hydrophobicity in Zhao & London score, averaged to the number of AAs in each TMD), (E) tail length and (F) tail charge in TA proteins, categorized by different cellular localization. The mid-line in the violin plots represents the median of the population and the extremities below and above the median correspond to the minimum and maximum values. Significance was determined by Kolmogorov-Smirnov test: *p≤0.05, **p≤ 0.01, ***p ≤ 0.001, ****p ≤ 0.0001.

**Figure S2.**
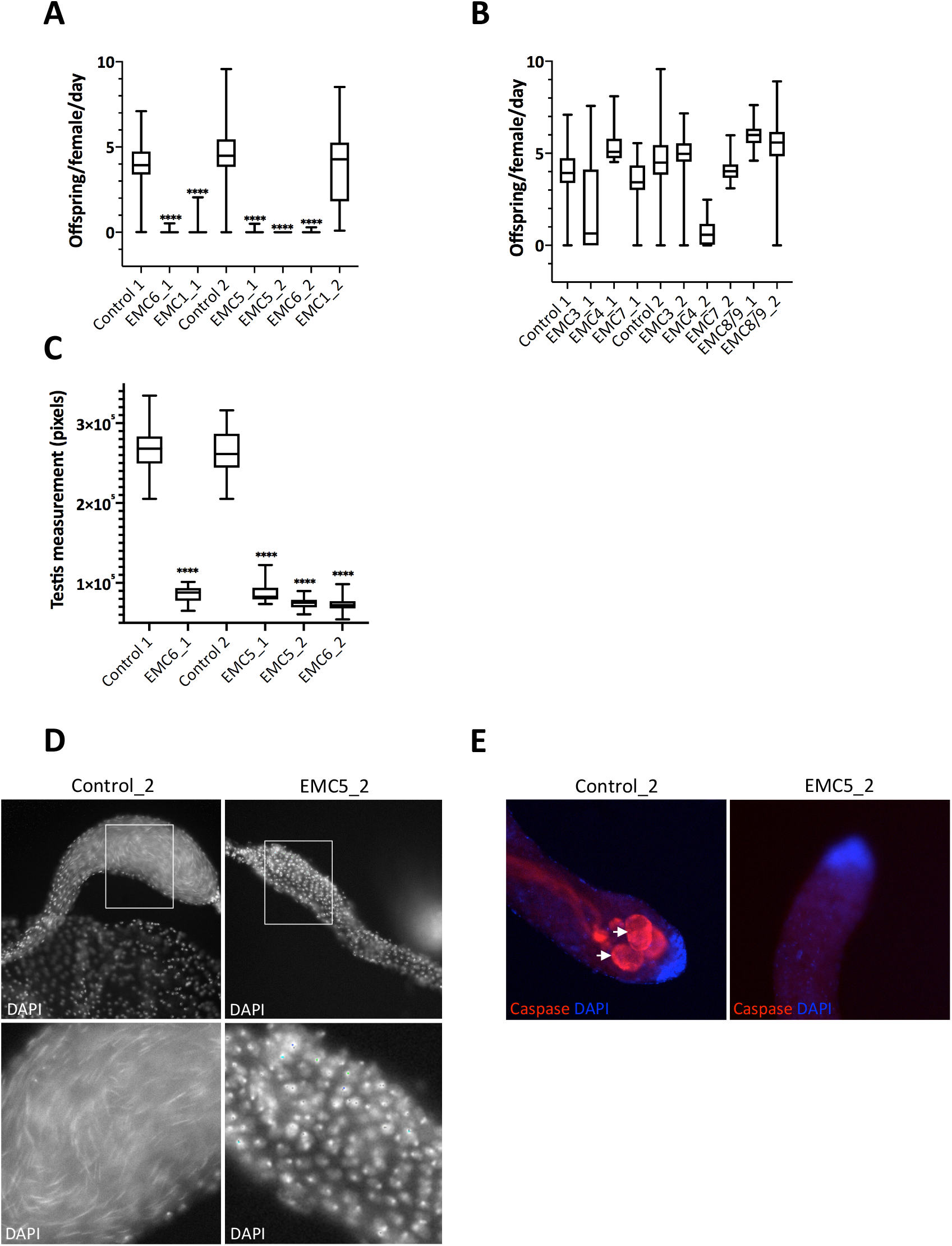
EMC is required for sperm differentiation in *Drosophila.* (A-B) Box-and-whisker plots of progeny per female per day for RNAi lines. Individual adult males of the represented RNAi lines were mated to wild-type virgin females in separate vials and the mean number of progeny for crosses of the different genotypes was counted. The mid-line in the box plots represents median progeny (per female/day), and whiskers below and above the box indicate the sample range. All tested EMC5 and EMC6 RNAi lines, and one EMC1 RNAi line (EMC1_1) have a significant reduction of progeny produced. Significance was determined by Welch’s t-test: ****p ≤ 0.0001. (C) Box-and-whisker plots of seminal vesicle size measured for 15 males per RNAi line. The box represents the 25-75^th^ percentiles, and the median is indicated. Whiskers below and above the box indicate the minimum and maximum seminal vesicle size. All tested EMC5 and EMC6 RNAi samples (Control_2) but not when lines show a significant reduction in sperm production compared to controls. Significance was determined by Welch’s t-test: ****p ≤ 0.0001. (D) Detection of sperm by nuclear staining with DAPI in seminal vesicles. Boxed areas are magnified below. The characteristic needle-like nuclei of mature sperm can be seen in control, but not when UAS-EMC5_2 RNAi is expressed (EMC5_2). (E) Immunostaining of testis with an antibody against active caspase-3 (red), showing presence of multiple cystic bulges in Control_2 (left panel) and absence of cystic bulges when UAS-EMC5_2 RNAi is expressed (right panel). Nuclear staining was performed with DAPI (blue).

**Figure S3.**
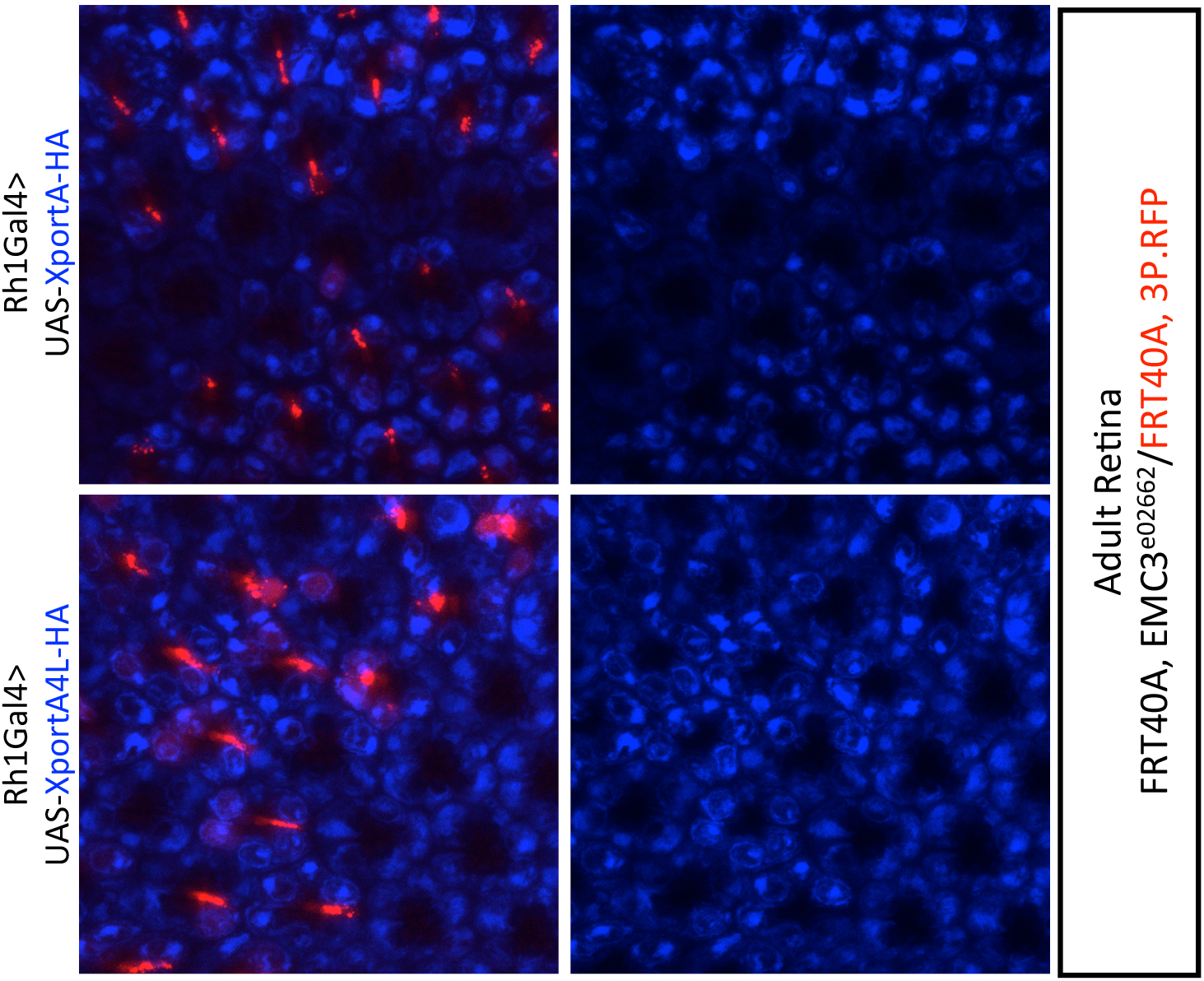
EMC3 is required for the expression of Xport-A in the adult retina, but not Xport-A4L. Immunostaining of mosaic adult retina with eyeless-Flippase-induced clones of cells homozygous for EMC3^e02662^, labelled by the absence of RFP (red), show loss of (top panel) UAS-Xport-A-HA (anti-HA, in blue) in EMC3 mutant homozygous cells. By contrast, expression of UAS- Xport-A4L-HA (bottom panel) is not lost in EMC3^e02662^ homozygous mutant clones.

**Figure S4.**
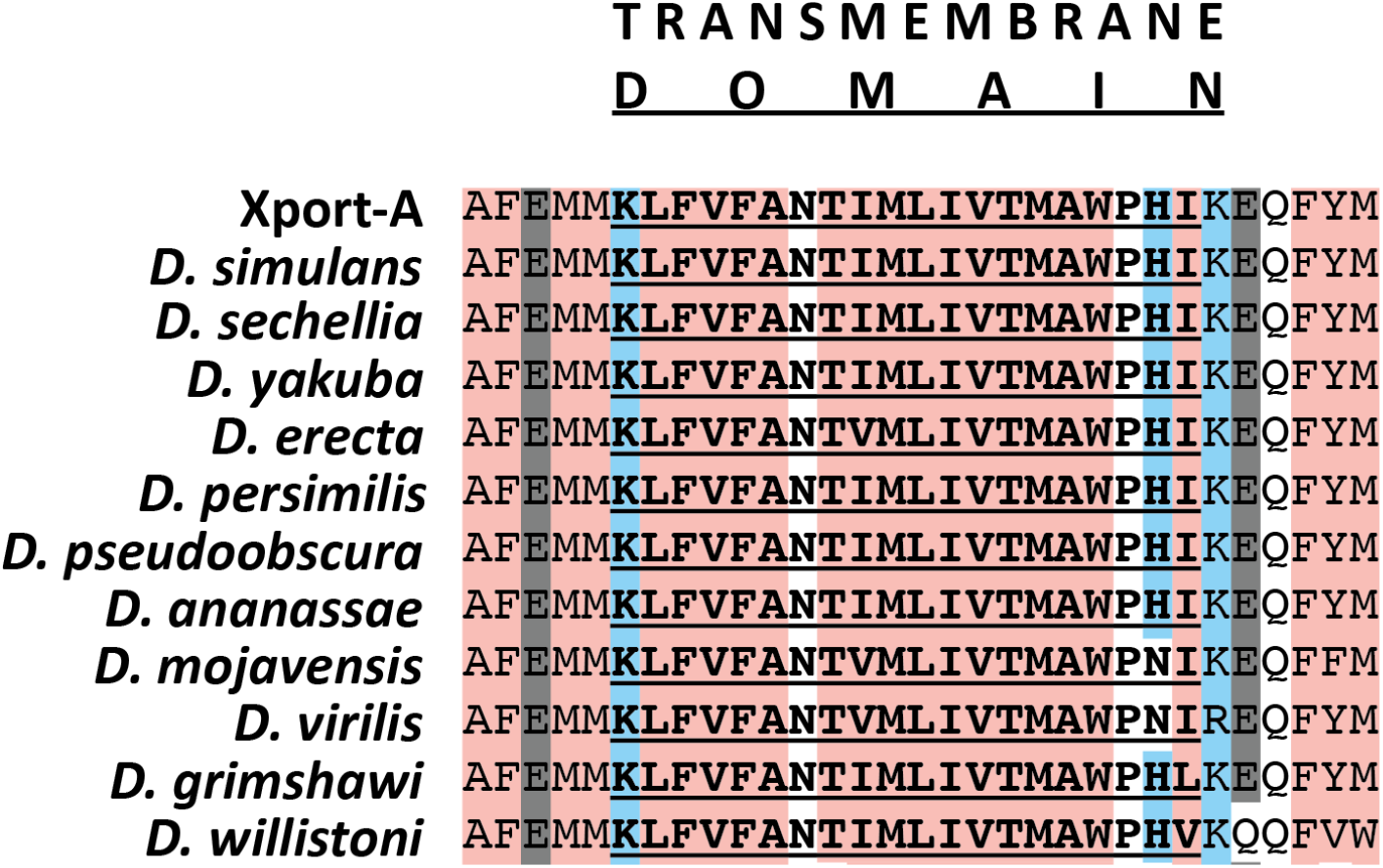
XportATMD protein alignment across *Drosophila.* Xport-A TMD sequences were aligned from 16 different *Drosophila* species, using Uniprot Blastp tool. Pink shading indicates hydrophobic residues, blue shading indicates positively charged residues, and grey shading indicates negatively charged residues. Uniprot accession numbers obtained from Blastp alignment are as follows: B4QSS5_DROSI, B4IHQ4_DROSE, B4GL95_DROYA, B3P066_DROER, B4GL95_DROPE, A0A6I8USC0_DROPS, B3LXT7_DROAN, B4KE44_DROMO, B4M4S9_DROVI, B4K0W3_DROGR, Q8I176_DROWI.

**Table S1.**
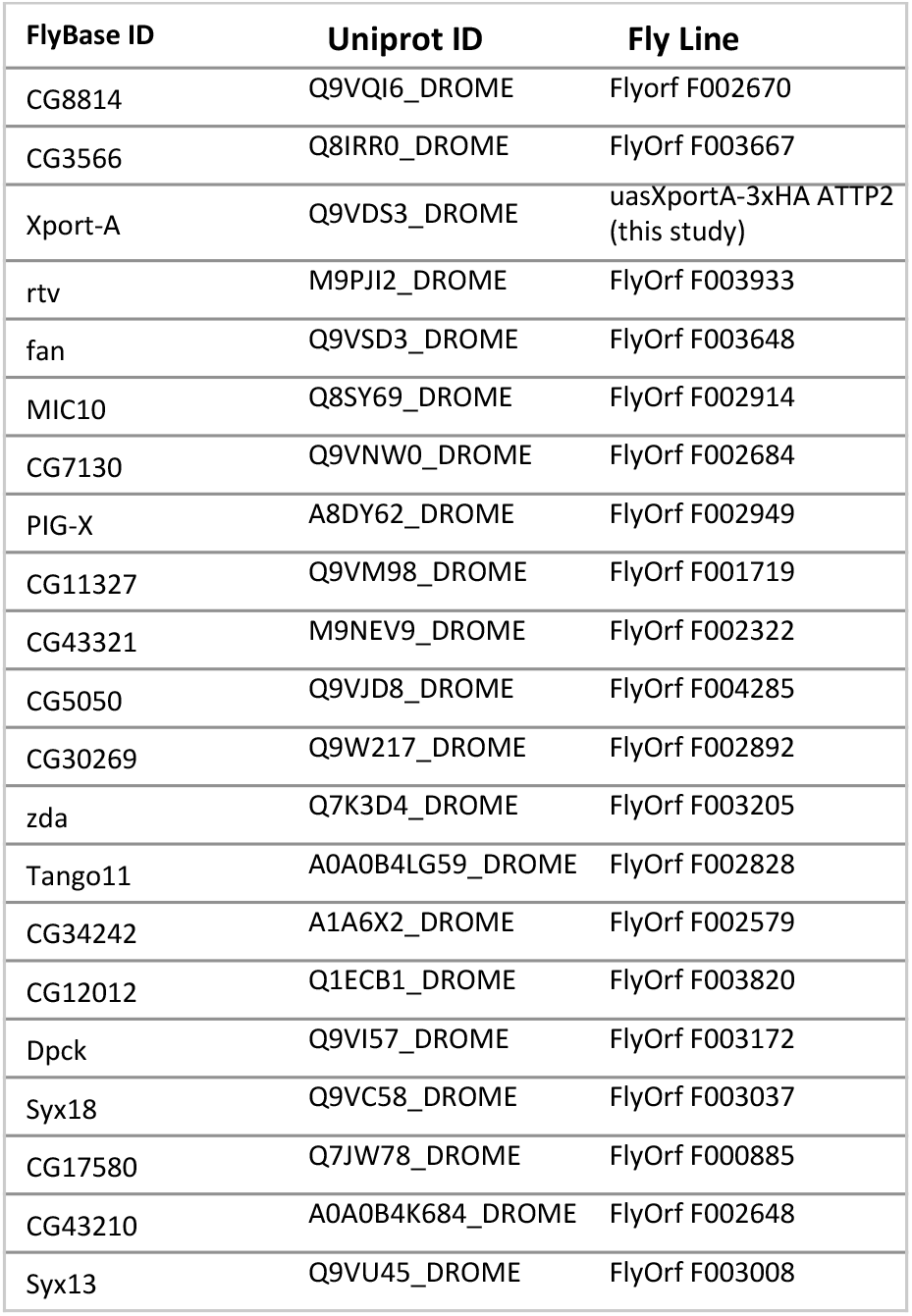
Fly Lines used to test TA proteins

**Table S2.**
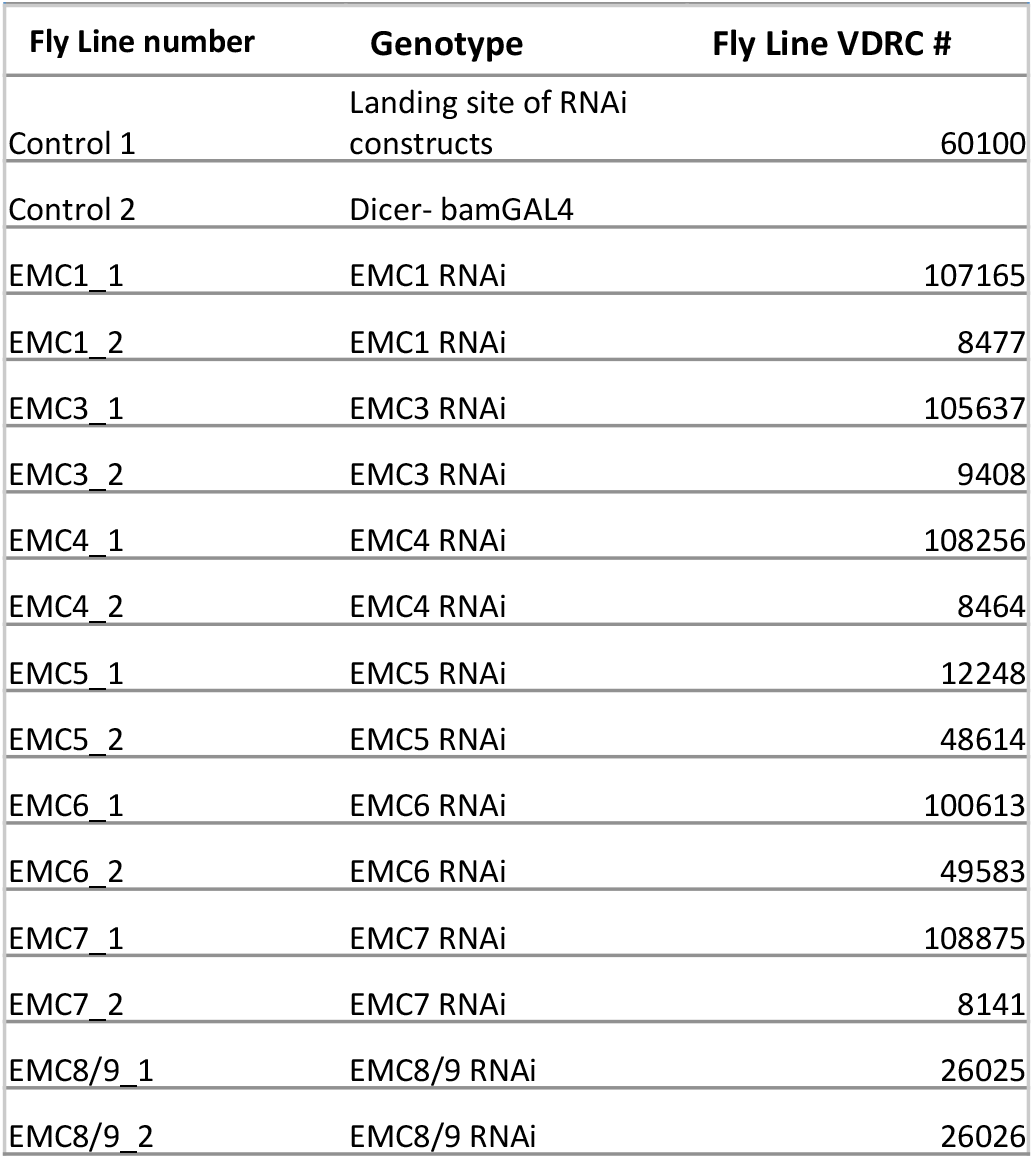
Fly Lines used for fertility experiments

## Notes

### Competing Interest Statement

The authors have declared no competing interest.

