## Supplementary file 1 for "EMC is required for biogenesis and membrane insertion of Xport-A, an essential chaperone of rhodopsin-1 and the TRP channel"

| Drosophila gene name | Uniprot ID | Most likely loc - DeepLoc | TMD HF sum | TM HF avg | tail charge | TMD seq | TMD len | tail seq | tail len | Unitprot link | Descriptions | DeepLoc all probabilities |
|---|---|---|---|---|---|---|---|---|---|---|---|---|
| | NAA60_DROME | Peroxisome, Membrane | -7,5 | -0,50 | 3 | LDHIKHYASMVRHTS | 15 | SLCAWLAGRVQQVVRWFYHKLLTRFNFI | 29 | https://www.uniprot.org/uniprot/Q95SX8 | RecName: Full=N-alpha-acetyltransferase | Peroxisome: 0.4461, Mitochondrion: 0.2328, Cytoplasm: 0.1326, Endoplasmic reticulum: 0.0744, Plastid: 0.0405, Nucleus: 0.0226, Extracellular: 0.0167, Lysosome/Vacuole: 0.0164, Cell membrane: 0.0115, Golgi apparatus: 0.0063 |
| | Q9VLC3_DROME | Cell membrane, Membrane | 5,7 | 0,36 | 1 | EPTWITLMISSMLVKL | 16 | YR | 2 | https://www.uniprot.org/uniprot/Q9VLC3 | SubName: Full=FI03373p {ECO:0000313\|E | Cell membrane: 0.736, Endoplasmic reticulum: 0.17, Lysosome/Vacuole: 0.0624, Golgi apparatus: 0.0198, Extracellular: 0.0061, Peroxisome: 0.0023, Nucleus: 0.0016, Mitochondrion: 0.0009, Cytoplasm: 0.0008, Plastid: 0.0002 |
| | X2JA11_DROME | Mitochondrion, Membrane | 6,5 | 0,31 | 0 | MRLGPGGAILLLVFEYSYDYL | 21 | LHNYS | 5 | https://www.uniprot.org/uniprot/X2JA11 | SubName: Full=Uncharacterized protein, i | Mitochondrion: 0.8719, Peroxisome: 0.0962, Endoplasmic reticulum: 0.0231, Plastid: 0.0024, Cell membrane: 0.0022, Cytoplasm: 0.0021, Lysosome/Vacuole: 0.0007, Extracellular: 0.0006, Golgi apparatus: 0.0006, Nucleus: 0.0002 |
| | Q2PDS0_DROME | Mitochondrion, Membrane | 6,5 | 0,43 | -1 | CFVLTIPYVACFTSQ | 15 | IGTLRFFTIELNQE | 14 | https://www.uniprot.org/uniprot/Q2PDS0 | SubName: Full=Uncharacterized protein, i | Mitochondrion: 0.3398, Cytoplasm: 0.2156, Endoplasmic reticulum: 0.0948, Nucleus: 0.0882, Peroxisome: 0.0813, Golgi apparatus: 0.0629, Lysosome/Vacuole: 0.043, Cell membrane: 0.0353, Extracellular: 0.0233, Plastid: 0.0158 |
| | M9PG24_DROME | Cell membrane, Membrane | 6,6 | 0,39 | 5 | MLLRCCLVPLTAAICWR | 17 | CGCRSRCRCFYPNIQHPRQDSKSKYSSTC | 29 | https://www.uniprot.org/uniprot/M9PG24 | SubName: Full=Uncharacterized protein {E | Cell membrane: 0.4888, Mitochondrion: 0.2243, Extracellular: 0.1565, Golgi apparatus: 0.0545, Nucleus: 0.0381, Cytoplasm: 0.0192, Lysosome/Vacuole: 0.0083, Plastid: 0.0052, Endoplasmic reticulum: 0.0048, Peroxisome: 0.0003 |
| | A0A0B4K7B8_DROME | Mitochondrion, Membrane | 6,7 | 0,32 | 0 | YHSWSFVKCSLAFCAGVLLVR | 21 | SLDGKLPASS | 10 | https://www.uniprot.org/uniprot/A0A0B4K7B8 | SubName: Full=Uncharacterized protein {E | Mitochondrion: 0.6952, Extracellular: 0.2053, Golgi apparatus: 0.0304, Cytoplasm: 0.0231, Endoplasmic reticulum: 0.0173, Cell membrane: 0.0107, Peroxisome: 0.0085, Lysosome/Vacuole: 0.0066, Nucleus: 0.0026, Plastid: 0.0002 |
| | Q9VRA5_DROME | Endoplasmic reticulum, Membrane | 7,5 | 0,31 | -1 | IYNDHFCWTFIKSYGLFSLAIPLA | 24 | KYFDGFQVLPTGDV | 14 | https://www.uniprot.org/uniprot/Q9VRA5 | SubName: Full=GEO12078p1 {ECO:000031 | Endoplasmic reticulum: 0.4576, Extracellular: 0.1706, Mitochondrion: 0.1515, Golgi apparatus: 0.0627, Lysosome/Vacuole: 0.0576, Cytoplasm: 0.051, Cell membrane: 0.0274, Peroxisome: 0.0108, Plastid: 0.006, Nucleus: 0.0048 |
| | A0A0B4KGW0_DROME | Mitochondrion, Membrane | 7,7 | 0,23 | 1 | FGLMTEDPKLWLKAGLGVAAATMLGFIVLKT | 34 | AGAHTR | 6 | https://www.uniprot.org/uniprot/A0A0B4KGW0 | RecName: Full=Mitochondrial Rho GTPase | Mitochondrion: 0.3584, Endoplasmic reticulum: 0.2343, Peroxisome: 0.1252, Golgi apparatus: 0.0829, Cell membrane: 0.078, Cytoplasm: 0.0655, Lysosome/Vacuole: 0.0318, Nucleus: 0.0181, Plastid: 0.0051, Extracellular: 0.0007 |
| | Q9VJX1_DROME | Cell membrane, Membrane | 7,8 | 0,33 | 1 | MLTDFASALYCSLAIMCCCTPNML | 24 | R | 1 | https://www.uniprot.org/uniprot/Q9VJX1 | SubName: Full=Uncharacterized protein {E | Cell membrane: 0.3184, Nucleus: 0.2017, Cytoplasm: 0.1863, Golgi apparatus: 0.0993, Lysosome/Vacuole: 0.0553, Extracellular: 0.0483, Mitochondrion: 0.0376, Endoplasmic reticulum: 0.035, Peroxisome: 0.0101, Plastid: 0.008 |
| | Q9VG01_DROME | Peroxisome, Membrane | 7,8 | 0,41 | 3 | YTLKYGLIGLGCYGAFRFF | 19 | RPRLTRLF | 8 | https://www.uniprot.org/uniprot/Q9VG01 | SubName: Full=RE12073p {ECO:0000313\|I | Peroxisome: 0.4273, Mitochondrion: 0.1895, Nucleus: 0.1887, Cytoplasm: 0.065, Plastid: 0.0493, Endoplasmic reticulum: 0.043, Golgi apparatus: 0.0201, Cell membrane: 0.0124, Lysosome/Vacuole: 0.0033, Extracellular: 0.0013 |
| | Q9VLF1_DROME | Mitochondrion, Membrane | 8 | 0,38 | 1 | LRVGPGGAMLLVTYEYLFEFL | 21 | KSQNI | 5 | https://www.uniprot.org/uniprot/Q9VLF1 | SubName: Full=AT23224p {ECO:0000313\|I | Mitochondrion: 0.7659, Peroxisome: 0.1758, Endoplasmic reticulum: 0.0529, Plastid: 0.002, Cell membrane: 0.0014, Cytoplasm: 0.0013, Golgi apparatus: 0.0003, Nucleus: 0.0001, Extracellular: 0.0001, Lysosome/Vacuole: 0.0001 |
| | Q8T8S5_DROME | Mitochondrion, Membrane | 8,1 | 0,39 | 0 | SFATELLRFFGNVFVDIFNAI | 21 | FN | 2 | https://www.uniprot.org/uniprot/Q8T8S5 | SubName: Full=AT30929p {ECO:0000313\|I | Mitochondrion: 0.3334, Cytoplasm: 0.1967, Plastid: 0.163, Nucleus: 0.0946, Lysosome/Vacuole: 0.0911, Endoplasmic reticulum: 0.0509, Extracellular: 0.0286, Golgi apparatus: 0.0232, Peroxisome: 0.0102, Cell membrane: 0.0083 |
| | A0A0B4KGR8_DROME | Mitochondrion, Membrane | 8,1 | 0,41 | 1 | WKRVLFLSLKCVYLMMVVKF | 20 | SQRCLEMAFKSQFAQRQTENAQVGSQS | 33 | https://www.uniprot.org/uniprot/A0A0B4KGR8 | SubName: Full=Uncharacterized protein {E | Mitochondrion: 0.7711, Extracellular: 0.1386, Golgi apparatus: 0.0191, Cell membrane: 0.0155, Cytoplasm: 0.0149, Endoplasmic reticulum: 0.012, Lysosome/Vacuole: 0.011, Plastid: 0.0081, Nucleus: 0.005, Peroxisome: 0.0047 |
| | Q9VLC9_DROME | Nucleus, Membrane | 8,1 | 0,41 | 1 | LRCSFFCCCMAFGVYMGFKM | 20 | MR | 2 | https://www.uniprot.org/uniprot/Q9VLC9 | SubName: Full=Uncharacterized protein, i | Nucleus: 0.3681, Cytoplasm: 0.152, Golgi apparatus: 0.1284, Cell membrane: 0.1277, Mitochondrion: 0.0849, Endoplasmic reticulum: 0.0738, Peroxisome: 0.0393, Lysosome/Vacuole: 0.017, Extracellular: 0.006, Plastid: 0.0028 |
| | Q9W513_DROME | Mitochondrion, Membrane | 8,3 | 0,36 | 0 | RGAVYACIGVTLYGTYILGMRYY | 23 | HYCTVIRPEKQQAELKLLDEGAHDKAKEL | 31 | https://www.uniprot.org/uniprot/Q9W513 | SubName: Full=GEO10196p1 {ECO:000031 | Mitochondrion: 0.9324, Golgi apparatus: 0.0203, Endoplasmic reticulum: 0.0164, Cytoplasm: 0.0136, Nucleus: 0.0083, Peroxisome: 0.0064, Plastid: 0.0018, Cell membrane: 0.0005, Lysosome/Vacuole: 0.0002, Extracellular: 0.0001 |
| | G4LU19_DROME | Mitochondrion, Membrane | 8,3 | 0,44 | 2 | YGPVLYVCGAAFCYAFRWY | 19 | STINTCLLNGKLQTLNARTHAH | 22 | https://www.uniprot.org/uniprot/G4LU19 | SubName: Full=MIP32906p1 {ECO:000031 | Mitochondrion: 0.7709, Extracellular: 0.1187, Cell membrane: 0.0328, Endoplasmic reticulum: 0.0165, Lysosome/Vacuole: 0.0136, Cytoplasm: 0.0135, Golgi apparatus: 0.0119, Nucleus: 0.0104, Peroxisome: 0.0087, Plastid: 0.0031 |
| | Q4V6L7_DROME | Cell membrane, Membrane | 8,6 | 0,43 | 7 | RIVGCLFVTFCCCCCFNYYV | 20 | PCKQTDRSHYCGNCGCYFGRSMKRRQP | 35 | https://www.uniprot.org/uniprot/Q4V6L7 | SubName: Full=IP04672p {ECO:0000313\|E | Cell membrane: 0.5583, Nucleus: 0.1417, Golgi apparatus: 0.1257, Cytoplasm: 0.0642, Mitochondrion: 0.0449, Endoplasmic reticulum: 0.0303, Lysosome/Vacuole: 0.0167, Extracellular: 0.0123, Peroxisome: 0.0045, Plastid: 0.0014 |
| | RL33_DROME | Mitochondrion, Membrane | 8,7 | 0,48 | 7 | ANVAKATWASLGLVYVLV | 18 | KMHRRNTKRRETKLYCKGCQQAMLHG | 26 | https://www.uniprot.org/uniprot/Q9VR93 | RecName: Full=Putative 60S ribosomal pro | Mitochondrion: 0.6047, Extracellular: 0.1801, Cytoplasm: 0.0613, Nucleus: 0.0434, Endoplasmic reticulum: 0.0299, Cell membrane: 0.0212, Plastid: 0.0201, Peroxisome: 0.0158, Lysosome/Vacuole: 0.0124, Golgi apparatus: 0.0112 |
| | Q7K5K0_DROME | Endoplasmic reticulum, Membrane | 9,1 | 0,41 | 0 | YFKYTTIILGISVFICLKCKWF | 22 | S | 1 | https://www.uniprot.org/uniprot/Q7K5K0 | SubName: Full=GH09064p {ECO:0000313\| | Endoplasmic reticulum: 0.2935, Golgi apparatus: 0.2587, Cell membrane: 0.1694, Lysosome/Vacuole: 0.0727, Mitochondrion: 0.0705, Peroxisome: 0.0461, Nucleus: 0.0333, Cytoplasm: 0.0305, Plastid: 0.0227, Extracellular: 0.0026 |
| | M9PBQ8_DROME | Mitochondrion, Membrane | 9,1 | 0,43 | 1 | GNYKLGAGAGFLGMAMFAAFA | 21 | HK | 2 | https://www.uniprot.org/uniprot/M9PBQ8 | SubName: Full=Gdap1, isoform B {ECO:00 | Mitochondrion: 0.5736, Peroxisome: 0.2544, Endoplasmic reticulum: 0.1139, Cytoplasm: 0.0383, Plastid: 0.0069, Cell membrane: 0.0059, Golgi apparatus: 0.0028, Lysosome/Vacuole: 0.0023, Nucleus: 0.0018, Extracellular: 0.0001 |
| | Q9VRJ3_DROME | Mitochondrion, Membrane | 9,1 | 0,43 | 1 | GNYKLGAGAGFLGMAMFAAFA | 21 | HK | 2 | https://www.uniprot.org/uniprot/Q9VRJ3 | SubName: Full=Gdap1, isoform A {ECO:00 | Mitochondrion: 0.5707, Peroxisome: 0.2611, Endoplasmic reticulum: 0.1116, Cytoplasm: 0.0376, Plastid: 0.0068, Cell membrane: 0.0057, Golgi apparatus: 0.0024, Lysosome/Vacuole: 0.0023, Nucleus: 0.0017, Extracellular: 0.0001 |
| | E1JH12_DROME | Mitochondrion, Membrane | 9,1 | 0,46 | -3 | RYAQIFIVMAVDLLTLSNLY | 20 | PKHSSYIARPFLLWQDLQEEDERSLDSPQ | 31 | https://www.uniprot.org/uniprot/E1JH12 | SubName: Full=Uncharacterized protein {E | Mitochondrion: 0.36, Endoplasmic reticulum: 0.3176, Extracellular: 0.1071, Cytoplasm: 0.0951, Golgi apparatus: 0.0518, Nucleus: 0.0361, Lysosome/Vacuole: 0.0225, Cell membrane: 0.0062, Peroxisome: 0.0024, Plastid: 0.0011 |
| | E2QD56_DROME | Golgi apparatus, Membrane | 9,2 | 0,42 | 0 | WDSVCCGVLLCCWHYCTVLTVI | 22 | P | 1 | https://www.uniprot.org/uniprot/E2QD56 | SubName: Full=Exchange factor for Arf 6, i | Golgi apparatus: 0.2877, Nucleus: 0.2673, Cell membrane: 0.2474, Cytoplasm: 0.1415, Endoplasmic reticulum: 0.0271, Lysosome/Vacuole: 0.0259, Mitochondrion: 0.0018, Plastid: 0.0005, Peroxisome: 0.0005, Extracellular: 0.0004 |
| | A0A1W5PX42_DROME | Mitochondrion, Membrane | 9,2 | 0,46 | 3 | LQYGVHLTTTLISAIGNFLY | 20 | RLWSPPLRRSC | 11 | https://www.uniprot.org/uniprot/A0A1W5PX42 | SubName: Full=Uncharacterized protein {E | Mitochondrion: 0.8251, Endoplasmic reticulum: 0.108, Peroxisome: 0.0233, Extracellular: 0.0136, Plastid: 0.0114, Cell membrane: 0.0079, Golgi apparatus: 0.0041, Lysosome/Vacuole: 0.0034, Cytoplasm: 0.0026, Nucleus: 0.0006 |
| | Q6IGW6_DROME | Mitochondrion, Membrane | 9,2 | 0,51 | -1 | QWMLVKSYGLFFLGVRLA | 18 | KEFVGVELMPSLGPA | 15 | https://www.uniprot.org/uniprot/Q6IGW6 | SubName: Full=GEO13367p1 {ECO:000031 | Mitochondrion: 0.7989, Cytoplasm: 0.077, Extracellular: 0.053, Peroxisome: 0.0278, Endoplasmic reticulum: 0.015, Golgi apparatus: 0.0136, Nucleus: 0.0054, Lysosome/Vacuole: 0.0042, Cell membrane: 0.0033, Plastid: 0.0017 |
| | A0A0B4KGL6_DROME | Endoplasmic reticulum, Membrane | 9,4 | 0,41 | 0 | KIVAGVGAVLAMGAFAAWRGIYN | 23 | QFWFNVFCWLQTLL | 14 | https://www.uniprot.org/uniprot/A0A0B4KGL6 | SubName: Full=Uncharacterized protein, i | Endoplasmic reticulum: 0.4312, Mitochondrion: 0.3024, Peroxisome: 0.2041, Plastid: 0.0177, Cell membrane: 0.0144, Cytoplasm: 0.0115, Golgi apparatus: 0.0089, Lysosome/Vacuole: 0.0048, Extracellular: 0.0038, Nucleus: 0.0011 |
| | Q9W219_DROME | Cell membrane, Membrane | 10 | 0,45 | 1 | TYCMAALLCLFLCWPCVCAPCC | 22 | CNCCYKTSQYCPNCNSCLGSF | 21 | https://www.uniprot.org/uniprot/Q9W219 | SubName: Full=GEO12575p1 {ECO:000031 | Cell membrane: 0.8007, Extracellular: 0.1211, Nucleus: 0.0438, Golgi apparatus: 0.0193, Cytoplasm: 0.0062, Lysosome/Vacuole: 0.0062, Endoplasmic reticulum: 0.0018, Mitochondrion: 0.0009, Plastid: 0.0, Peroxisome: 0.0 |
| | E2QCN8_DROME | Nucleus, Membrane | 10,4 | 0,45 | 1 | IPQPIIAATTIITIIITTTADMI | 23 | RL | 2 | https://www.uniprot.org/uniprot/E2QCN8 | SubName: Full=Uncharacterized protein {E | Nucleus: 0.393, Endoplasmic reticulum: 0.168, Mitochondrion: 0.1142, Cytoplasm: 0.1136, Plastid: 0.0848, Lysosome/Vacuole: 0.0617, Golgi apparatus: 0.0431, Peroxisome: 0.0148, Cell membrane: 0.0065, Extracellular: 0.0004 |
| | A8DZ28_DROME | Mitochondrion, Membrane | 10,4 | 0,50 | 1 | ITIISTKVLLYISSLMEHMYL | 21 | RER | 3 | https://www.uniprot.org/uniprot/A8DZ28 | SubName: Full=Defective proboscis extens | Mitochondrion: 0.4348, Peroxisome: 0.3823, Cytoplasm: 0.0664, Endoplasmic reticulum: 0.0557, Nucleus: 0.0263, Cell membrane: 0.0129, Lysosome/Vacuole: 0.0067, Extracellular: 0.006, Plastid: 0.0052, Golgi apparatus: 0.0035 |
| CG8814 | Q9VQI6_DROME | Endoplasmic reticulum, Membrane | 10,8 | 0,54 | 2 | SPLWFAVLILWPFLVRRFAR | 20 | MLQSNPQRRH | 10 | https://www.uniprot.org/uniprot/Q9VQI6 | SubName: Full=LD25952p {ECO:0000313\|I | Endoplasmic reticulum: 0.6496, Peroxisome: 0.1298, Mitochondrion: 0.0919, Golgi apparatus: 0.0826, Cell membrane: 0.0254, Cytoplasm: 0.0072, Lysosome/Vacuole: 0.0053, Nucleus: 0.0046, Plastid: 0.0035, Extracellular: 0.0001 |
| | Q9W1B2_DROME | Mitochondrion, Membrane | 11 | 0,55 | 1 | FCLKGGSGLIIGSAVTLFFT | 20 | RPQTYPIWLGLGVGMGVAYDCCQARWI | 29 | https://www.uniprot.org/uniprot/Q9W1B2 | RecName: Full=MICOS complex subunit M | Mitochondrion: 0.8758, Cell membrane: 0.0438, Extracellular: 0.0414, Endoplasmic reticulum: 0.0172, Plastid: 0.0078, Golgi apparatus: 0.0068, Cytoplasm: 0.0029, Lysosome/Vacuole: 0.0029, Peroxisome: 0.001, Nucleus: 0.0004 |
| | E1JJ97_DROME | Nucleus, Membrane | 11,2 | 0,59 | 0 | LFVRLLCLILDLLQAKLYL | 19 | QYT | 3 | https://www.uniprot.org/uniprot/E1JJ97 | SubName: Full=Uncharacterized protein {E | Nucleus: 0.4549, Endoplasmic reticulum: 0.2295, Cytoplasm: 0.1741, Peroxisome: 0.0543, Mitochondrion: 0.0391, Golgi apparatus: 0.0342, Cell membrane: 0.0085, Lysosome/Vacuole: 0.0038, Extracellular: 0.0014, Plastid: 0.0002 |
| | A0A0C4DHC8_DROME | Mitochondrion, Membrane | 11,2 | 0,62 | 2 | HLSHVLAIGLGVGLTVLT | 18 | NYYLQGKLLSK | 11 | https://www.uniprot.org/uniprot/A0A0C4DHC8 | SubName: Full=Aldehyde dehydrogenase t | Mitochondrion: 0.5257, Peroxisome: 0.2773, Endoplasmic reticulum: 0.1302, Cell membrane: 0.0231, Plastid: 0.0147, Cytoplasm: 0.0116, Golgi apparatus: 0.0064, Nucleus: 0.006, Lysosome/Vacuole: 0.0047, Extracellular: 0.0003 |
| | M9PGE1_DROME | Nucleus, Membrane | 11,2 | 0,62 | 0 | TFLMGLSQACLPGLLLSL | 18 | L | 1 | https://www.uniprot.org/uniprot/M9PGE1 | SubName: Full=Dpr-interacting protein alp | Nucleus: 0.2759, Cell membrane: 0.2504, Endoplasmic reticulum: 0.195, Golgi apparatus: 0.1402, Cytoplasm: 0.0509, Peroxisome: 0.0482, Lysosome/Vacuole: 0.0229, Mitochondrion: 0.0127, Extracellular: 0.0031, Plastid: 0.0007 |
| | M9PF87_DROME | Peroxisome, Membrane | 11,2 | 0,66 | 1 | AVLLPLVVLAALSRTSI | 17 | FRGF | 4 | https://www.uniprot.org/uniprot/M9PF87 | SubName: Full=Carbonic anhydrase 2, isof | Peroxisome: 0.5072, Endoplasmic reticulum: 0.133, Mitochondrion: 0.1324, Cytoplasm: 0.1252, Plastid: 0.0319, Lysosome/Vacuole: 0.0279, Cell membrane: 0.0251, Nucleus: 0.0064, Golgi apparatus: 0.0058, Extracellular: 0.0051 |
| | A8DZ27_DROME | Endoplasmic reticulum, Membrane | 11,7 | 0,49 | 0 | RIRAFTAMITIISTKVLLYISSLM | 24 | EHMYLRER | 8 | https://www.uniprot.org/uniprot/A8DZ27 | SubName: Full=Defective proboscis extens | Endoplasmic reticulum: 0.4131, Golgi apparatus: 0.1831, Cell membrane: 0.1523, Mitochondrion: 0.132, Peroxisome: 0.082, Lysosome/Vacuole: 0.0297, Cytoplasm: 0.003, Nucleus: 0.0027, Extracellular: 0.0018, Plastid: 0.0002 |
| | Q2PE16_DROME | Cell membrane, Membrane | 11,7 | 0,56 | 5 | HMVAGLLCLVGFCCCACFVPY | 21 | CMNSCRTGNHYCRKCNTFLGAYNPKGM | 29 | https://www.uniprot.org/uniprot/Q2PE16 | SubName: Full=FI16615p1 {ECO:0000313\| | Cell membrane: 0.7061, Golgi apparatus: 0.1848, Nucleus: 0.0496, Cytoplasm: 0.0255, Mitochondrion: 0.0137, Endoplasmic reticulum: 0.0133, Extracellular: 0.0049, Lysosome/Vacuole: 0.0011, Peroxisome: 0.0007, Plastid: 0.0002 |
| | Q6IDF5_DROME | Mitochondrion, Membrane | 11,7 | 0,59 | 2 | FRTGLFAVVLPIALYAWALK | 20 | AERDGREEKYRTGQVAYKDRQFKFI | 25 | https://www.uniprot.org/uniprot/Q6IDF5 | SubName: Full=NADH dehydrogenase (Ubi | Mitochondrion: 0.7042, Peroxisome: 0.1504, Endoplasmic reticulum: 0.0679, Cytoplasm: 0.0573, Nucleus: 0.0114, Plastid: 0.0048, Golgi apparatus: 0.0019, Cell membrane: 0.0013, Extracellular: 0.0005, Lysosome/Vacuole: 0.0003 |
| | Q9VW25_DROME | Nucleus, Membrane | 11,8 | 0,59 | 1 | PLLLTAKCVFVASLISTVYL | 20 | KLY | 3 | https://www.uniprot.org/uniprot/Q9VW25 | SubName: Full=FI04505p {ECO:0000313\|E | Nucleus: 0.503, Cell membrane: 0.1899, Endoplasmic reticulum: 0.1252, Mitochondrion: 0.0532, Peroxisome: 0.0459, Cytoplasm: 0.0389, Golgi apparatus: 0.0286, Lysosome/Vacuole: 0.009, Plastid: 0.0055, Extracellular: 0.0008 |
| | A0A0B4K8A8_DROME | Cell membrane, Membrane | 11,9 | 0,54 | 1 | HLLALLICMLGGICCCCIPYCT | 22 | DSCQSAKHTCTSCGAYVGTYKN | 22 | https://www.uniprot.org/uniprot/A0A0B4K8A8 | SubName: Full=Uncharacterized protein, i | Cell membrane: 0.7584, Golgi apparatus: 0.0782, Nucleus: 0.0514, Endoplasmic reticulum: 0.0423, Extracellular: 0.0347, Cytoplasm: 0.0167, Mitochondrion: 0.0155, Lysosome/Vacuole: 0.0018, Peroxisome: 0.0007, Plastid: 0.0003 |
| | Q9W218_DROME | Cell membrane, Membrane | 11,9 | 0,54 | 1 | HLLALLICMLGGICCCCIPYCT | 22 | DSCQSAKHTCTSCGAYVGTYKN | 22 | https://www.uniprot.org/uniprot/Q9W218 | SubName: Full=GEO08374p1 {ECO:000031 | Cell membrane: 0.7547, Golgi apparatus: 0.0784, Extracellular: 0.053, Endoplasmic reticulum: 0.0521, Nucleus: 0.0343, Mitochondrion: 0.0131, Cytoplasm: 0.0128, Lysosome/Vacuole: 0.0011, Peroxisome: 0.0004, Plastid: 0.0002 |
| | A1Z6Z4_DROME | Mitochondrion, Membrane | 12 | 0,60 | 2 | LSHVLAIGLGVGLTVLTNYY | 20 | LQGKLLSK | 8 | https://www.uniprot.org/uniprot/A1Z6Z4 | SubName: Full=Aldehyde dehydrogenase t | Mitochondrion: 0.5217, Peroxisome: 0.2762, Endoplasmic reticulum: 0.1344, Cell membrane: 0.0238, Plastid: 0.0142, Cytoplasm: 0.0117, Golgi apparatus: 0.0067, Nucleus: 0.0062, Lysosome/Vacuole: 0.0048, Extracellular: 0.0003 |
| CG3566 | Q8IRR0_DROME | Mitochondrion, Membrane | 12,1 | 0,55 | 1 | HVALALGAAFIGISLVYVIRRG | 22 | VARN | 4 | https://www.uniprot.org/uniprot/Q8IRR0 | SubName: Full=Uncharacterized protein, i | Mitochondrion: 0.4838, Peroxisome: 0.3699, Endoplasmic reticulum: 0.1214, Cytoplasm: 0.0122, Plastid: 0.0055, Cell membrane: 0.005, Nucleus: 0.0009, Golgi apparatus: 0.0007, Lysosome/Vacuole: 0.0005, Extracellular: 0.0002 |
| | Q59DV8_DROME | Cell membrane, Membrane | 12,1 | 0,64 | -1 | LLYAAMLALAYLSSWTTLR | 19 | MSDVAT | 6 | https://www.uniprot.org/uniprot/Q59DV8 | SubName: Full=Glial cell line-derived neur | Cell membrane: 0.5369, Extracellular: 0.2842, Nucleus: 0.0603, Endoplasmic reticulum: 0.0508, Cytoplasm: 0.032, Golgi apparatus: 0.0258, Mitochondrion: 0.0054, Lysosome/Vacuole: 0.0028, Peroxisome: 0.0014, Plastid: 0.0003 |
| | Q0E8X7_DROME | Mitochondrion, Membrane | 12,2 | 0,61 | 0 | WIWRYYFIAAVACVPVFYKI | 20 | SKLANSPENKKAWAESQAKEHAEHH | 25 | https://www.uniprot.org/uniprot/Q0E8X7 | SubName: Full=Reduction of Rh1, isoform | Mitochondrion: 0.7642, Endoplasmic reticulum: 0.168, Golgi apparatus: 0.0252, Peroxisome: 0.0151, Cytoplasm: 0.0087, Extracellular: 0.0081, Cell membrane: 0.0077, Nucleus: 0.002, Plastid: 0.0006, Lysosome/Vacuole: 0.0004 |
| | A0A0C4FEI5_DROME | Mitochondrion, Membrane | 12,4 | 0,52 | 0 | LHLSHVLAIGLGVGLTVLTNYYLQ | 24 | KSSTD | 5 | https://www.uniprot.org/uniprot/A0A0C4FEI5 | SubName: Full=Aldehyde dehydrogenase t | Mitochondrion: 0.466, Peroxisome: 0.244, Endoplasmic reticulum: 0.0985, Cytoplasm: 0.0619, Plastid: 0.0442, Nucleus: 0.0266, Cell membrane: 0.0233, Golgi apparatus: 0.0189, Lysosome/Vacuole: 0.0147, Extracellular: 0.002 |
| | A4UZ69_DROME | Mitochondrion, Membrane | 12,4 | 0,56 | 2 | HLSHVLAIGLGVGLTVLTNYYL | 22 | QGKLLSK | 7 | https://www.uniprot.org/uniprot/A4UZ69 | SubName: Full=Aldehyde dehydrogenase t | Mitochondrion: 0.5264, Peroxisome: 0.2711, Endoplasmic reticulum: 0.1335, Cell membrane: 0.0231, Plastid: 0.0154, Cytoplasm: 0.013, Golgi apparatus: 0.0064, Nucleus: 0.0055, Lysosome/Vacuole: 0.0053, Extracellular: 0.0003 |
| | Q4QQ69_DROME | Nucleus, Membrane | 12,5 | 0,66 | 0 | AGLLLHRALLFGSGIYLTL | 19 | L | 1 | https://www.uniprot.org/uniprot/Q4QQ69 | SubName: Full=Dpr-interacting protein de | Nucleus: 0.2263, Endoplasmic reticulum: 0.2099, Mitochondrion: 0.1832, Peroxisome: 0.1442, Golgi apparatus: 0.1281, Cell membrane: 0.0572, Cytoplasm: 0.031, Lysosome/Vacuole: 0.015, Plastid: 0.0038, Extracellular: 0.0013 |
| | M9NDW2_DROME | Endoplasmic reticulum, Membrane | 12,5 | 0,69 | 1 | GLGWFCGLNLNATLVYLI | 18 | RCIL | 4 | https://www.uniprot.org/uniprot/M9NDW2 | SubName: Full=AMP deaminase, isoform I | Endoplasmic reticulum: 0.4103, Peroxisome: 0.169, Nucleus: 0.1421, Cytoplasm: 0.0788, Mitochondrion: 0.0671, Golgi apparatus: 0.0656, Cell membrane: 0.038, Lysosome/Vacuole: 0.0144, Plastid: 0.0138, Extracellular: 0.0011 |
| | Q8MRM7_DROME | Mitochondrion, Membrane | 12,7 | 0,60 | 1 | LFDYFHLAAISVGIHASLVAL | 21 | HQPYIK | 6 | https://www.uniprot.org/uniprot/Q8MRM7 | SubName: Full=GH14903p {ECO:0000313\| | Mitochondrion: 0.4142, Peroxisome: 0.2438, Nucleus: 0.1175, Endoplasmic reticulum: 0.0892, Cytoplasm: 0.0849, Cell membrane: 0.0174, Golgi apparatus: 0.0113, Plastid: 0.0093, Lysosome/Vacuole: 0.0064, Extracellular: 0.0059 |
| | X2JCS2_DROME | Endoplasmic reticulum, Membrane | 12,8 | 0,64 | 3 | IAGDYVLCVSMTIIWMKMLL | 20 | IKTNLPFRDLAMYLKVLLKGQI | 22 | https://www.uniprot.org/uniprot/X2JCS2 | SubName: Full=Crk oncogene, isoform G {I | Endoplasmic reticulum: 0.4016, Mitochondrion: 0.1699, Peroxisome: 0.1695, Cytoplasm: 0.076, Lysosome/Vacuole: 0.0509, Extracellular: 0.0423, Golgi apparatus: 0.0378, Cell membrane: 0.0336, Nucleus: 0.0135, Plastid: 0.0049 |
| | F0JAK5_DROME | Cell membrane, Membrane | 12,8 | 0,67 | 2 | WLGLAILSLFVPCLWCYPP | 19 | LRACHLAGIHCGLCGGQHKPHV | 22 | https://www.uniprot.org/uniprot/F0JAK5 | SubName: Full=LD31195p {ECO:0000313\|I | Cell membrane: 0.6227, Nucleus: 0.2078, Golgi apparatus: 0.0462, Cytoplasm: 0.0419, Mitochondrion: 0.0339, Endoplasmic reticulum: 0.0238, Peroxisome: 0.0136, Extracellular: 0.0062, Lysosome/Vacuole: 0.0027, Plastid: 0.0012 |
| | Q7K0G4_DROME | Cell membrane, Membrane | 12,8 | 0,67 | 2 | WLGLAILSLFVPCLWCYPP | 19 | LRACHLAGIHCGLCGGQHKPHV | 22 | https://www.uniprot.org/uniprot/Q7K0G4 | SubName: Full=SD10629p {ECO:0000313\|I | Cell membrane: 0.645, Nucleus: 0.1895, Golgi apparatus: 0.0433, Cytoplasm: 0.0416, Mitochondrion: 0.0322, Endoplasmic reticulum: 0.0219, Peroxisome: 0.0148, Extracellular: 0.008, Lysosome/Vacuole: 0.0027, Plastid: 0.0011 |
| | MTX1_DROME | Mitochondrion, Membrane | 12,9 | 0,59 | -9 | IVAGVGAVLAMGAFAAWRGIYN | 22 | QLTTRSSTDYDGIDYEDDEMEEGLD | 25 | https://www.uniprot.org/uniprot/Q9VHB6 | RecName: Full=Metaxin-1 homolog; | Mitochondrion: 0.3524, Peroxisome: 0.3522, Cytoplasm: 0.1188, Plastid: 0.0735, Endoplasmic reticulum: 0.0495, Extracellular: 0.0175, Lysosome/Vacuole: 0.0116, Cell membrane: 0.0114, Nucleus: 0.0111, Golgi apparatus: 0.0019 |
| Xport-A | Q9VDS3_DROME | Endoplasmic reticulum, Membrane | 12,9 | 0,65 | -2 | KLFVFANTIMLIVTMAWPHI | 20 | KEQFYMVNQWLESFREEHQQ | 20 | https://www.uniprot.org/uniprot/Q9VDS3 | SubName: Full=Exit protein of rhodopsin a | Endoplasmic reticulum: 0.4245, Nucleus: 0.3421, Mitochondrion: 0.1323, Peroxisome: 0.0303, Cytoplasm: 0.029, Golgi apparatus: 0.0216, Cell membrane: 0.0114, Extracellular: 0.0059, Lysosome/Vacuole: 0.0021, Plastid: 0.0007 |
| | D0IQC0_DROME | Mitochondrion, Membrane | 13 | 0,59 | 1 | YGPAVRIGIAVLVLHTVGWLGW | 22 | KAVNQGAESKEAAAKAQQEQLRAVFAEI | 28 | https://www.uniprot.org/uniprot/D0IQC0 | SubName: Full=MIP12466p {ECO:0000313 | Mitochondrion: 0.8121, Cytoplasm: 0.041, Endoplasmic reticulum: 0.0375, Golgi apparatus: 0.0322, Plastid: 0.0298, Peroxisome: 0.0238, Extracellular: 0.0067, Cell membrane: 0.0067, Lysosome/Vacuole: 0.0051, Nucleus: 0.005 |
| | Q9W2E8_DROME | Mitochondrion, Membrane | 13 | 0,59 | -1 | LFRGLGVGFCAFLATVAVEYAL | 22 | GIGKGQGGHGHGHGHEEHGDKGHH | 24 | https://www.uniprot.org/uniprot/Q9W2E8 | SubName: Full=NADH dehydrogenase (Ubi | Mitochondrion: 0.7733, Endoplasmic reticulum: 0.0589, Golgi apparatus: 0.0532, Cell membrane: 0.0475, Nucleus: 0.0322, Extracellular: 0.0127, Lysosome/Vacuole: 0.0086, Plastid: 0.0058, Cytoplasm: 0.0056, Peroxisome: 0.0022 |
| | Q9VIM9_DROME | Peroxisome, Membrane | 13,1 | 0,52 | 7 | LDIYALIGAISIGLLLMLKKVVQIT | 25 | CRRLRTKLNKVKKQ | 14 | https://www.uniprot.org/uniprot/Q9VIM9 | SubName: Full=UDP-glycosyltransferase 3 | Peroxisome: 0.5424, Mitochondrion: 0.1661, Endoplasmic reticulum: 0.1425, Plastid: 0.0552, Cytoplasm: 0.0377, Cell membrane: 0.0268, Lysosome/Vacuole: 0.0142, Golgi apparatus: 0.0078, Nucleus: 0.0047, Extracellular: 0.0027 |
| | A0A0B4K6C3_DROME | Mitochondrion, Membrane | 13,1 | 0,60 | 0 | VLYRVTVALALVGIGGMGKLFY | 22 | ELSVPKKE | 8 | https://www.uniprot.org/uniprot/A0A0B4K6C3 | SubName: Full=Cytochrome c oxidase sub | Mitochondrion: 0.6733, Peroxisome: 0.1779, Plastid: 0.0714, Endoplasmic reticulum: 0.052, Cytoplasm: 0.0209, Nucleus: 0.0019, Cell membrane: 0.0012, Lysosome/Vacuole: 0.0009, Extracellular: 0.0004, Golgi apparatus: 0.0002 |
| | A0A0B4K6E6_DROME | Endoplasmic reticulum, Membrane | 13,1 | 0,62 | -1 | GSRLILTLAMSLTSIWGHFLM | 21 | SLCDITVNAV | 10 | https://www.uniprot.org/uniprot/A0A0B4K6E6 | SubName: Full=Uncharacterized protein, i | Endoplasmic reticulum: 0.2994, Nucleus: 0.19, Mitochondrion: 0.1709, Cytoplasm: 0.1148, Peroxisome: 0.0878, Golgi apparatus: 0.0789, Lysosome/Vacuole: 0.0256, Cell membrane: 0.0229, Extracellular: 0.0078, Plastid: 0.0019 |
| | M9PJG3_DROME | Cell membrane, Membrane | 13,2 | 0,55 | 4 | TNLWALCLCTFGWCCCACLFPYLW | 24 | NGCRTTNHYCSACEIFLGAHYPRSCCCCC | 33 | https://www.uniprot.org/uniprot/M9PJG3 | SubName: Full=Uncharacterized protein, i | Cell membrane: 0.7666, Golgi apparatus: 0.1249, Nucleus: 0.0731, Lysosome/Vacuole: 0.0123, Endoplasmic reticulum: 0.0087, Mitochondrion: 0.0071, Cytoplasm: 0.0047, Extracellular: 0.0022, Peroxisome: 0.0003, Plastid: 0.0001 |
| | Q9W220_DROME | Cell membrane, Membrane | 13,2 | 0,57 | 1 | THMCALTLYICCCWPFICLPYFI | 23 | NYCKSVQHYCPNCGCHIGSYSI | 22 | https://www.uniprot.org/uniprot/Q9W220 | SubName: Full=Uncharacterized protein, i | Cell membrane: 0.6977, Extracellular: 0.1636, Cytoplasm: 0.0413, Golgi apparatus: 0.0412, Nucleus: 0.0286, Mitochondrion: 0.011, Lysosome/Vacuole: 0.0106, Endoplasmic reticulum: 0.0058, Plastid: 0.0002, Peroxisome: 0.0001 |
| | Q8IRX1_DROME | Endoplasmic reticulum, Membrane | 13,3 | 0,67 | 0 | ACYWIGLAHLLMIISCMRGL | 20 | F | 1 | https://www.uniprot.org/uniprot/Q8IRX1 | SubName: Full=IP15948p {ECO:0000313\|E | Endoplasmic reticulum: 0.2942, Cell membrane: 0.2576, Mitochondrion: 0.2239, Golgi apparatus: 0.0971, Peroxisome: 0.0667, Nucleus: 0.0342, Cytoplasm: 0.0141, Lysosome/Vacuole: 0.0103, Plastid: 0.001, Extracellular: 0.0008 |
| rtv | M9PJI2_DROME | Extracellular, Membrane | 13,3 | 0,70 | 2 | SAPQMILITMLPLLGSWLL | 19 | QRMRN | 5 | https://www.uniprot.org/uniprot/M9PJI2 | RecName: Full=Protein quiver {ECO:00002 | Extracellular: 0.2494, Cell membrane: 0.2175, Endoplasmic reticulum: 0.1942, Mitochondrion: 0.192, Lysosome/Vacuole: 0.0569, Golgi apparatus: 0.0476, Peroxisome: 0.0292, Cytoplasm: 0.007, Nucleus: 0.0055, Plastid: 0.0006 |
| | E2QCK0_DROME | Endoplasmic reticulum, Membrane | 13,3 | 0,70 | 1 | KIVYCLGLAYFVLKTIFWL | 19 | NRN | 3 | https://www.uniprot.org/uniprot/E2QCK0 | SubName: Full=Transport and golgi organi | Endoplasmic reticulum: 0.388, Nucleus: 0.1543, Peroxisome: 0.1332, Golgi apparatus: 0.1054, Mitochondrion: 0.0889, Cytoplasm: 0.0411, Lysosome/Vacuole: 0.0354, Cell membrane: 0.0281, Extracellular: 0.0158, Plastid: 0.0099 |
| | TNG11_DROME | Endoplasmic reticulum, Membrane | 13,3 | 0,70 | 1 | KIVYCLGLAYFVLKTIFWL | 19 | NRN | 3 | https://www.uniprot.org/uniprot/Q961C9 | RecName: Full=Transport and Golgi organi | Endoplasmic reticulum: 0.388, Nucleus: 0.1543, Peroxisome: 0.1332, Golgi apparatus: 0.1054, Mitochondrion: 0.0889, Cytoplasm: 0.0411, Lysosome/Vacuole: 0.0354, Cell membrane: 0.0281, Extracellular: 0.0158, Plastid: 0.0099 |
| | Q8MQP2_DROME | Endoplasmic reticulum, Membrane | 13,4 | 0,89 | 1 | FSFMSLCFWALSSWM | 15 | QRL | 3 | https://www.uniprot.org/uniprot/Q8MQP2 | SubName: Full=MIP16184p {ECO:0000313 | Endoplasmic reticulum: 0.4196, Cell membrane: 0.1811, Lysosome/Vacuole: 0.1003, Mitochondrion: 0.0779, Cytoplasm: 0.0592, Peroxisome: 0.0555, Golgi apparatus: 0.0497, Extracellular: 0.0271, Plastid: 0.0235, Nucleus: 0.0062 |
| | A8JUP5_DROME | Cell membrane, Membrane | 13,5 | 0,59 | 4 | NLWALCLCTFGWCCCACLFPYLW | 23 | NGCRTTNHYCSACEIFLGAHYPRSCCCCC | 33 | https://www.uniprot.org/uniprot/A8JUP5 | SubName: Full=Uncharacterized protein, i | Cell membrane: 0.7726, Golgi apparatus: 0.0981, Nucleus: 0.0942, Lysosome/Vacuole: 0.0147, Cytoplasm: 0.0071, Endoplasmic reticulum: 0.0062, Mitochondrion: 0.0045, Extracellular: 0.0021, Peroxisome: 0.0002, Plastid: 0.0001 |
| | Q59E13_DROME | Nucleus, Membrane | 13,7 | 0,69 | 0 | LQLLLHLLLLPELLLLRAGL | 20 | AAGN | 4 | https://www.uniprot.org/uniprot/Q59E13 | SubName: Full=Defective proboscis extens | Nucleus: 0.4218, Cell membrane: 0.2475, Endoplasmic reticulum: 0.1736, Cytoplasm: 0.0733, Golgi apparatus: 0.0612, Peroxisome: 0.0098, Lysosome/Vacuole: 0.0061, Extracellular: 0.0037, Mitochondrion: 0.0028, Plastid: 0.0004 |
| fan | Q9VSD3_DROME | Endoplasmic reticulum, Membrane | 13,8 | 0,55 | 0 | TFFYMAAIVIIILSAVAGAYYGKTR | 25 | L | 1 | https://www.uniprot.org/uniprot/Q9VSD3 | SubName: Full=AT11025p {ECO:0000313\|I | Endoplasmic reticulum: 0.4773, Mitochondrion: 0.2266, Peroxisome: 0.0865, Plastid: 0.0764, Cell membrane: 0.0628, Lysosome/Vacuole: 0.0454, Cytoplasm: 0.0117, Golgi apparatus: 0.0077, Nucleus: 0.0056, Extracellular: 0.0001 |
| | ATPK_DROME | Mitochondrion, Membrane | 13,8 | 0,66 | 4 | GIAPFFQLTVASMTFFYLINY | 21 | TKLKHHRNYKYH | 12 | https://www.uniprot.org/uniprot/Q9W141 | RecName: Full=Putative ATP synthase sub | Mitochondrion: 0.7254, Endoplasmic reticulum: 0.1089, Peroxisome: 0.0877, Cytoplasm: 0.0383, Extracellular: 0.0148, Cell membrane: 0.0091, Golgi apparatus: 0.0067, Nucleus: 0.0063, Plastid: 0.0017, Lysosome/Vacuole: 0.0012 |
| | A0A126GV01_DROME | Endoplasmic reticulum, Membrane | 13,9 | 0,63 | 0 | QTTLLLVLMIEVMLTFNIPIPM | 22 | VIGAGI | 6 | https://www.uniprot.org/uniprot/A0A126GV01 | SubName: Full=Uncharacterized protein, i | Endoplasmic reticulum: 0.2841, Lysosome/Vacuole: 0.282, Extracellular: 0.1544, Cell membrane: 0.1077, Golgi apparatus: 0.0931, Mitochondrion: 0.0485, Plastid: 0.0147, Nucleus: 0.0086, Cytoplasm: 0.0057, Peroxisome: 0.0012 |
| | Q8IMU2_DROME | Endoplasmic reticulum, Membrane | 14,1 | 0,61 | 4 | ITSLATLSIFALVALLFVGKKVA | 23 | KIFSSEPAKTKNKSK | 15 | https://www.uniprot.org/uniprot/Q8IMU2 | SubName: Full=RE10237p {ECO:0000313\|I | Endoplasmic reticulum: 0.5495, Peroxisome: 0.2059, Mitochondrion: 0.1266, Cell membrane: 0.0325, Cytoplasm: 0.0236, Nucleus: 0.019, Golgi apparatus: 0.017, Plastid: 0.0134, Lysosome/Vacuole: 0.0125, Extracellular: 0.0001 |
| | Q8SXV1_DROME | Endoplasmic reticulum, Membrane | 14,3 | 0,68 | 2 | QGLVSFLVSICSLLAPFLVSA | 21 | HMRKHF | 6 | https://www.uniprot.org/uniprot/Q8SXV1 | SubName: Full=GEO13282p1 {ECO:000031 | Endoplasmic reticulum: 0.2941, Cell membrane: 0.2893, Mitochondrion: 0.1395, Plastid: 0.0909, Lysosome/Vacuole: 0.069, Extracellular: 0.0502, Golgi apparatus: 0.0306, Peroxisome: 0.0268, Nucleus: 0.0059, Cytoplasm: 0.0036 |
| | B7YZN6_DROME | Mitochondrion, Membrane | 14,3 | 0,72 | 0 | LFQISFITGSLLIFFSKPLF | 20 | DCFIADPLPPPENKVPPHKR | 20 | https://www.uniprot.org/uniprot/B7YZN6 | SubName: Full=GH02741p2 {ECO:0000313 | Mitochondrion: 0.5635, Nucleus: 0.2253, Golgi apparatus: 0.0533, Cytoplasm: 0.0483, Peroxisome: 0.0415, Endoplasmic reticulum: 0.0375, Extracellular: 0.0125, Cell membrane: 0.0104, Plastid: 0.0045, Lysosome/Vacuole: 0.003 |
| | M9MS32_DROME | Endoplasmic reticulum, Membrane | 14,4 | 0,69 | 2 | SYGVVDVQICISLLVLTITFI | 21 | RKA | 3 | https://www.uniprot.org/uniprot/M9MS32 | SubName: Full=Uncharacterized protein, i | Endoplasmic reticulum: 0.3749, Peroxisome: 0.1414, Cell membrane: 0.1254, Lysosome/Vacuole: 0.1103, Cytoplasm: 0.069, Mitochondrion: 0.0558, Plastid: 0.053, Golgi apparatus: 0.0318, Nucleus: 0.0262, Extracellular: 0.0122 |
| MIC10 | Q8SY69_DROME | Mitochondrion, Membrane | 14,5 | 0,73 | 4 | VLIKGCGGVIIGSAVSFLIL | 20 | KRRAWPVWLGAGFGMGIAYRTCEKDLN | 30 | https://www.uniprot.org/uniprot/Q8SY69 | RecName: Full=MICOS complex subunit M | Mitochondrion: 0.916, Endoplasmic reticulum: 0.0299, Peroxisome: 0.0223, Plastid: 0.0142, Cell membrane: 0.006, Cytoplasm: 0.0037, Golgi apparatus: 0.0031, Lysosome/Vacuole: 0.0022, Extracellular: 0.0017, Nucleus: 0.0008 |
| | Q6IHK5_DROME | Golgi apparatus, Membrane | 14,6 | 0,77 | 0 | LPNFKILMCCCYFIIIVLM | 19 | I | 1 | https://www.uniprot.org/uniprot/Q6IHK5 | SubName: Full=HDC02355 {ECO:0000313\| | Golgi apparatus: 0.3681, Extracellular: 0.2495, Cell membrane: 0.2103, Lysosome/Vacuole: 0.0946, Mitochondrion: 0.0376, Endoplasmic reticulum: 0.0215, Nucleus: 0.0134, Cytoplasm: 0.0024, Plastid: 0.0024, Peroxisome: 0.0002 |
| | A0A0B4KEL2_DROME | Peroxisome, Membrane | 14,6 | 0,97 | 1 | ALILGSAFLLEYLFF | 15 | QTRF | 4 | https://www.uniprot.org/uniprot/A0A0B4KEL2 | RecName: Full=Phospholipid scramblase {I | Peroxisome: 0.2771, Mitochondrion: 0.2091, Cytoplasm: 0.1821, Nucleus: 0.1673, Endoplasmic reticulum: 0.0628, Golgi apparatus: 0.0292, Cell membrane: 0.0242, Plastid: 0.0237, Lysosome/Vacuole: 0.0161, Extracellular: 0.0083 |
| | A1Z8F5_DROME | Peroxisome, Membrane | 14,6 | 0,97 | 1 | ALILGSAFLLEYLFF | 15 | QTRF | 4 | https://www.uniprot.org/uniprot/A1Z8F5 | RecName: Full=Phospholipid scramblase {I | Peroxisome: 0.2758, Mitochondrion: 0.1979, Cytoplasm: 0.1807, Nucleus: 0.167, Endoplasmic reticulum: 0.0676, Golgi apparatus: 0.0325, Plastid: 0.0263, Cell membrane: 0.0251, Lysosome/Vacuole: 0.019, Extracellular: 0.0083 |
| | A0A0B4KH06_DROME | Cell membrane, Membrane | 14,7 | 0,82 | 0 | LLYAAMLALAYLSSWTTL | 18 | RMSDVATXQPHS | 12 | https://www.uniprot.org/uniprot/A0A0B4KH06 | SubName: Full=Glial cell line-derived neur | Cell membrane: 0.4899, Extracellular: 0.3123, Nucleus: 0.0731, Endoplasmic reticulum: 0.0497, Cytoplasm: 0.0352, Golgi apparatus: 0.0286, Lysosome/Vacuole: 0.0052, Mitochondrion: 0.0043, Peroxisome: 0.0013, Plastid: 0.0002 |
| | A0A0B4K785_DROME | Cell membrane, Membrane | 14,7 | 0,82 | 0 | LLYAAMLALAYLSSWTTL | 18 | RMSDVAT | 7 | https://www.uniprot.org/uniprot/A0A0B4K785 | SubName: Full=Glial cell line-derived neur | Cell membrane: 0.4871, Extracellular: 0.3296, Nucleus: 0.0619, Endoplasmic reticulum: 0.0511, Cytoplasm: 0.0348, Golgi apparatus: 0.024, Lysosome/Vacuole: 0.0056, Mitochondrion: 0.004, Peroxisome: 0.0016, Plastid: 0.0003 |
| | A0A126GUZ9_DROME | Mitochondrion, Membrane | 14,8 | 0,62 | -2 | FRFLKVINLGLPMLFYGLAAIRLI | 24 | SHTEIVHVSETVKVPIYFLYPEDKGSSF | 28 | https://www.uniprot.org/uniprot/A0A126GUZ9 | SubName: Full=Uncharacterized protein, i | Mitochondrion: 0.3336, Endoplasmic reticulum: 0.2844, Cytoplasm: 0.1778, Extracellular: 0.0429, Nucleus: 0.0428, Peroxisome: 0.0379, Lysosome/Vacuole: 0.0319, Golgi apparatus: 0.0306, Cell membrane: 0.011, Plastid: 0.0069 |
| | Q9VY65_DROME | Mitochondrion, Membrane | 14,9 | 0,75 | 5 | SLVKGVGGLVIGSVVTLLFF | 20 | RRRIWPVWLGTGFGVGVAYRGCEKELNI | 35 | https://www.uniprot.org/uniprot/Q9VY65 | RecName: Full=MICOS complex subunit M | Mitochondrion: 0.9661, Endoplasmic reticulum: 0.0169, Plastid: 0.0075, Peroxisome: 0.007, Cell membrane: 0.001, Golgi apparatus: 0.0008, Lysosome/Vacuole: 0.0003, Cytoplasm: 0.0002, Nucleus: 0.0001, Extracellular: 0.0 |
| CG7130 | Q9VNW0_DROME | Mitochondrion, Membrane | 14,9 | 0,78 | 1 | MLPFMCAVGGTVLFAFAAY | 19 | KTFQFFNRKKEATHGDGSSSD | 21 | https://www.uniprot.org/uniprot/Q9VNW0 | SubName: Full=GEO11142p1 {ECO:000031 | Mitochondrion: 0.6002, Endoplasmic reticulum: 0.1984, Peroxisome: 0.1056, Plastid: 0.0328, Cell membrane: 0.0226, Cytoplasm: 0.0222, Nucleus: 0.0072, Golgi apparatus: 0.0065, Lysosome/Vacuole: 0.0044, Extracellular: 0.0001 |
| | Q9W380_DROME | Mitochondrion, Membrane | 15 | 0,65 | 0 | IVGGSALALAGIVMATIGVANYY | 23 | AKDGDNRRYKLGYVVYRHDDPRALKVRN | 32 | https://www.uniprot.org/uniprot/Q9W380 | SubName: Full=RH45008p {ECO:0000313\| | Mitochondrion: 0.9342, Plastid: 0.0281, Peroxisome: 0.0128, Cytoplasm: 0.0086, Extracellular: 0.0049, Endoplasmic reticulum: 0.003, Nucleus: 0.0029, Lysosome/Vacuole: 0.0029, Cell membrane: 0.0013, Golgi apparatus: 0.0013 |
| | A0A0B4LES9_DROME | Peroxisome, Membrane | 15 | 0,65 | 5 | YGPVLCGTILLGWSLALWTIIHS | 23 | MANTRRINQRLNEQRLLQQKVK | 22 | https://www.uniprot.org/uniprot/A0A0B4LES9 | SubName: Full=Phosphatidylinositol glycar | Peroxisome: 0.4865, Mitochondrion: 0.2409, Endoplasmic reticulum: 0.1095, Cytoplasm: 0.0608, Cell membrane: 0.0423, Nucleus: 0.0285, Golgi apparatus: 0.0131, Plastid: 0.0108, Lysosome/Vacuole: 0.0068, Extracellular: 0.0007 |
| PIG-X | A8DY62_DROME | Peroxisome, Membrane | 15 | 0,65 | 5 | YGPVLCGTILLGWSLALWTIIHS | 23 | MANTRRINQRLNEQRLLQQKVK | 22 | https://www.uniprot.org/uniprot/A8DY62 | SubName: Full=Phosphatidylinositol glycar | Peroxisome: 0.5126, Mitochondrion: 0.2438, Endoplasmic reticulum: 0.1328, Cytoplasm: 0.042, Cell membrane: 0.0241, Nucleus: 0.0174, Golgi apparatus: 0.0109, Lysosome/Vacuole: 0.0088, Plastid: 0.0073, Extracellular: 0.0003 |
| | A1Z8I6_DROME | Mitochondrion, Membrane | 15 | 0,75 | 8 | FYVPAIVVFTLAALFGYKLY | 20 | KSLTEKELKKQEKLKSKQQKKAKKSN | 26 | https://www.uniprot.org/uniprot/A1Z8I6 | SubName: Full=Uncharacterized protein, i | Mitochondrion: 0.6298, Endoplasmic reticulum: 0.2647, Golgi apparatus: 0.0412, Plastid: 0.03, Cell membrane: 0.0234, Peroxisome: 0.0033, Extracellular: 0.0031, Lysosome/Vacuole: 0.0019, Nucleus: 0.0017, Cytoplasm: 0.0009 |
| | Q7K036_DROME | Endoplasmic reticulum, Membrane | 15 | 0,75 | 0 | VVQTGFHWGFVPLVLYLGFM | 20 | KGAEPGMPPLNLFSLLWQ | 18 | https://www.uniprot.org/uniprot/Q7K036 | SubName: Full=AT23266p {ECO:0000313\|I | Endoplasmic reticulum: 0.4432, Mitochondrion: 0.4225, Golgi apparatus: 0.0884, Peroxisome: 0.0172, Extracellular: 0.0111, Cell membrane: 0.0071, Lysosome/Vacuole: 0.0046, Plastid: 0.0032, Cytoplasm: 0.0022, Nucleus: 0.0003 |
| CG11327 | Q9VM98_DROME | Golgi apparatus, Membrane | 15 | 0,75 | 1 | LKLSYPILFCSLAFSLVYII | 20 | RKE | 3 | https://www.uniprot.org/uniprot/Q9VM98 | SubName: Full=AT06841p1 {ECO:0000313 | Golgi apparatus: 0.5744, Endoplasmic reticulum: 0.2246, Mitochondrion: 0.0874, Nucleus: 0.082, Peroxisome: 0.0106, Cytoplasm: 0.009, Lysosome/Vacuole: 0.0047, Plastid: 0.0045, Cell membrane: 0.0028, Extracellular: 0.0 |
| | PTP61_DROME | Mitochondrion, Membrane | 15,1 | 0,76 | 1 | LLTYIAAGVVVGVICAYAYT | 20 | KLG | 3 | https://www.uniprot.org/uniprot/Q9W0G1 | RecName: Full=Tyrosine-protein phosphat | Mitochondrion: 0.2958, Nucleus: 0.2608, Peroxisome: 0.1701, Endoplasmic reticulum: 0.1669, Golgi apparatus: 0.0535, Cell membrane: 0.0373, Cytoplasm: 0.0073, Lysosome/Vacuole: 0.0043, Plastid: 0.004, Extracellular: 0.0001 |
| | Q8IRH4_DROME | Mitochondrion, Membrane | 15,1 | 0,76 | 1 | LLTYIAAGVVVGVICAYAYT | 20 | KLG | 3 | https://www.uniprot.org/uniprot/Q8IRH4 | SubName: Full=Protein tyrosine phosphat | Mitochondrion: 0.3687, Nucleus: 0.3004, Peroxisome: 0.135, Endoplasmic reticulum: 0.1052, Golgi apparatus: 0.0587, Cell membrane: 0.0231, Cytoplasm: 0.004, Plastid: 0.0024, Lysosome/Vacuole: 0.0022, Extracellular: 0.0002 |
| | M9NEV7_DROME | Cell membrane, Membrane | 15,1 | 0,79 | 0 | SLAPIAGAILLFFGVARLL | 19 | A | 1 | https://www.uniprot.org/uniprot/M9NEV7 | SubName: Full=Uncharacterized protein, i | Cell membrane: 0.3965, Mitochondrion: 0.2404, Endoplasmic reticulum: 0.1486, Peroxisome: 0.076, Golgi apparatus: 0.0338, Nucleus: 0.0306, Cytoplasm: 0.0258, Lysosome/Vacuole: 0.0233, Extracellular: 0.0186, Plastid: 0.0063 |
| | Q9VZF2_DROME | Nucleus, Membrane | 15,2 | 0,76 | 0 | IYAVSILTVVGAIVVAFYAD | 20 | S | 1 | https://www.uniprot.org/uniprot/Q9VZF2 | SubName: Full=LD05244p {ECO:0000313\|I | Nucleus: 0.8211, Cell membrane: 0.1173, Cytoplasm: 0.0267, Golgi apparatus: 0.0215, Mitochondrion: 0.0054, Endoplasmic reticulum: 0.0039, Extracellular: 0.0036, Plastid: 0.0003, Peroxisome: 0.0002, Lysosome/Vacuole: 0.0 |
| | Q9W221_DROME | Cell membrane, Membrane | 15,3 | 0,73 | -2 | CWALILCLFICWPCVCLPYCM | 21 | DSCQNANHYCPNCSAYIGTYEN | 22 | https://www.uniprot.org/uniprot/Q9W221 | SubName: Full=MIP04282p {ECO:0000313 | Cell membrane: 0.6649, Extracellular: 0.1832, Cytoplasm: 0.044, Nucleus: 0.0359, Golgi apparatus: 0.0273, Lysosome/Vacuole: 0.0184, Mitochondrion: 0.0154, Endoplasmic reticulum: 0.0106, Plastid: 0.0002, Peroxisome: 0.0001 |
| CG43321 | M9NEV9_DROME | Mitochondrion, Membrane | 15,3 | 0,73 | -4 | YKYIAVFITVIPGIILGGFMG | 21 | KKLAQFLEVFDLYAPDVSEDD | 21 | https://www.uniprot.org/uniprot/M9NEV9 | SubName: Full=Uncharacterized protein {E | Mitochondrion: 0.44, Golgi apparatus: 0.2526, Endoplasmic reticulum: 0.077, Nucleus: 0.0707, Extracellular: 0.0448, Cytoplasm: 0.0445, Peroxisome: 0.028, Lysosome/Vacuole: 0.0248, Cell membrane: 0.0117, Plastid: 0.0059 |
| CG30269 | Q9W217_DROME | Cell membrane, Membrane | 15,4 | 0,70 | 1 | HLIALILCLFQLYCCVCLPYCI | 22 | SSCMNTNHYCGMCDRYLGTYDRK | 23 | https://www.uniprot.org/uniprot/Q9W217 | SubName: Full=Uncharacterized protein, i | Cell membrane: 0.5969, Golgi apparatus: 0.1695, Nucleus: 0.0943, Extracellular: 0.0644, Cytoplasm: 0.0251, Mitochondrion: 0.0225, Endoplasmic reticulum: 0.0159, Lysosome/Vacuole: 0.0101, Peroxisome: 0.0009, Plastid: 0.0004 |
| | A0A126GUZ5_DROME | Endoplasmic reticulum, Membrane | 15,4 | 0,73 | 0 | VQTTLLLVLMIEVMLTFNIPI | 21 | PMVIGAGI | 8 | https://www.uniprot.org/uniprot/A0A126GUZ5 | SubName: Full=Uncharacterized protein, i | Endoplasmic reticulum: 0.5329, Golgi apparatus: 0.1655, Lysosome/Vacuole: 0.1185, Cell membrane: 0.0813, Mitochondrion: 0.061, Extracellular: 0.0308, Peroxisome: 0.0035, Plastid: 0.0023, Cytoplasm: 0.0022, Nucleus: 0.002 |
| CG5050 | Q9VJD8_DROME | Endoplasmic reticulum, Membrane | 15,4 | 0,81 | 2 | YSFNVIYLFALYLLPGMSL | 19 | KYRQN | 5 | https://www.uniprot.org/uniprot/Q9VJD8 | SubName: Full=AT13217p1 {ECO:0000313 | Endoplasmic reticulum: 0.2851, Cytoplasm: 0.2223, Peroxisome: 0.1276, Mitochondrion: 0.1244, Nucleus: 0.1217, Golgi apparatus: 0.0943, Cell membrane: 0.01, Lysosome/Vacuole: 0.0082, Plastid: 0.0053, Extracellular: 0.0012 |
| | A0A0B4K870_DROME | Cell membrane, Membrane | 15,6 | 0,58 | 6 | LGQLSCLLSTLSCCFPIFALTFVYSCL | 27 | QSRLKSKRVFCSRCGGHLGFHWRPT | 25 | https://www.uniprot.org/uniprot/A0A0B4K870 | SubName: Full=Uncharacterized protein, i | Cell membrane: 0.5366, Endoplasmic reticulum: 0.1492, Golgi apparatus: 0.1022, Mitochondrion: 0.089, Nucleus: 0.0878, Lysosome/Vacuole: 0.0137, Peroxisome: 0.0108, Plastid: 0.0065, Cytoplasm: 0.0042, Extracellular: 0.0 |
| | M9PB41_DROME | Endoplasmic reticulum, Membrane | 15,6 | 0,92 | 1 | IRTYYILVYILLSVWCF | 17 | R | 1 | https://www.uniprot.org/uniprot/M9PB41 | SubName: Full=Uncharacterized protein, i | Endoplasmic reticulum: 0.3631, Golgi apparatus: 0.3319, Cell membrane: 0.0904, Nucleus: 0.0765, Cytoplasm: 0.0528, Lysosome/Vacuole: 0.0482, Mitochondrion: 0.0212, Peroxisome: 0.0107, Plastid: 0.0046, Extracellular: 0.0006 |
| | A0A0B4LG03_DROME | Endoplasmic reticulum, Membrane | 15,7 | 0,68 | 2 | NLWLILRLVIMACSVCALIYHGY | 23 | FLSKIGKMYL | 10 | https://www.uniprot.org/uniprot/A0A0B4LG03 | SubName: Full=Uncharacterized protein, i | Endoplasmic reticulum: 0.6056, Mitochondrion: 0.1812, Peroxisome: 0.1417, Nucleus: 0.0184, Golgi apparatus: 0.0171, Plastid: 0.0168, Cytoplasm: 0.0103, Lysosome/Vacuole: 0.0043, Cell membrane: 0.0039, Extracellular: 0.0006 |
| | CYB5_DROME | Endoplasmic reticulum, Membrane | 15,8 | 0,72 | 1 | VKSWLVPLVLCLVATLFYKFFF | 22 | GGAKQ | 5 | https://www.uniprot.org/uniprot/Q9V4N3 | RecName: Full=Cytochrome b5; | Endoplasmic reticulum: 0.8991, Lysosome/Vacuole: 0.0278, Golgi apparatus: 0.0238, Cell membrane: 0.0185, Mitochondrion: 0.0151, Peroxisome: 0.0081, Nucleus: 0.0043, Cytoplasm: 0.0029, Plastid: 0.0005, Extracellular: 0.0 |
| | COA3_DROME | Mitochondrion, Membrane | 15,8 | 0,75 | -2 | LLTAGALGVSVLAIYGYSIFS | 21 | VQQEKFLDDFEEPKKVSS | 18 | https://www.uniprot.org/uniprot/P0DKM0 | RecName: Full=Cytochrome c oxidase asse | Mitochondrion: 0.9535, Nucleus: 0.0244, Endoplasmic reticulum: 0.0118, Golgi apparatus: 0.0036, Cytoplasm: 0.0023, Plastid: 0.0019, Peroxisome: 0.0019, Cell membrane: 0.0005, Extracellular: 0.0001, Lysosome/Vacuole: 0.0001 |
| | FIS1_DROME | Mitochondrion, Membrane | 15,9 | 0,64 | 4 | AKGMVVAGGAALVLGGILGLGIAMA | 25 | RNKQKREK | 8 | https://www.uniprot.org/uniprot/B7YZT2 | RecName: Full=Mitochondrial fission 1 pro | Mitochondrion: 0.4743, Peroxisome: 0.2869, Plastid: 0.0824, Cytoplasm: 0.077, Endoplasmic reticulum: 0.0483, Cell membrane: 0.0134, Lysosome/Vacuole: 0.0131, Nucleus: 0.0035, Golgi apparatus: 0.0009, Extracellular: 0.0002 |
| | Q9VEQ3_DROME | Endoplasmic reticulum, Membrane | 16 | 0,67 | 0 | SATVAMCATCVAVVAVSVFLPWYL | 24 | MEKVGSL | 7 | https://www.uniprot.org/uniprot/Q9VEQ3 | SubName: Full=GEO09703p1 {ECO:000031 | Endoplasmic reticulum: 0.366, Mitochondrion: 0.3526, Plastid: 0.0815, Cell membrane: 0.073, Golgi apparatus: 0.056, Peroxisome: 0.0295, Lysosome/Vacuole: 0.0213, Nucleus: 0.0152, Cytoplasm: 0.0043, Extracellular: 0.0006 |
| zda | Q7K3D4_DROME | Mitochondrion, Membrane | 16,1 | 0,77 | 1 | LLGYLMGTILIGVAGVAIYRY | 21 | KY | 2 | https://www.uniprot.org/uniprot/Q7K3D4 | RecName: Full=Peptidylprolyl isomerase {E | Mitochondrion: 0.4918, Endoplasmic reticulum: 0.3563, Peroxisome: 0.0814, Golgi apparatus: 0.0229, Lysosome/Vacuole: 0.0164, Cell membrane: 0.0104, Plastid: 0.0095, Nucleus: 0.0076, Cytoplasm: 0.0036, Extracellular: 0.0 |
| | Q8I079_DROME | Mitochondrion, Membrane | 16,3 | 0,65 | 0 | TFSTVCAVGGVLVGLFLGFGAFKAL | 25 | SGSNNNH | 7 | https://www.uniprot.org/uniprot/Q8I079 | SubName: Full=Uncharacterized protein {E | Mitochondrion: 0.4231, Endoplasmic reticulum: 0.2203, Plastid: 0.1087, Peroxisome: 0.0855, Cell membrane: 0.0563, Golgi apparatus: 0.0398, Lysosome/Vacuole: 0.0312, Cytoplasm: 0.0179, Nucleus: 0.0164, Extracellular: 0.0009 |
| | Q8IN56_DROME | Endoplasmic reticulum, Membrane | 16,3 | 0,82 | 1 | KSLIYALIGLAFSLILRWLF | 20 | K | 1 | https://www.uniprot.org/uniprot/Q8IN56 | RecName: Full=Oxysterol-binding protein | Endoplasmic reticulum: 0.3905, Cell membrane: 0.1783, Peroxisome: 0.1627, Nucleus: 0.0917, Golgi apparatus: 0.0705, Mitochondrion: 0.0646, Lysosome/Vacuole: 0.0204, Cytoplasm: 0.0178, Plastid: 0.0032, Extracellular: 0.0002 |
| | Q9W4N8_DROME | Endoplasmic reticulum, Membrane | 16,5 | 0,79 | 0 | PVFYIAVAIAAAIVSLLLGKF | 21 | FL | 2 | https://www.uniprot.org/uniprot/Q9W4N8 | SubName: Full=LD30122p {ECO:0000313\|I | Endoplasmic reticulum: 0.4916, Mitochondrion: 0.153, Peroxisome: 0.1262, Golgi apparatus: 0.1026, Cell membrane: 0.0574, Plastid: 0.0269, Lysosome/Vacuole: 0.023, Cytoplasm: 0.0097, Nucleus: 0.0095, Extracellular: 0.0001 |
| | A0A0B4JCV9_DROME | Endoplasmic reticulum, Membrane | 16,5 | 0,83 | 0 | LPWWCALLESLFLLLLVKLL | 20 | QL | 2 | https://www.uniprot.org/uniprot/A0A0B4JCV9 | SubName: Full=Uncharacterized protein {E | Endoplasmic reticulum: 0.5952, Cell membrane: 0.1419, Mitochondrion: 0.0956, Golgi apparatus: 0.0851, Peroxisome: 0.0314, Cytoplasm: 0.017, Nucleus: 0.0168, Extracellular: 0.0089, Lysosome/Vacuole: 0.0077, Plastid: 0.0003 |
| | CDKAL_DROME | Endoplasmic reticulum, Membrane | 16,6 | 0,75 | 0 | RYFGIALVLGSLAFLIQLVVRL | 22 | L | 1 | https://www.uniprot.org/uniprot/Q7K4W1 | RecName: Full=Threonylcarbamoyladenos | Endoplasmic reticulum: 0.5188, Nucleus: 0.1629, Golgi apparatus: 0.1082, Peroxisome: 0.0768, Mitochondrion: 0.0596, Cell membrane: 0.0375, Cytoplasm: 0.0176, Lysosome/Vacuole: 0.0126, Plastid: 0.0058, Extracellular: 0.0001 |
| | HID_DROME | Nucleus, Membrane | 16,6 | 0,79 | 1 | FTWPTVVTVFVLAMGCGFFAA | 21 | R | 1 | https://www.uniprot.org/uniprot/Q24106 | RecName: Full=Cell death protein hid; | Nucleus: 0.3518, Cell membrane: 0.2996, Endoplasmic reticulum: 0.118, Mitochondrion: 0.1027, Golgi apparatus: 0.0852, Peroxisome: 0.0225, Plastid: 0.0119, Cytoplasm: 0.0051, Lysosome/Vacuole: 0.0029, Extracellular: 0.0001 |
| hid | M9PFL5_DROME | Nucleus, Membrane | 16,6 | 0,79 | 1 | FTWPTVVTVFVLAMGCGFFAA | 21 | R | 1 | https://www.uniprot.org/uniprot/M9PFL5 | SubName: Full=Head involution defective, | Nucleus: 0.3518, Cell membrane: 0.2996, Endoplasmic reticulum: 0.118, Mitochondrion: 0.1027, Golgi apparatus: 0.0852, Peroxisome: 0.0225, Plastid: 0.0119, Cytoplasm: 0.0051, Lysosome/Vacuole: 0.0029, Extracellular: 0.0001 |
| Tango11 | A0A0B4LG59_DROME | Mitochondrion, Membrane | 16,7 | 0,93 | 1 | IVYCLGLAYFVLKTIFWL | 18 | NRN | 3 | https://www.uniprot.org/uniprot/A0A0B4LG59 | SubName: Full=Transport and golgi organi | Mitochondrion: 0.4136, Endoplasmic reticulum: 0.1614, Peroxisome: 0.1142, Cell membrane: 0.1111, Golgi apparatus: 0.083, Nucleus: 0.0564, Cytoplasm: 0.0377, Extracellular: 0.0099, Lysosome/Vacuole: 0.0096, Plastid: 0.0031 |
| CG34242 | A1A6X2_DROME | Mitochondrion, Membrane | 16,8 | 0,70 | 2 | GWRFVGFVSCIVGAVGLTLYPVIV | 24 | DPMVNTEKYKTLQEYSKIKRDELQHIKRQ | 29 | https://www.uniprot.org/uniprot/A1A6X2 | SubName: Full=IP16893p {ECO:0000313\|E | Mitochondrion: 0.9691, Plastid: 0.0066, Endoplasmic reticulum: 0.0062, Golgi apparatus: 0.0049, Extracellular: 0.0036, Peroxisome: 0.0034, Cytoplasm: 0.0025, Nucleus: 0.0018, Cell membrane: 0.0015, Lysosome/Vacuole: 0.0004 |
| | Q8MSQ3_DROME | Cell membrane, Membrane | 16,8 | 0,80 | 1 | ALKIITFMLIGSAFLITSAAI | 21 | KQAYY | 5 | https://www.uniprot.org/uniprot/Q8MSQ3 | SubName: Full=SD10416p {ECO:0000313\|I | Cell membrane: 0.4739, Nucleus: 0.3589, Endoplasmic reticulum: 0.0562, Mitochondrion: 0.0314, Golgi apparatus: 0.0298, Extracellular: 0.0196, Cytoplasm: 0.0154, Peroxisome: 0.0083, Plastid: 0.004, Lysosome/Vacuole: 0.0024 |
| | M9PB07_DROME | Endoplasmic reticulum, Membrane | 16,9 | 0,85 | 3 | TILYYTTLAVGFSCAWFWWW | 20 | TKFRSAPRLERISVQT | 16 | https://www.uniprot.org/uniprot/M9PB07 | SubName: Full=Uncharacterized protein, i | Endoplasmic reticulum: 0.4465, Mitochondrion: 0.1622, Peroxisome: 0.1418, Cell membrane: 0.1255, Golgi apparatus: 0.0528, Plastid: 0.0389, Nucleus: 0.0155, Lysosome/Vacuole: 0.0091, Cytoplasm: 0.0073, Extracellular: 0.0004 |
| | Q95TS8_DROME | Endoplasmic reticulum, Membrane | 16,9 | 0,85 | 3 | TILYYTTLAVGFSCAWFWWW | 20 | TKFRSAPRLERISVQT | 16 | https://www.uniprot.org/uniprot/Q95TS8 | SubName: Full=LD25288p {ECO:0000313\|I | Endoplasmic reticulum: 0.4385, Peroxisome: 0.1584, Mitochondrion: 0.1451, Cell membrane: 0.131, Golgi apparatus: 0.0517, Plastid: 0.0377, Nucleus: 0.0198, Lysosome/Vacuole: 0.0091, Cytoplasm: 0.008, Extracellular: 0.0005 |
| | M9MS63_DROME | Endoplasmic reticulum, Membrane | 17,2 | 0,72 | 1 | FLLKLRFLAQVIYTLVAVLSVIHV | 24 | K | 1 | https://www.uniprot.org/uniprot/M9MS63 | SubName: Full=Uncharacterized protein, i | Endoplasmic reticulum: 0.5473, Mitochondrion: 0.2952, Peroxisome: 0.0866, Extracellular: 0.0258, Golgi apparatus: 0.0244, Cytoplasm: 0.013, Nucleus: 0.0037, Cell membrane: 0.0032, Lysosome/Vacuole: 0.0006, Plastid: 0.0002 |
| | M9MSP0_DROME | Endoplasmic reticulum, Membrane | 17,2 | 0,75 | 1 | FLLKLRFLAQVIYTLVAVVIGTL | 23 | KT | 2 | https://www.uniprot.org/uniprot/M9MSP0 | SubName: Full=GEO02690p1 {ECO:000031 | Endoplasmic reticulum: 0.4819, Mitochondrion: 0.3201, Peroxisome: 0.0824, Golgi apparatus: 0.0475, Extracellular: 0.0334, Cytoplasm: 0.0249, Nucleus: 0.0043, Cell membrane: 0.0042, Lysosome/Vacuole: 0.001, Plastid: 0.0003 |

| | | | | | | | | | | | | |
|---|---|---|---|---|---|---|---|---|---|---|---|---|
| | E1JGT2_DROME | Endoplasmic reticulum, Membrane | 17,3 | 0,75 | 3 | WHKVVFFMLGLISALTICVLVRL | 23 | SNRLASDAEKKALMHFRELSSGRNGHRF | 30 | https://www.uniprot.org/uniprot/E1JGT2 | SubName: Full=Uncharacterized protein {E | Endoplasmic reticulum: 0.3822, Golgi apparatus: 0.2389, Mitochondrion: 0.2051, Cell membrane: 0.0915, Lysosome/Vacuole: 0.0253, Extracellular: 0.0191, Peroxisome: 0.0136, Plastid: 0.0131, Nucleus: 0.0073, Cytoplasm: 0.0039 |
| | Q9W5R5_DROME | Golgi apparatus, Membrane | 17,4 | 0,83 | 1 | LLCRFLPLGCVAIGLLIYFLS | 21 | SRI | 3 | https://www.uniprot.org/uniprot/Q9W5R5 | SubName: Full=LD47843p {ECO:0000313| | Golgi apparatus: 0.4429, Endoplasmic reticulum: 0.2992, Nucleus: 0.0872, Cytoplasm: 0.077, Mitochondrion: 0.0473, Lysosome/Vacuole: 0.0224, Peroxisome: 0.0102, Cell membrane: 0.009, Plastid: 0.0046, Extracellular: 0.0 |
| | X2JDM7_DROME | Endoplasmic reticulum, Membrane | 17,6 | 0,80 | 3 | WPLLMQLLLFVLWLNAKFWQLV | 22 | NEQVTYRRRRWH | 12 | https://www.uniprot.org/uniprot/X2JDM7 | RecName: Full=Guanine nucleotide-bindin | Endoplasmic reticulum: 0.3178, Nucleus: 0.2819, Peroxisome: 0.2372, Mitochondrion: 0.0477, Cytoplasm: 0.045, Golgi apparatus: 0.0391, Cell membrane: 0.0245, Lysosome/Vacuole: 0.0045, Extracellular: 0.0019, Plastid: 0.0003 |
| | M9PDW0_DROME | Endoplasmic reticulum, Membrane | 17,7 | 0,84 | 0 | LKMMIIMGVIGLVVVGIIASN | 21 | FM | 2 | https://www.uniprot.org/uniprot/M9PDW0 | SubName: Full=Neuronal synaptobrevin, is | Endoplasmic reticulum: 0.3998, Golgi apparatus: 0.3247, Mitochondrion: 0.1212, Cell membrane: 0.0639, Lysosome/Vacuole: 0.0418, Nucleus: 0.0298, Plastid: 0.0088, Peroxisome: 0.0053, Cytoplasm: 0.0035, Extracellular: 0.0012 |
| CG12012 | Q1ECB1_DROME | Cell membrane, Membrane | 17,8 | 0,77 | 2 | FSACGLCCAIFFFPLGILCCLAM | 23 | REKRCSNCGTVF | 12 | https://www.uniprot.org/uniprot/Q1ECB1 | SubName: Full=IP05417p {ECO:0000313|E | Cell membrane: 0.8955, Golgi apparatus: 0.0462, Lysosome/Vacuole: 0.0257, Extracellular: 0.0181, Mitochondrion: 0.005, Endoplasmic reticulum: 0.0043, Nucleus: 0.0028, Plastid: 0.0014, Cytoplasm: 0.0011, Peroxisome: 0.0001 |
| | Q9VZP9_DROME | Cell membrane, Membrane | 17,8 | 0,77 | 2 | FSACGLCCAIFFFPLGILCCLAM | 23 | REKRCSNCGTVF | 12 | https://www.uniprot.org/uniprot/Q9VZP9 | SubName: Full=IP05517p {ECO:0000313|E | Cell membrane: 0.9289, Golgi apparatus: 0.0204, Nucleus: 0.0115, Lysosome/Vacuole: 0.0115, Endoplasmic reticulum: 0.0076, Mitochondrion: 0.0074, Cytoplasm: 0.0073, Extracellular: 0.0028, Plastid: 0.0017, Peroxisome: 0.0009 |
| | Q6IK96_DROME | Cell membrane, Membrane | 18,2 | 1,21 | 1 | WVICLLGLVILLCCL | 15 | K | 1 | https://www.uniprot.org/uniprot/Q6IK96 | SubName: Full=FI16918p1 {ECO:0000313| | Cell membrane: 0.5043, Golgi apparatus: 0.1916, Endoplasmic reticulum: 0.1793, Mitochondrion: 0.0336, Nucleus: 0.0324, Lysosome/Vacuole: 0.0264, Peroxisome: 0.0131, Cytoplasm: 0.0106, Extracellular: 0.0082, Plastid: 0.0005 |
| | B8A3Y3_DROME | Endoplasmic reticulum, Membrane | 18,3 | 0,76 | 4 | FFANLIFFVAIVTPLYFCLLKDVI | 24 | AMITKPKKR | 9 | https://www.uniprot.org/uniprot/B8A3Y3 | SubName: Full=MIP01424p {ECO:0000313 | Endoplasmic reticulum: 0.4097, Mitochondrion: 0.2233, Golgi apparatus: 0.2172, Cell membrane: 0.13, Extracellular: 0.0119, Lysosome/Vacuole: 0.0033, Peroxisome: 0.0023, Plastid: 0.0012, Nucleus: 0.0006, Cytoplasm: 0.0005 |
| Vap33 | Q7KVX5_DROME | Endoplasmic reticulum, Membrane | 18,5 | 0,84 | 0 | PVFYIAVAIAAAIVSLLLGKFF | 22 | L | 1 | https://www.uniprot.org/uniprot/Q7KVX5 | SubName: Full=VAMP-associated protein | Endoplasmic reticulum: 0.6215, Mitochondrion: 0.1403, Golgi apparatus: 0.0799, Peroxisome: 0.076, Cell membrane: 0.0349, Lysosome/Vacuole: 0.0175, Plastid: 0.016, Cytoplasm: 0.0075, Nucleus: 0.0063, Extracellular: 0.0 |
| | Q9W1P0_DROME | Cell membrane, Membrane | 18,6 | 0,74 | -1 | WIAYVASGIVLFTTCGIGCWLIPCI | 25 | LDCFNEIHHSCPVCKATLGIVPQQDQRW | 30 | https://www.uniprot.org/uniprot/Q9W1P0 | SubName: Full=FI16602p1 {ECO:0000313| | Cell membrane: 0.4246, Mitochondrion: 0.206, Golgi apparatus: 0.1513, Lysosome/Vacuole: 0.0574, Extracellular: 0.0386, Endoplasmic reticulum: 0.0354, Nucleus: 0.0353, Plastid: 0.0323, Cytoplasm: 0.0127, Peroxisome: 0.0065 |
| | Q7JUM3_DROME | Endoplasmic reticulum, Membrane | 18,6 | 0,89 | 3 | GSWFLVFFVVAHFLMRLLELW | 21 | RPRKHVEIAQDVQRRAS | 17 | https://www.uniprot.org/uniprot/Q7JUM3 | SubName: Full=LD44262p {ECO:0000313| | Endoplasmic reticulum: 0.3292, Peroxisome: 0.2591, Nucleus: 0.2019, Cell membrane: 0.0721, Mitochondrion: 0.0511, Cytoplasm: 0.0509, Golgi apparatus: 0.025, Lysosome/Vacuole: 0.0057, Extracellular: 0.0028, Plastid: 0.0023 |
| | Q8IPS9_DROME | Mitochondrion, Membrane | 18,8 | 0,94 | 2 | IIMGLGLTAIAGVFGYIAYM | 20 | RYKYESLGYYVAVQENGQEKFIKKKSNWI | 30 | https://www.uniprot.org/uniprot/Q8IPS9 | SubName: Full=Uncharacterized protein, i | Mitochondrion: 0.9487, Endoplasmic reticulum: 0.0205, Peroxisome: 0.0175, Golgi apparatus: 0.0053, Cytoplasm: 0.004, Plastid: 0.002, Nucleus: 0.001, Extracellular: 0.0006, Cell membrane: 0.0004, Lysosome/Vacuole: 0.0001 |
| | Q6IHY5_DROME | Mitochondrion, Membrane | 19 | 0,86 | 3 | ILFGITAGLCGLGIVSFVHLVY | 22 | TMGFAKKKA | 9 | https://www.uniprot.org/uniprot/Q6IHY5 | SubName: Full=HDC00331 {ECO:0000313| | Mitochondrion: 0.7761, Endoplasmic reticulum: 0.1278, Golgi apparatus: 0.0454, Plastid: 0.0213, Cell membrane: 0.0148, Peroxisome: 0.0117, Lysosome/Vacuole: 0.0013, Cytoplasm: 0.0009, Extracellular: 0.0004, Nucleus: 0.0003 |
| | Q7JWT7_DROME | Endoplasmic reticulum, Membrane | 19,5 | 0,85 | 8 | SSIVSAVIFLLVSIFALFGYKLY | 23 | KSLTEKELKKQEKLKSKQQKKAKKSN | 26 | https://www.uniprot.org/uniprot/Q7JWT7 | SubName: Full=AT13707p {ECO:0000313| | Endoplasmic reticulum: 0.8347, Cell membrane: 0.0694, Mitochondrion: 0.0509, Golgi apparatus: 0.031, Peroxisome: 0.01, Lysosome/Vacuole: 0.0017, Plastid: 0.0011, Nucleus: 0.0009, Cytoplasm: 0.0004, Extracellular: 0.0 |
| | Q9W4E0_DROME | Nucleus, Membrane | 19,5 | 0,85 | -1 | LKTVLVGMVQIAVILVLIMAIGS | 23 | PDIIC | 5 | https://www.uniprot.org/uniprot/Q9W4E0 | SubName: Full=GH14578p {ECO:0000313| | Nucleus: 0.675, Golgi apparatus: 0.1227, Cell membrane: 0.0971, Endoplasmic reticulum: 0.0437, Cytoplasm: 0.035, Mitochondrion: 0.0087, Lysosome/Vacuole: 0.0081, Peroxisome: 0.0061, Plastid: 0.0021, Extracellular: 0.0015 |
| | B7Z0M9_DROME | Endoplasmic reticulum, Membrane | 19,7 | 0,90 | 1 | VIKSLIYALIGLAFSLILRWLF | 22 | K | 1 | https://www.uniprot.org/uniprot/B7Z0M9 | RecName: Full=Oxysterol-binding protein | Endoplasmic reticulum: 0.2465, Nucleus: 0.2246, Cell membrane: 0.213, Peroxisome: 0.1442, Golgi apparatus: 0.065, Cytoplasm: 0.05, Mitochondrion: 0.0348, Lysosome/Vacuole: 0.018, Plastid: 0.0035, Extracellular: 0.0003 |
| | Q7KSB3_DROME | Nucleus, Membrane | 19,7 | 0,90 | 1 | VIKSLIYALIGLAFSLILRWLF | 22 | K | 1 | https://www.uniprot.org/uniprot/Q7KSB3 | RecName: Full=Oxysterol-binding protein | Nucleus: 0.2595, Endoplasmic reticulum: 0.2273, Cell membrane: 0.2065, Peroxisome: 0.144, Golgi apparatus: 0.0589, Cytoplasm: 0.051, Mitochondrion: 0.0328, Lysosome/Vacuole: 0.0146, Plastid: 0.005, Extracellular: 0.0003 |
| | BOCKS_DROME | Endoplasmic reticulum, Membrane | 19,8 | 0,99 | 0 | KFYLILVVSVMLATMVYVVL | 20 | TPNA | 4 | https://www.uniprot.org/uniprot/Q709R6 | RecName: Full=LEM domain-containing pr | Endoplasmic reticulum: 0.2862, Nucleus: 0.2727, Golgi apparatus: 0.18, Cytoplasm: 0.0894, Lysosome/Vacuole: 0.0536, Cell membrane: 0.0386, Mitochondrion: 0.0378, Peroxisome: 0.0209, Plastid: 0.0193, Extracellular: 0.0015 |
| | A0A0B4JCZ0_DROME | Nucleus, Membrane | 19,8 | 1,04 | 1 | SLIYALIGLAFSLILRWLF | 19 | K | 1 | https://www.uniprot.org/uniprot/A0A0B4JCZ0 | RecName: Full=Oxysterol-binding protein | Nucleus: 0.279, Cell membrane: 0.1788, Endoplasmic reticulum: 0.1784, Peroxisome: 0.1163, Golgi apparatus: 0.1159, Cytoplasm: 0.0567, Mitochondrion: 0.0335, Plastid: 0.0281, Lysosome/Vacuole: 0.013, Extracellular: 0.0002 |
| | A0A0B4JDD8_DROME | Endoplasmic reticulum, Membrane | 19,8 | 1,04 | 1 | SLIYALIGLAFSLILRWLF | 19 | K | 1 | https://www.uniprot.org/uniprot/A0A0B4JDD8 | RecName: Full=Oxysterol-binding protein | Endoplasmic reticulum: 0.3531, Peroxisome: 0.1794, Cell membrane: 0.1451, Nucleus: 0.1305, Mitochondrion: 0.0741, Golgi apparatus: 0.0688, Cytoplasm: 0.0217, Lysosome/Vacuole: 0.0207, Plastid: 0.0064, Extracellular: 0.0003 |
| | A0A0B4JDG2_DROME | Nucleus, Membrane | 19,8 | 1,04 | 1 | SLIYALIGLAFSLILRWLF | 19 | K | 1 | https://www.uniprot.org/uniprot/A0A0B4JDG2 | RecName: Full=Oxysterol-binding protein | Nucleus: 0.2783, Cell membrane: 0.194, Endoplasmic reticulum: 0.1821, Golgi apparatus: 0.1239, Peroxisome: 0.1205, Cytoplasm: 0.0436, Mitochondrion: 0.0355, Plastid: 0.0122, Lysosome/Vacuole: 0.0095, Extracellular: 0.0003 |
| | Q9I7I8_DROME | Endoplasmic reticulum, Membrane | 19,8 | 1,04 | 1 | SLIYALIGLAFSLILRWLF | 19 | K | 1 | https://www.uniprot.org/uniprot/Q9I7I8 | RecName: Full=Oxysterol-binding protein | Endoplasmic reticulum: 0.3847, Peroxisome: 0.1588, Cell membrane: 0.1547, Nucleus: 0.1094, Golgi apparatus: 0.0783, Mitochondrion: 0.0674, Lysosome/Vacuole: 0.0233, Cytoplasm: 0.0195, Plastid: 0.0036, Extracellular: 0.0002 |
| | M9PHM9_DROME | Mitochondrion, Membrane | 19,9 | 0,95 | -2 | LYPAVAAVALFIVGVIIVMLR | 21 | YGPRLCGLRHHALPDDEDLRGKTYEHEIS | 31 | https://www.uniprot.org/uniprot/M9PHM9 | SubName: Full=Uncharacterized protein, i | Mitochondrion: 0.2764, Extracellular: 0.1604, Golgi apparatus: 0.1354, Plastid: 0.1254, Cell membrane: 0.1189, Lysosome/Vacuole: 0.0884, Endoplasmic reticulum: 0.0879, Cytoplasm: 0.0044, Nucleus: 0.0026, Peroxisome: 0.0003 |
| | Q86BQ0_DROME | Golgi apparatus, Membrane | 19,9 | 1,05 | -1 | LKMMIIMGVIGLVVVGIIA | 19 | KKDEE | 5 | https://www.uniprot.org/uniprot/Q86BQ0 | SubName: Full=GH19566p {ECO:0000313| | Golgi apparatus: 0.2453, Endoplasmic reticulum: 0.2293, Mitochondrion: 0.1749, Nucleus: 0.1056, Cytoplasm: 0.073, Cell membrane: 0.069, Extracellular: 0.0313, Lysosome/Vacuole: 0.0274, Peroxisome: 0.0255, Plastid: 0.0187 |
| | B7Z071_DROME | Cell membrane, Membrane | 20 | 0,80 | 1 | CAKVVTAVVLVALSFMVFLIVYKFV | 25 | RT | 2 | https://www.uniprot.org/uniprot/B7Z071 | SubName: Full=Uncharacterized protein, i | Cell membrane: 0.7005, Endoplasmic reticulum: 0.1023, Nucleus: 0.0929, Peroxisome: 0.0408, Golgi apparatus: 0.02, Cytoplasm: 0.0174, Mitochondrion: 0.0136, Lysosome/Vacuole: 0.008, Plastid: 0.0043, Extracellular: 0.0001 |
| | A0A0B4K7E8_DROME | Cell membrane, Membrane | 20,1 | 0,80 | -1 | MMVVATLTWVMLSCLTLLIYCCNYW | 25 | QILRPHSRRNPEASEAQQSTLLSVED | 26 | https://www.uniprot.org/uniprot/A0A0B4K7E8 | SubName: Full=Uncharacterized protein {E | Cell membrane: 0.8323, Endoplasmic reticulum: 0.1051, Golgi apparatus: 0.0524, Lysosome/Vacuole: 0.0073, Mitochondrion: 0.0019, Nucleus: 0.0007, Plastid: 0.0003, Peroxisome: 0.0001, Cytoplasm: 0.0, Extracellular: 0.0 |
| | O77434_DROME | Golgi apparatus, Membrane | 20,1 | 1,06 | 0 | YVKAMALGMILLVFIMYFW | 19 | VL | 2 | https://www.uniprot.org/uniprot/O77434 | SubName: Full=EG:34F3.8 protein {ECO:00 | Golgi apparatus: 0.6132, Endoplasmic reticulum: 0.2694, Mitochondrion: 0.0382, Lysosome/Vacuole: 0.035, Cell membrane: 0.0178, Nucleus: 0.0132, Peroxisome: 0.0068, Cytoplasm: 0.0035, Plastid: 0.0016, Extracellular: 0.0014 |
| | X2JG21_DROME | Cell membrane, Membrane | 20,3 | 0,88 | -5 | LCVVTALLPLVTLFTCFVTAYVF | 23 | QYDDVHETHCRVSKWAPGDPRVFSGQD | 31 | https://www.uniprot.org/uniprot/X2JG21 | SubName: Full=Uncharacterized protein, i | Cell membrane: 0.367, Endoplasmic reticulum: 0.2727, Lysosome/Vacuole: 0.141, Golgi apparatus: 0.1357, Mitochondrion: 0.0426, Nucleus: 0.0174, Plastid: 0.0131, Cytoplasm: 0.0088, Peroxisome: 0.0011, Extracellular: 0.0006 |
| | Q8INN3_DROME | Cell membrane, Membrane | 20,3 | 0,88 | -1 | ALGLAALLFAVGVYLTLKMVIVA | 23 | SEIG | 4 | https://www.uniprot.org/uniprot/Q8INN3 | SubName: Full=GEO10772p1 {ECO:000031 | Cell membrane: 0.8593, Extracellular: 0.0488, Mitochondrion: 0.035, Lysosome/Vacuole: 0.0189, Golgi apparatus: 0.015, Nucleus: 0.0098, Endoplasmic reticulum: 0.007, Cytoplasm: 0.0049, Plastid: 0.0007, Peroxisome: 0.0007 |
| Dpck | Q9VI57_DROME | Endoplasmic reticulum, Membrane | 20,4 | 0,97 | 1 | WLNRISFLGLFLIVGFTIYML | 21 | LKVFNRLPESWQM | 13 | https://www.uniprot.org/uniprot/Q9VI57 | SubName: Full=Dephospho-CoA kinase {E | Endoplasmic reticulum: 0.6848, Mitochondrion: 0.1186, Peroxisome: 0.0782, Golgi apparatus: 0.0681, Cytoplasm: 0.022, Nucleus: 0.0092, Cell membrane: 0.007, Lysosome/Vacuole: 0.0055, Extracellular: 0.0043, Plastid: 0.0024 |
| | Q9VFY3_DROME | Cell membrane, Membrane | 20,5 | 1,03 | 0 | FLKWFQLGCVLLIFLIWSVL | 20 | SLF | 3 | https://www.uniprot.org/uniprot/Q9VFY3 | SubName: Full=Uncharacterized protein {E | Cell membrane: 0.4141, Endoplasmic reticulum: 0.2865, Golgi apparatus: 0.182, Nucleus: 0.0745, Lysosome/Vacuole: 0.0178, Mitochondrion: 0.0143, Cytoplasm: 0.0056, Peroxisome: 0.0023, Plastid: 0.0017, Extracellular: 0.001 |
| | Q8I935_DROME | Endoplasmic reticulum, Membrane | 20,6 | 0,98 | 4 | NLQYVTLALLFLLFAGFGFLM | 21 | YQQLGQHAATCPKATAYSCAKRK | 23 | https://www.uniprot.org/uniprot/Q8I935 | SubName: Full=Uncharacterized protein, i | Endoplasmic reticulum: 0.4374, Cell membrane: 0.1676, Golgi apparatus: 0.1574, Nucleus: 0.0737, Mitochondrion: 0.0504, Peroxisome: 0.0473, Lysosome/Vacuole: 0.0381, Cytoplasm: 0.0256, Plastid: 0.0023, Extracellular: 0.0003 |
| | B7YZJ3_DROME | Mitochondrion, Membrane | 20,7 | 0,90 | 0 | FELFVLSTVIPAVFVFLWTLISG | 23 | DMKNFKGSKLAYSVYTAKDPINCYQDYEI | 35 | https://www.uniprot.org/uniprot/B7YZJ3 | SubName: Full=Uncharacterized protein {E | Mitochondrion: 0.2918, Endoplasmic reticulum: 0.1981, Extracellular: 0.1978, Golgi apparatus: 0.1707, Lysosome/Vacuole: 0.049, Cell membrane: 0.0376, Cytoplasm: 0.0298, Nucleus: 0.0123, Plastid: 0.0092, Peroxisome: 0.0036 |
| | Q9W0N6_DROME | Golgi apparatus, Membrane | 20,7 | 0,94 | 0 | VVLYGVGVCFVVAVGVSLYLTF | 22 | APSSSSTASS | 10 | https://www.uniprot.org/uniprot/Q9W0N6 | SubName: Full=LD27967p {ECO:0000313| | Golgi apparatus: 0.7151, Mitochondrion: 0.146, Endoplasmic reticulum: 0.0929, Lysosome/Vacuole: 0.02, Cell membrane: 0.0097, Peroxisome: 0.0077, Plastid: 0.0036, Cytoplasm: 0.0024, Nucleus: 0.0023, Extracellular: 0.0001 |
| | Q7K142_DROME | Endoplasmic reticulum, Membrane | 20,7 | 0,99 | 5 | YLLDVVAVYLILLYVLILALR | 21 | RIDFPYEKVRSLYSLMAPFITIAKTKSKKI | 30 | https://www.uniprot.org/uniprot/Q7K142 | RecName: Full=UDP-glucuronosyltransfera | Endoplasmic reticulum: 0.7357, Peroxisome: 0.138, Mitochondrion: 0.0959, Golgi apparatus: 0.0169, Cell membrane: 0.0073, Cytoplasm: 0.0026, Nucleus: 0.0011, Plastid: 0.0011, Lysosome/Vacuole: 0.0009, Extracellular: 0.0006 |
| | S61G2_DROME | Endoplasmic reticulum, Membrane | 20,9 | 0,87 | 0 | IAIATAVGFCIMGFIGFFVKLIHI | 24 | PINNIIVGS | 9 | https://www.uniprot.org/uniprot/Q9VWE9 | RecName: Full=Protein transport protein S | Endoplasmic reticulum: 0.4637, Mitochondrion: 0.2536, Golgi apparatus: 0.2295, Cell membrane: 0.0273, Lysosome/Vacuole: 0.0173, Peroxisome: 0.0032, Extracellular: 0.0028, Cytoplasm: 0.0011, Plastid: 0.001, Nucleus: 0.0006 |
| | X2JCL6_DROME | Endoplasmic reticulum, Membrane | 20,9 | 0,87 | 0 | IAIATAVGFCIMGFIGFFVKLIHI | 24 | PINNIIVGS | 9 | https://www.uniprot.org/uniprot/X2JCL6 | SubName: Full=Sec61 gamma subunit, iso | Endoplasmic reticulum: 0.4637, Mitochondrion: 0.2536, Golgi apparatus: 0.2295, Cell membrane: 0.0273, Lysosome/Vacuole: 0.0173, Peroxisome: 0.0032, Extracellular: 0.0028, Cytoplasm: 0.0011, Plastid: 0.001, Nucleus: 0.0006 |
| | Q9VRR4_DROME | Endoplasmic reticulum, Membrane | 20,9 | 0,95 | 4 | LQYVTLALLFLLFAGFGFLMYQ | 22 | QLGQHAATCPKATAYSCAKRK | 21 | https://www.uniprot.org/uniprot/Q9VRR4 | SubName: Full=LP10490p {ECO:0000313| | Endoplasmic reticulum: 0.255, Nucleus: 0.2078, Golgi apparatus: 0.1652, Cell membrane: 0.1472, Peroxisome: 0.0792, Mitochondrion: 0.0782, Cytoplasm: 0.0339, Lysosome/Vacuole: 0.0152, Plastid: 0.0147, Extracellular: 0.0036 |
| | A1ZBB6_DROME | Endoplasmic reticulum, Membrane | 21,1 | 0,88 | 2 | LWLILRLVIMACSVCALIYHGYFL | 24 | SKIGKMYL | 8 | https://www.uniprot.org/uniprot/A1ZBB6 | SubName: Full=Uncharacterized protein, i | Endoplasmic reticulum: 0.572, Mitochondrion: 0.2063, Peroxisome: 0.1426, Nucleus: 0.0212, Golgi apparatus: 0.0207, Plastid: 0.0156, Cytoplasm: 0.0111, Lysosome/Vacuole: 0.0056, Cell membrane: 0.004, Extracellular: 0.0008 |
| Syx18 | Q9VC58_DROME | Nucleus, Membrane | 21,1 | 0,96 | -1 | LRVWSLFFLLVMSFSLLFLDWY | 22 | YD | 2 | https://www.uniprot.org/uniprot/Q9VC58 | SubName: Full=LD37002p {ECO:0000313| | Nucleus: 0.6936, Endoplasmic reticulum: 0.1734, Golgi apparatus: 0.1073, Cytoplasm: 0.0111, Lysosome/Vacuole: 0.008, Cell membrane: 0.0026, Mitochondrion: 0.0023, Plastid: 0.001, Peroxisome: 0.0006, Extracellular: 0.0001 |
| | Q7K7B0_DROME | Endoplasmic reticulum, Membrane | 21,4 | 1,19 | 9 | LDIYALVLAILIFLVFLT | 18 | RLTVKFLFSKLLGKAKVPARKKKQ | 24 | https://www.uniprot.org/uniprot/Q7K7B0 | SubName: Full=EG:EG0003.4 protein {ECO | Endoplasmic reticulum: 0.7718, Cell membrane: 0.0843, Peroxisome: 0.0762, Golgi apparatus: 0.026, Mitochondrion: 0.0207, Lysosome/Vacuole: 0.01, Nucleus: 0.0036, Cytoplasm: 0.0034, Plastid: 0.0032, Extracellular: 0.001 |
| | A1A6Z0_DROME | Endoplasmic reticulum, Membrane | 21,5 | 1,08 | 2 | VVNVQLLSIAVGIFFVVMLL | 20 | KNNFRSFSVT | 10 | https://www.uniprot.org/uniprot/A1A6Z0 | SubName: Full=GEO11048p1 {ECO:000031 | Endoplasmic reticulum: 0.2158, Cell membrane: 0.1959, Extracellular: 0.1925, Mitochondrion: 0.163, Golgi apparatus: 0.1285, Lysosome/Vacuole: 0.0921, Plastid: 0.0053, Peroxisome: 0.0029, Cytoplasm: 0.0026, Nucleus: 0.0014 |
| | A0A0B4KHQ5_DROME | Endoplasmic reticulum, Membrane | 21,6 | 0,86 | 0 | YNKIILILIIIILEVGILVGLLVLKF | 25 | AHL | 3 | https://www.uniprot.org/uniprot/A0A0B4KHQ5 | SubName: Full=Vesicle transport through i | Endoplasmic reticulum: 0.6447, Golgi apparatus: 0.2614, Mitochondrion: 0.0646, Cell membrane: 0.021, Peroxisome: 0.0054, Nucleus: 0.0013, Lysosome/Vacuole: 0.0008, Plastid: 0.0005, Cytoplasm: 0.0004, Extracellular: 0.0 |
| | Q8IPU7_DROME | Endoplasmic reticulum, Membrane | 21,6 | 0,90 | 2 | ILYSLLVTNVLSLLLGAGFGLWLS | 24 | KRGILLTRVVID | 12 | https://www.uniprot.org/uniprot/Q8IPU7 | SubName: Full=RE54374p {ECO:0000313| | Endoplasmic reticulum: 0.402, Mitochondrion: 0.2482, Peroxisome: 0.1333, Golgi apparatus: 0.104, Nucleus: 0.0501, Cell membrane: 0.03, Lysosome/Vacuole: 0.0141, Cytoplasm: 0.0101, Plastid: 0.0075, Extracellular: 0.0006 |
| | S61G1_DROME | Endoplasmic reticulum, Membrane | 21,6 | 0,90 | 0 | IAIATAVGFAIMGFIGFFVKLIHI | 24 | PINNIIVGS | 9 | https://www.uniprot.org/uniprot/Q9V668 | RecName: Full=Protein transport protein S | Endoplasmic reticulum: 0.4439, Mitochondrion: 0.382, Golgi apparatus: 0.1152, Cell membrane: 0.0307, Lysosome/Vacuole: 0.0179, Peroxisome: 0.0045, Cytoplasm: 0.0018, Plastid: 0.0018, Extracellular: 0.0016, Nucleus: 0.0007 |
| | M9PGI6_DROME | Golgi apparatus, Membrane | 21,6 | 1,08 | 0 | YVKAMALGMILLVFIMYFWV | 20 | L | 1 | https://www.uniprot.org/uniprot/M9PGI6 | SubName: Full=Secretory 22, isoform B {E | Golgi apparatus: 0.6035, Endoplasmic reticulum: 0.2652, Mitochondrion: 0.0432, Lysosome/Vacuole: 0.0387, Cell membrane: 0.0214, Nucleus: 0.0131, Peroxisome: 0.0073, Cytoplasm: 0.0038, Plastid: 0.0022, Extracellular: 0.0016 |
| | C8VV60_DROME | Endoplasmic reticulum, Membrane | 21,8 | 0,87 | 1 | LYMVGIAVALLVLLSSVALHFVLAE | 25 | R | 1 | https://www.uniprot.org/uniprot/C8VV60 | SubName: Full=FI03659p {ECO:0000313|E | Endoplasmic reticulum: 0.4816, Lysosome/Vacuole: 0.1392, Golgi apparatus: 0.1277, Cell membrane: 0.1075, Mitochondrion: 0.0577, Nucleus: 0.0416, Peroxisome: 0.0247, Cytoplasm: 0.0144, Plastid: 0.0051, Extracellular: 0.0006 |
| | ODR4_DROME | Endoplasmic reticulum, Membrane | 21,8 | 0,87 | 1 | LYMVGIAVALLVLLSSVALHFVLAE | 25 | R | 1 | https://www.uniprot.org/uniprot/Q9VTX8 | RecName: Full=Protein odr-4 homolog; | Endoplasmic reticulum: 0.4816, Lysosome/Vacuole: 0.1392, Golgi apparatus: 0.1277, Cell membrane: 0.1075, Mitochondrion: 0.0577, Nucleus: 0.0416, Peroxisome: 0.0247, Cytoplasm: 0.0144, Plastid: 0.0051, Extracellular: 0.0006 |
| | OTE_DROME | Golgi apparatus, Membrane | 22 | 1,16 | 0 | SKLFIVLLVLLLIGVYYIF | 19 | Y | 1 | https://www.uniprot.org/uniprot/P20240 | RecName: Full=Otefin {ECO:0000303|Publ | Golgi apparatus: 0.5049, Endoplasmic reticulum: 0.2298, Nucleus: 0.1582, Cell membrane: 0.0547, Mitochondrion: 0.0172, Lysosome/Vacuole: 0.0118, Cytoplasm: 0.0109, Peroxisome: 0.0087, Plastid: 0.0037, Extracellular: 0.0003 |
| | Q9VEJ4_DROME | Endoplasmic reticulum, Membrane | 22,1 | 0,88 | 0 | WQSVYSNLVIIICFAIFALIVNYVI | 25 | KNLNQE | 6 | https://www.uniprot.org/uniprot/Q9VEJ4 | SubName: Full=HL05730p {ECO:0000313| | Endoplasmic reticulum: 0.5859, Cell membrane: 0.2208, Golgi apparatus: 0.0702, Lysosome/Vacuole: 0.0517, Peroxisome: 0.0234, Mitochondrion: 0.0213, Nucleus: 0.0176, Cytoplasm: 0.0043, Extracellular: 0.0042, Plastid: 0.0006 |
| | Q8IPU8_DROME | Endoplasmic reticulum, Membrane | 22,1 | 0,96 | 2 | ILYSLLVTNVLSLLLGAGFGLWL | 23 | SKRGILLTRVVID | 13 | https://www.uniprot.org/uniprot/Q8IPU8 | SubName: Full=Uncharacterized protein, i | Endoplasmic reticulum: 0.3069, Mitochondrion: 0.3033, Peroxisome: 0.124, Golgi apparatus: 0.0889, Cell membrane: 0.0565, Plastid: 0.0518, Nucleus: 0.0394, Lysosome/Vacuole: 0.019, Cytoplasm: 0.0091, Extracellular: 0.0012 |
| | Q9VPD6_DROME | Endoplasmic reticulum, Membrane | 22,1 | 0,96 | 2 | ILYSLLVTNVLSLLLGAGFGLWL | 23 | SKRGILLTRVVID | 13 | https://www.uniprot.org/uniprot/Q9VPD6 | SubName: Full=RE48077p {ECO:0000313| | Endoplasmic reticulum: 0.464, Mitochondrion: 0.1814, Peroxisome: 0.1475, Golgi apparatus: 0.0814, Nucleus: 0.0473, Cell membrane: 0.0374, Lysosome/Vacuole: 0.0231, Cytoplasm: 0.0142, Plastid: 0.0035, Extracellular: 0.0003 |
| | Q0E8W6_DROME | Cell membrane, Membrane | 22,2 | 1,17 | 2 | SGFSVAMIFLIVAMFMYIV | 19 | NPSTWGNFTSNKRAGGRRDNNQDGSPI | 35 | https://www.uniprot.org/uniprot/Q0E8W6 | SubName: Full=IP15630p {ECO:0000313|E | Cell membrane: 0.4341, Endoplasmic reticulum: 0.2981, Golgi apparatus: 0.1168, Lysosome/Vacuole: 0.108, Extracellular: 0.0381, Mitochondrion: 0.0032, Plastid: 0.0011, Nucleus: 0.0003, Cytoplasm: 0.0001, Peroxisome: 0.0001 |
| | SMCO4_DROME | Endoplasmic reticulum, Membrane | 22,3 | 1,01 | 2 | IRTVVLPALGVLAVFLVLFVYL | 22 | KTRPAVLA | 8 | https://www.uniprot.org/uniprot/Q8SY72 | RecName: Full=Single-pass membrane and | Endoplasmic reticulum: 0.3817, Golgi apparatus: 0.3341, Mitochondrion: 0.1716, Cell membrane: 0.0749, Nucleus: 0.0193, Peroxisome: 0.0081, Extracellular: 0.0056, Lysosome/Vacuole: 0.0031, Cytoplasm: 0.0015, Plastid: 0.0002 |
| | A0A1Z1CK50_DROME | Endoplasmic reticulum, Membrane | 22,4 | 0,66 | 0 | IFLTPSLHQRFINFVIFLFMLAMYLHAAVTPSL | 34 | L | 1 | https://www.uniprot.org/uniprot/A0A1Z1CK50 | SubName: Full=Uncharacterized protein {E | Endoplasmic reticulum: 0.4305, Nucleus: 0.2175, Golgi apparatus: 0.1069, Mitochondrion: 0.0751, Peroxisome: 0.0587, Cytoplasm: 0.0446, Cell membrane: 0.044, Lysosome/Vacuole: 0.0152, Extracellular: 0.0073, Plastid: 0.0003 |
| | E1JI87_DROME | Cell membrane, Membrane | 22,5 | 0,87 | 3 | AVTVTVVFVTLTFLLGIVLATTTNLF | 26 | GKTLNQRFLTQLISLNQRMILKSMWMEN | 28 | https://www.uniprot.org/uniprot/E1JI87 | SubName: Full=Peptidoglycan recognition | Cell membrane: 0.2328, Endoplasmic reticulum: 0.1826, Peroxisome: 0.1541, Nucleus: 0.1531, Mitochondrion: 0.0922, Plastid: 0.0708, Golgi apparatus: 0.0645, Lysosome/Vacuole: 0.0322, Cytoplasm: 0.0144, Extracellular: 0.0032 |
| | B7YZW3_DROME | Nucleus, Membrane | 22,6 | 1,03 | 1 | AVCIVVSFLVGTFLVAMLVNML | 22 | RG | 2 | https://www.uniprot.org/uniprot/B7YZW3 | SubName: Full=Uncharacterized protein, i | Nucleus: 0.5736, Cell membrane: 0.2174, Cytoplasm: 0.143, Endoplasmic reticulum: 0.0277, Golgi apparatus: 0.0266, Peroxisome: 0.0048, Lysosome/Vacuole: 0.0044, Mitochondrion: 0.0016, Plastid: 0.0004, Extracellular: 0.0003 |
| | Q9VJJ8_DROME | Cell membrane, Membrane | 22,6 | 1,03 | 1 | AVCIVVSFLVGTFLVAMLVNML | 22 | RG | 2 | https://www.uniprot.org/uniprot/Q9VJJ8 | SubName: Full=LD30602p {ECO:0000313| | Cell membrane: 0.55, Nucleus: 0.2005, Golgi apparatus: 0.0686, Cytoplasm: 0.068, Endoplasmic reticulum: 0.0499, Peroxisome: 0.0247, Lysosome/Vacuole: 0.0203, Mitochondrion: 0.016, Extracellular: 0.0015, Plastid: 0.0006 |
| | A0A0B4JCZ4_DROME | Endoplasmic reticulum, Membrane | 22,7 | 0,84 | 1 | IMILICLTVLGILAASYVSSYFIIQAS | 27 | K | 1 | https://www.uniprot.org/uniprot/A0A0B4JCZ4 | SubName: Full=Syntaxin 1A, isoform B {E | Endoplasmic reticulum: 0.5712, Golgi apparatus: 0.2527, Mitochondrion: 0.1051, Nucleus: 0.0359, Peroxisome: 0.0147, Lysosome/Vacuole: 0.007, Plastid: 0.0055, Cytoplasm: 0.0045, Cell membrane: 0.0035, Extracellular: 0.0 |
| CG17580 | Q7JW78_DROME | Cell membrane, Membrane | 22,7 | 0,99 | -2 | RLVYWLGVLIMVIGIGVGMYFTL | 23 | KSDFGECSSYDVRCD | 15 | https://www.uniprot.org/uniprot/Q7JW78 | SubName: Full=AT18054p {ECO:0000313| | Cell membrane: 0.4915, Extracellular: 0.2284, Mitochondrion: 0.1049, Golgi apparatus: 0.0936, Lysosome/Vacuole: 0.0471, Endoplasmic reticulum: 0.0295, Nucleus: 0.0035, Cytoplasm: 0.0007, Plastid: 0.0005, Peroxisome: 0.0004 |
| | Q8I0D5_DROME | Endoplasmic reticulum, Membrane | 22,7 | 1,08 | 4 | LQYVTLALLFLLFAGFGFLMY | 21 | QQLGQHAATCPKATAYSCAKRK | 22 | https://www.uniprot.org/uniprot/Q8I0D5 | SubName: Full=GH09028p {ECO:0000313| | Endoplasmic reticulum: 0.3017, Nucleus: 0.2103, Golgi apparatus: 0.143, Cell membrane: 0.1227, Peroxisome: 0.0851, Mitochondrion: 0.0614, Cytoplasm: 0.0396, Lysosome/Vacuole: 0.0184, Plastid: 0.0143, Extracellular: 0.0035 |
| | Q8I934_DROME | Endoplasmic reticulum, Membrane | 22,7 | 1,08 | 4 | LQYVTLALLFLLFAGFGFLMY | 21 | QQLGQHAATCPKATAYSCAKRK | 22 | https://www.uniprot.org/uniprot/Q8I934 | SubName: Full=Uncharacterized protein, i | Endoplasmic reticulum: 0.2973, Cell membrane: 0.1993, Golgi apparatus: 0.154, Nucleus: 0.1327, Mitochondrion: 0.0722, Peroxisome: 0.0621, Plastid: 0.0324, Cytoplasm: 0.0244, Lysosome/Vacuole: 0.0181, Extracellular: 0.0076 |
| | A8DZ13_DROME | Golgi apparatus, Membrane | 22,7 | 1,08 | 0 | FGFWCGLVGLAVFIIFLAYFH | 21 | INRERIDPTS | 10 | https://www.uniprot.org/uniprot/A8DZ13 | SubName: Full=GEO11020p1 {ECO:000031 | Golgi apparatus: 0.2872, Endoplasmic reticulum: 0.2598, Cell membrane: 0.1837, Mitochondrion: 0.1473, Lysosome/Vacuole: 0.0542, Extracellular: 0.0419, Plastid: 0.0242, Nucleus: 0.0006, Peroxisome: 0.0006, Cytoplasm: 0.0003 |
| | A0A0B4K675_DROME | Endoplasmic reticulum, Membrane | 22,8 | 1,04 | -3 | VAIFFGLLSTCLIVALVIVAVH | 22 | RDNLLLQLDEAKRMLDVLEEYIQSHLLSTI | 35 | https://www.uniprot.org/uniprot/A0A0B4K675 | SubName: Full=Uncharacterized protein {E | Endoplasmic reticulum: 0.2605, Mitochondrion: 0.2508, Golgi apparatus: 0.2469, Nucleus: 0.074, Lysosome/Vacuole: 0.0421, Cytoplasm: 0.0383, Cell membrane: 0.0348, Extracellular: 0.0198, Peroxisome: 0.0197, Plastid: 0.013 |
| | A8JRE2_DROME | Endoplasmic reticulum, Membrane | 22,9 | 1,00 | 1 | LINGFFLVMLLLVLFSSFMYLPN | 23 | YMQTLMQFVTPPNWHNSPDSAAYVAQ | 31 | https://www.uniprot.org/uniprot/A8JRE2 | SubName: Full=MIP14036p {ECO:0000313 | Endoplasmic reticulum: 0.3874, Golgi apparatus: 0.2631, Mitochondrion: 0.2507, Extracellular: 0.0313, Lysosome/Vacuole: 0.0237, Cell membrane: 0.0175, Peroxisome: 0.0157, Cytoplasm: 0.0061, Nucleus: 0.0041, Plastid: 0.0002 |
| | GOSR2_DROME | Golgi apparatus, Membrane | 23,2 | 1,16 | 0 | RIFIGGVVVTLLIIALIIYF | 20 | LVL | 3 | https://www.uniprot.org/uniprot/Q9VRL2 | RecName: Full=Probable Golgi SNAP recep | Golgi apparatus: 0.7594, Endoplasmic reticulum: 0.1544, Cell membrane: 0.038, Lysosome/Vacuole: 0.03, Mitochondrion: 0.008, Nucleus: 0.0067, Plastid: 0.0014, Peroxisome: 0.0014, Cytoplasm: 0.0006, Extracellular: 0.0001 |
| CG43210 | A0A0B4K684_DROME | Endoplasmic reticulum, Membrane | 23,3 | 0,97 | 1 | IHLSYFICGPVFILMVFMYFYYWY | 24 | LNVRECRRSRGREATPTLPDDISVERLQV | 30 | https://www.uniprot.org/uniprot/A0A0B4K684 | SubName: Full=Uncharacterized protein, i | Endoplasmic reticulum: 0.451, Golgi apparatus: 0.2716, Cell membrane: 0.1506, Lysosome/Vacuole: 0.0379, Extracellular: 0.0363, Mitochondrion: 0.0325, Plastid: 0.0094, Nucleus: 0.005, Cytoplasm: 0.0036, Peroxisome: 0.002 |
| | GOGA5_DROME | Golgi apparatus, Membrane | 23,3 | 1,01 | 0 | MMRVSVIVYVALLHLWVMFVLLS | 23 | TTPN | 4 | https://www.uniprot.org/uniprot/Q8SZ63 | RecName: Full=Golgin-84; | Golgi apparatus: 0.9828, Endoplasmic reticulum: 0.0143, Mitochondrion: 0.001, Lysosome/Vacuole: 0.0009, Nucleus: 0.0005, Cell membrane: 0.0004, Cytoplasm: 0.0001, Extracellular: 0.0, Plastid: 0.0, Peroxisome: 0.0 |
| | R9UAE0_DROME | Endoplasmic reticulum, Membrane | 23,3 | 1,06 | 2 | FVGLCVLYNFCMLLIVLSIYIY | 22 | KRWN | 4 | https://www.uniprot.org/uniprot/R9UAE0 | SubName: Full=MIP36958p1 {ECO:000031 | Endoplasmic reticulum: 0.8236, Cell membrane: 0.0957, Golgi apparatus: 0.0528, Lysosome/Vacuole: 0.0121, Mitochondrion: 0.0102, Nucleus: 0.0026, Plastid: 0.0018, Peroxisome: 0.0008, Cytoplasm: 0.0002, Extracellular: 0.0001 |
| | Q0E985_DROME | Endoplasmic reticulum, Membrane | 23,4 | 1,02 | 0 | FYMFAFCFVCATILVAFILSRFI | 23 | TIKPVDF | 7 | https://www.uniprot.org/uniprot/Q0E985 | SubName: Full=RH74701p {ECO:0000313| | Endoplasmic reticulum: 0.5982, Golgi apparatus: 0.2748, Mitochondrion: 0.0579, Peroxisome: 0.0196, Cell membrane: 0.0194, Lysosome/Vacuole: 0.0132, Nucleus: 0.011, Cytoplasm: 0.0031, Extracellular: 0.0022, Plastid: 0.0007 |
| | Q9W1U8_DROME | Cell membrane, Membrane | 23,6 | 1,03 | 3 | LLLGTSTLVLCLIISFLMFAILQ | 23 | WKASRLRGSKSVQGGPCES | 19 | https://www.uniprot.org/uniprot/Q9W1U8 | SubName: Full=IP06655p {ECO:0000313|E | Cell membrane: 0.5029, Endoplasmic reticulum: 0.2818, Golgi apparatus: 0.1052, Lysosome/Vacuole: 0.0464, Peroxisome: 0.0369, Mitochondrion: 0.0178, Nucleus: 0.0037, Plastid: 0.003, Cytoplasm: 0.0021, Extracellular: 0.0002 |
| | GOSR1_DROME | Golgi apparatus, Membrane | 23,7 | 1,19 | 0 | SLILGAVIGFCVILLLLYAF | 20 | N | 1 | https://www.uniprot.org/uniprot/Q9VE50 | RecName: Full=Golgi SNAP receptor comp | Golgi apparatus: 0.8638, Endoplasmic reticulum: 0.0758, Lysosome/Vacuole: 0.0215, Mitochondrion: 0.0172, Cell membrane: 0.0141, Nucleus: 0.004, Peroxisome: 0.0019, Cytoplasm: 0.0014, Plastid: 0.0002, Extracellular: 0.0001 |
| | Q9VE99_DROME | Endoplasmic reticulum, Membrane | 23,8 | 0,95 | 1 | FLSIAFGGALLVIITVSVYAFYSLY | 25 | HAILKKFPSRHEEL | 14 | https://www.uniprot.org/uniprot/Q9VE99 | SubName: Full=GEO12060p1 {ECO:000031 | Endoplasmic reticulum: 0.4471, Mitochondrion: 0.3375, Cell membrane: 0.0875, Plastid: 0.0452, Golgi apparatus: 0.0383, Peroxisome: 0.0276, Lysosome/Vacuole: 0.0128, Nucleus: 0.0033, Cytoplasm: 0.0006, Extracellular: 0.0001 |
| | M9NE87_DROME | Endoplasmic reticulum, Membrane | 23,8 | 0,99 | 0 | SLYMVGIAVALLVLLSSVALHFVL | 24 | AER | 3 | https://www.uniprot.org/uniprot/M9NE87 | SubName: Full=Uncharacterized protein, i | Endoplasmic reticulum: 0.4835, Golgi apparatus: 0.1249, Cell membrane: 0.1244, Lysosome/Vacuole: 0.1082, Mitochondrion: 0.0615, Nucleus: 0.0516, Peroxisome: 0.0259, Cytoplasm: 0.0151, Plastid: 0.0042, Extracellular: 0.0006 |
| | Q9VNI9_DROME | Endoplasmic reticulum, Membrane | 23,9 | 1,33 | 2 | MLLFFAFSLFLACVFYIV | 18 | QKRLF | 5 | https://www.uniprot.org/uniprot/Q9VNI9 | SubName: Full=IP18173p {ECO:0000313|E | Endoplasmic reticulum: 0.6613, Golgi apparatus: 0.2329, Cell membrane: 0.0371, Lysosome/Vacuole: 0.0291, Mitochondrion: 0.0185, Peroxisome: 0.0107, Nucleus: 0.0069, Cytoplasm: 0.0025, Plastid: 0.0009, Extracellular: 0.0 |
| | M9MRI9_DROME | Golgi apparatus, Membrane | 24 | 1,20 | 0 | FFNIFLILFSVLFCVLHFVA | 20 | S | 1 | https://www.uniprot.org/uniprot/M9MRI9 | SubName: Full=Quick-to-court, isoform H | Golgi apparatus: 0.7601, Endoplasmic reticulum: 0.1479, Nucleus: 0.0377, Cytoplasm: 0.0195, Cell membrane: 0.0108, Mitochondrion: 0.0085, Lysosome/Vacuole: 0.0083, Plastid: 0.007, Peroxisome: 0.0001, Extracellular: 0.0 |
| | A8WH76_DROME | Endoplasmic reticulum, Membrane | 24,2 | 0,97 | 1 | LLTLLLYMGGVSGMGLTLAVYYLFI | 25 | WDSRMPPLPVFKHTHPIG | 18 | https://www.uniprot.org/uniprot/A8WH76 | SubName: Full=GEO10024p1 {ECO:000031 | Endoplasmic reticulum: 0.4884, Mitochondrion: 0.3697, Golgi apparatus: 0.0637, Peroxisome: 0.0312, Cell membrane: 0.0162, Extracellular: 0.0082, Nucleus: 0.0065, Lysosome/Vacuole: 0.0062, Plastid: 0.0059, Cytoplasm: 0.004 |
| | Q8IQC4_DROME | Endoplasmic reticulum, Membrane | 24,3 | 1,01 | 0 | ITIHVIFMVLFCVFWYFNYSVLLS | 24 | LAGLIGPTSASIADIIPGVQGHGLKVT | 27 | https://www.uniprot.org/uniprot/Q8IQC4 | SubName: Full=Ghiberti {ECO:0000313|EM | Endoplasmic reticulum: 0.6569, Golgi apparatus: 0.2958, Cell membrane: 0.024, Lysosome/Vacuole: 0.0111, Mitochondrion: 0.0064, Extracellular: 0.005, Peroxisome: 0.0005, Plastid: 0.0002, Nucleus: 0.0001, Cytoplasm: 0.0001 |
| | Q4QPY0_DROME | Cell membrane, Membrane | 24,3 | 1,10 | 0 | LFGIMMVPLMVLSLGPFYLLLL | 22 | YW | 2 | https://www.uniprot.org/uniprot/Q4QPY0 | SubName: Full=GEO12067p1 {ECO:000031 | Cell membrane: 0.4061, Endoplasmic reticulum: 0.2963, Golgi apparatus: 0.2095, Lysosome/Vacuole: 0.0592, Mitochondrion: 0.0136, Nucleus: 0.0085, Plastid: 0.0037, Peroxisome: 0.0022, Cytoplasm: 0.0007, Extracellular: 0.0003 |
| | A8DYH6_DROME | Golgi apparatus, Membrane | 24,4 | 0,84 | -2 | FLNVLLGSFVSLILYASFAFVPGYCVTVF | 29 | QLLWPQTSVQNLTSVCSTSTEGFCGNES | 33 | https://www.uniprot.org/uniprot/A8DYH6 | SubName: Full=FI16908p1 {ECO:0000313| | Golgi apparatus: 0.358, Cell membrane: 0.2251, Lysosome/Vacuole: 0.2226, Endoplasmic reticulum: 0.1495, Mitochondrion: 0.0258, Extracellular: 0.0083, Nucleus: 0.0034, Peroxisome: 0.0028, Cytoplasm: 0.0026, Plastid: 0.0019 |
| Syx13 | Q9VU45_DROME | Golgi apparatus, Membrane | 25 | 1,14 | 0 | ILILLVIAVIIGLIVTGVIVAK | 22 | LNS | 3 | https://www.uniprot.org/uniprot/Q9VU45 | SubName: Full=LD27581p {ECO:0000313| | Golgi apparatus: 0.6177, Endoplasmic reticulum: 0.2814, Mitochondrion: 0.0374, Lysosome/Vacuole: 0.0251, Cell membrane: 0.0235, Nucleus: 0.0097, Peroxisome: 0.0027, Cytoplasm: 0.0016, Plastid: 0.0009, Extracellular: 0.0 |
| | Q4V477_DROME | Cell membrane, Membrane | 25,1 | 1,14 | 1 | TLYLIAACFALLLLMIIYMIHV | 22 | ARRYNHVRAYGSSACGSTLFYSYSDCDVF | 30 | https://www.uniprot.org/uniprot/Q4V477 | SubName: Full=IP03610p {ECO:0000313|E | Cell membrane: 0.5818, Endoplasmic reticulum: 0.1139, Lysosome/Vacuole: 0.1076, Extracellular: 0.1057, Golgi apparatus: 0.085, Mitochondrion: 0.0026, Nucleus: 0.0022, Plastid: 0.0006, Cytoplasm: 0.0004, Peroxisome: 0.0004 |
| | Q9VX52_DROME | Golgi apparatus, Membrane | 25,4 | 1,06 | 3 | IMWVSIGMGVTIVLLITIALCYIA | 24 | REKCQKRQREYYVTA | 15 | https://www.uniprot.org/uniprot/Q9VX52 | SubName: Full=IP03239p {ECO:0000313|E | Golgi apparatus: 0.39, Mitochondrion: 0.2529, Cell membrane: 0.2197, Extracellular: 0.0794, Endoplasmic reticulum: 0.0393, Lysosome/Vacuole: 0.0087, Plastid: 0.0043, Nucleus: 0.003, Peroxisome: 0.0017, Cytoplasm: 0.0011 |
| | Q9VV76_DROME | Endoplasmic reticulum, Membrane | 25,4 | 1,41 | 0 | GYWLVIIALFVAIIVVVF | 18 | V | 1 | https://www.uniprot.org/uniprot/Q9VV76 | SubName: Full=Syntaxin 8 {ECO:0000313| | Endoplasmic reticulum: 0.4015, Golgi apparatus: 0.2909, Lysosome/Vacuole: 0.1575, Cell membrane: 0.1185, Mitochondrion: 0.0116, Nucleus: 0.0087, Cytoplasm: 0.0052, Plastid: 0.0033, Peroxisome: 0.0025, Extracellular: 0.0001 |
| | Q9VPA4_DROME | Endoplasmic reticulum, Membrane | 25,6 | 1,22 | 1 | LAYVFMSLLVLIVCFILVNVV | 21 | HKLYRSLNGEVSTGLVSPAQQLKTSIMELI | 35 | https://www.uniprot.org/uniprot/Q9VPA4 | SubName: Full=MIP08452p {ECO:0000313 | Endoplasmic reticulum: 0.4019, Golgi apparatus: 0.2716, Cell membrane: 0.125, Mitochondrion: 0.0869, Lysosome/Vacuole: 0.0487, Plastid: 0.0266, Extracellular: 0.0264, Nucleus: 0.0049, Cytoplasm: 0.004, Peroxisome: 0.0038 |
| | Q8T8Y5_DROME | Endoplasmic reticulum, Membrane | 25,7 | 0,99 | 2 | PLEWTTLVIGVVFGLILVFMILRFIF | 26 | NLIVPKIYQRFTNS | 14 | https://www.uniprot.org/uniprot/Q8T8Y5 | SubName: Full=AT16536p {ECO:0000313| | Endoplasmic reticulum: 0.4064, Peroxisome: 0.2595, Plastid: 0.1152, Mitochondrion: 0.1075, Cell membrane: 0.048, Golgi apparatus: 0.0242, Nucleus: 0.0204, Lysosome/Vacuole: 0.0103, Cytoplasm: 0.0083, Extracellular: 0.0004 |
| | M9MRN1_DROME | Cell membrane, Membrane | 25,8 | 1,36 | 0 | LLTLCLIVIGVIIIIAIAL | 19 | T | 1 | https://www.uniprot.org/uniprot/M9MRN1 | SubName: Full=Supervillin, isoform AG {E | Cell membrane: 0.3719, Nucleus: 0.2665, Cytoplasm: 0.1188, Golgi apparatus: 0.0938, Endoplasmic reticulum: 0.0797, Lysosome/Vacuole: 0.0479, Plastid: 0.0098, Mitochondrion: 0.0097, Peroxisome: 0.0016, Extracellular: 0.0004 |
| | Q9W018_DROME | Cell membrane, Membrane | 25,8 | 1,36 | 0 | LLTLCLIVIGVIIIIAIAL | 19 | T | 1 | https://www.uniprot.org/uniprot/Q9W018 | SubName: Full=Supervillin, isoform AF {E | Cell membrane: 0.3312, Nucleus: 0.2933, Cytoplasm: 0.1453, Golgi apparatus: 0.0861, Endoplasmic reticulum: 0.0733, Lysosome/Vacuole: 0.0489, Plastid: 0.0108, Mitochondrion: 0.0088, Peroxisome: 0.0017, Extracellular: 0.0006 |
| | Q6IIF2_DROME | Endoplasmic reticulum, Membrane | 26 | 1,04 | 3 | YFVIVIYLCGLCSLGFFLSIYHIFF | 25 | WDSRMPPVYKGQKKGPAFG | 19 | https://www.uniprot.org/uniprot/Q6IIF2 | SubName: Full=GEO11103p1 {ECO:000031 | Endoplasmic reticulum: 0.6026, Plastid: 0.101, Cell membrane: 0.0827, Mitochondrion: 0.0668, Golgi apparatus: 0.0604, Peroxisome: 0.0559, Lysosome/Vacuole: 0.0158, Nucleus: 0.0086, Cytoplasm: 0.0049, Extracellular: 0.0012 |
| | USE1_DROME | Endoplasmic reticulum, Membrane | 26 | 1,13 | 4 | CWLWLMIAFVIATFIGMVLFMKI | 23 | MKKKKS | 6 | https://www.uniprot.org/uniprot/Q9VSU7 | RecName: Full=Vesicle transport protein U | Endoplasmic reticulum: 0.8383, Golgi apparatus: 0.0604, Cell membrane: 0.0436, Mitochondrion: 0.033, Peroxisome: 0.0172, Nucleus: 0.0031, Lysosome/Vacuole: 0.0031, Plastid: 0.0008, Cytoplasm: 0.0005, Extracellular: 0.0 |
| | Q7JYX5_DROME | Golgi apparatus, Membrane | 26,2 | 1,01 | 0 | VYVVVGLVITFIVYVIVSMACGGLAW | 26 | QSCV | 4 | https://www.uniprot.org/uniprot/Q7JYX5 | SubName: Full=RH15778p {ECO:0000313| | Golgi apparatus: 0.5155, Endoplasmic reticulum: 0.3051, Mitochondrion: 0.0868, Lysosome/Vacuole: 0.0394, Cell membrane: 0.0285, Peroxisome: 0.0147, Cytoplasm: 0.006, Nucleus: 0.0036, Plastid: 0.0003, Extracellular: 0.0002 |
| | ZFPL1_DROME | Golgi apparatus, Membrane | 26,6 | 1,21 | -3 | TWFLVTAGILAFVLFVYLMAWL | 22 | GRGGSDAVDEGWNNPNPQPNHYE | 23 | https://www.uniprot.org/uniprot/Q9VD26 | RecName: Full=Zinc finger protein-like 1 h | Golgi apparatus: 0.5501, Cell membrane: 0.3297, Endoplasmic reticulum: 0.0661, Mitochondrion: 0.0281, Lysosome/Vacuole: 0.0105, Nucleus: 0.0098, Extracellular: 0.0026, Peroxisome: 0.0018, Cytoplasm: 0.0012, Plastid: 0.0 |
| | Q7JR34_DROME | Cell membrane, Membrane | 26,7 | 0,99 | -4 | VIGIFAGFGILLLVASPLLLLAAPCII | 27 | CCKCRGCTGSKIDEVDAELEEEVTALQG | 28 | https://www.uniprot.org/uniprot/Q7JR34 | RecName: Full=RBR-type E3 ubiquitin tran | Cell membrane: 0.5476, Nucleus: 0.2459, Golgi apparatus: 0.0868, Cytoplasm: 0.0856, Endoplasmic reticulum: 0.0149, Mitochondrion: 0.0126, Extracellular: 0.0021, Lysosome/Vacuole: 0.0019, Plastid: 0.0018, Peroxisome: 0.0007 |
| | A8JUT0_DROME | Endoplasmic reticulum, Membrane | 27 | 1,08 | -1 | IISIFLYLGGISGLGMVLALYYLMF | 25 | FDSSMPDIHLKFPVSIGGHPVQKVHEYQ | 28 | https://www.uniprot.org/uniprot/A8JUT0 | SubName: Full=InaF-A {ECO:0000313|EM | Endoplasmic reticulum: 0.7218, Golgi apparatus: 0.1566, Mitochondrion: 0.0734, Cell membrane: 0.0252, Lysosome/Vacuole: 0.0124, Peroxisome: 0.0047, Nucleus: 0.0033, Cytoplasm: 0.0012, Plastid: 0.0008, Extracellular: 0.0005 |
| | A1Z8I8_DROME | Endoplasmic reticulum, Membrane | 27,1 | 1,18 | 8 | YLVFFVMLALILIVFALFGYKLY | 23 | KSLTEKELKKQEKLKSKQQKKAKKSN | 26 | https://www.uniprot.org/uniprot/A1Z8I8 | SubName: Full=Uncharacterized protein, i | Endoplasmic reticulum: 0.8413, Cell membrane: 0.1003, Golgi apparatus: 0.0273, Mitochondrion: 0.0145, Lysosome/Vacuole: 0.0077, Peroxisome: 0.0047, Nucleus: 0.0029, Plastid: 0.0008, Cytoplasm: 0.0005, Extracellular: 0.0 |
| | M9ND64_DROME | Nucleus, Membrane | 27,1 | 1,29 | 0 | ISALVILFIAITLAVLVFLLT | 21 | GRQVHTHHWDYFNPPGNDHGRQT | 23 | https://www.uniprot.org/uniprot/M9ND64 | SubName: Full=Uncharacterized protein, i | Nucleus: 0.8828, Cell membrane: 0.0631, Golgi apparatus: 0.0219, Cytoplasm: 0.021, Endoplasmic reticulum: 0.0102, Lysosome/Vacuole: 0.0007, Plastid: 0.0002, Mitochondrion: 0.0001, Peroxisome: 0.0001, Extracellular: 0.0 |
| | M9NF47_DROME | Nucleus, Membrane | 27,1 | 1,29 | 0 | ISALVILFIAITLAVLVFLLT | 21 | GRQVHTHHWDYFNPPGNDHGRQT | 23 | https://www.uniprot.org/uniprot/M9NF47 | SubName: Full=Uncharacterized protein, i | Nucleus: 0.8366, Cytoplasm: 0.1046, Golgi apparatus: 0.0333, Cell membrane: 0.0119, Endoplasmic reticulum: 0.0111, Lysosome/Vacuole: 0.0016, Plastid: 0.0005, Mitochondrion: 0.0003, Peroxisome: 0.0002, Extracellular: 0.0 |
| | M9NFP8_DROME | Nucleus, Membrane | 27,1 | 1,29 | 0 | ISALVILFIAITLAVLVFLLT | 21 | GRQVHTHHWDYFNPPGNDHGRQT | 23 | https://www.uniprot.org/uniprot/M9NFP8 | SubName: Full=Uncharacterized protein, i | Nucleus: 0.8216, Cytoplasm: 0.0888, Golgi apparatus: 0.046, Cell membrane: 0.0247, Endoplasmic reticulum: 0.0156, Lysosome/Vacuole: 0.0017, Mitochondrion: 0.0007, Plastid: 0.0006, Peroxisome: 0.0002, Extracellular: 0.0001 |
| | M9PBX1_DROME | Nucleus, Membrane | 27,1 | 1,29 | 0 | ISALVILFIAITLAVLVFLLT | 21 | GRQVHTHHWDYFNPPGNDHGRQT | 23 | https://www.uniprot.org/uniprot/M9PBX1 | SubName: Full=Uncharacterized protein, i | Nucleus: 0.8274, Cytoplasm: 0.1043, Golgi apparatus: 0.0418, Endoplasmic reticulum: 0.0128, Cell membrane: 0.0107, Lysosome/Vacuole: 0.0021, Plastid: 0.0005, Mitochondrion: 0.0002, Peroxisome: 0.0001, Extracellular: 0.0 |
| | M9PET1_DROME | Nucleus, Membrane | 27,1 | 1,29 | 0 | ISALVILFIAITLAVLVFLLT | 21 | GRQVHTHHWDYFNPPGNDHGRQT | 23 | https://www.uniprot.org/uniprot/M9PET1 | SubName: Full=Uncharacterized protein, i | Nucleus: 0.8042, Cytoplasm: 0.086, Golgi apparatus: 0.0676, Cell membrane: 0.0214, Endoplasmic reticulum: 0.0175, Lysosome/Vacuole: 0.0021, Plastid: 0.0006, Mitochondrion: 0.0004, Peroxisome: 0.0002, Extracellular: 0.0001 |
| | M9PHT9_DROME | Nucleus, Membrane | 27,1 | 1,29 | 0 | ISALVILFIAITLAVLVFLLT | 21 | GRQVHTHHWDYFNPPGNDHGRQT | 23 | https://www.uniprot.org/uniprot/M9PHT9 | SubName: Full=Uncharacterized protein, i | Nucleus: 0.8365, Cytoplasm: 0.105, Golgi apparatus: 0.033, Cell membrane: 0.0119, Endoplasmic reticulum: 0.0112, Lysosome/Vacuole: 0.0015, Plastid: 0.0005, Mitochondrion: 0.0003, Peroxisome: 0.0002, Extracellular: 0.0 |
| | A1Z763_DROME | Nucleus, Membrane | 27,2 | 1,18 | 3 | FITTFLSLVFFLLTVMVVSLVLT | 23 | RGKRMFGSMI | 10 | https://www.uniprot.org/uniprot/A1Z763 | SubName: Full=MIP13269p {ECO:0000313 | Nucleus: 0.2568, Endoplasmic reticulum: 0.2408, Golgi apparatus: 0.2328, Cell membrane: 0.1846, Peroxisome: 0.0305, Mitochondrion: 0.0209, Cytoplasm: 0.0161, Lysosome/Vacuole: 0.01, Plastid: 0.0071, Extracellular: 0.0004 |
| | Q7K0X9_DROME | Golgi apparatus, Membrane | 27,2 | 1,30 | 1 | LILVGILSAVLLAIILILVFQ | 21 | FKN | 3 | https://www.uniprot.org/uniprot/Q7K0X9 | SubName: Full=LD23667p {ECO:0000313| | Golgi apparatus: 0.6731, Endoplasmic reticulum: 0.2648, Lysosome/Vacuole: 0.029, Nucleus: 0.012, Cell membrane: 0.0094, Mitochondrion: 0.0079, Peroxisome: 0.0019, Cytoplasm: 0.0014, Plastid: 0.0004, Extracellular: 0.0001 |
| | Q8IRS2_DROME | Cell membrane, Membrane | 27,2 | 1,60 | 1 | AMVYIIFIFIIMFIYLF | 17 | K | 1 | https://www.uniprot.org/uniprot/Q8IRS2 | SubName: Full=IP13349p {ECO:0000313|E | Cell membrane: 0.3854, Nucleus: 0.2977, Golgi apparatus: 0.1266, Endoplasmic reticulum: 0.0973, Cytoplasm: 0.0572, Extracellular: 0.0114, Mitochondrion: 0.0085, Lysosome/Vacuole: 0.0074, Plastid: 0.0068, Peroxisome: 0.0018 |
| | Q6AWN6_DROME | Endoplasmic reticulum, Membrane | 27,3 | 1,37 | 0 | LMLLVLIFNVALALLFVYML | 20 | T | 1 | https://www.uniprot.org/uniprot/Q6AWN6 | SubName: Full=GEO12823p1 {ECO:000031 | Endoplasmic reticulum: 0.5636, Golgi apparatus: 0.3396, Cell membrane: 0.0433, Lysosome/Vacuole: 0.0384, Nucleus: 0.007, Mitochondrion: 0.0046, Peroxisome: 0.002, Cytoplasm: 0.0009, Plastid: 0.0005, Extracellular: 0.0001 |
| | Q7JV09_DROME | Cell membrane, Membrane | 27,3 | 1,37 | 0 | LMLLVLIFNVALALLFVYML | 20 | T | 1 | https://www.uniprot.org/uniprot/Q7JV09 | SubName: Full=GH28348p {ECO:0000313| | Cell membrane: 0.4933, Golgi apparatus: 0.2348, Nucleus: 0.1593, Endoplasmic reticulum: 0.093, Lysosome/Vacuole: 0.0092, Cytoplasm: 0.0072, Extracellular: 0.0011, Mitochondrion: 0.0011, Peroxisome: 0.0009, Plastid: 0.0001 |
| | Q9VR90_DROME | Golgi apparatus, Membrane | 27,6 | 1,31 | 1 | MCVILVLAAVTFFMLLLLILT | 21 | KL | 2 | https://www.uniprot.org/uniprot/Q9VR90 | SubName: Full=Syntaxin 16 {ECO:0000313 | Golgi apparatus: 0.5052, Endoplasmic reticulum: 0.395, Lysosome/Vacuole: 0.0596, Cell membrane: 0.0213, Nucleus: 0.0071, Mitochondrion: 0.0065, Cytoplasm: 0.0022, Peroxisome: 0.0021, Plastid: 0.0011, Extracellular: 0.0 |
| | Q7JRF1_DROME | Golgi apparatus, Membrane | 27,9 | 1,33 | 0 | QWAAILILSGLLLFVIILFII | 21 | L | 1 | https://www.uniprot.org/uniprot/Q7JRF1 | SubName: Full=RE48509p {ECO:0000313| | Golgi apparatus: 0.4151, Endoplasmic reticulum: 0.3467, Lysosome/Vacuole: 0.1179, Nucleus: 0.0504, Cell membrane: 0.0403, Cytoplasm: 0.0135, Mitochondrion: 0.0106, Peroxisome: 0.0031, Plastid: 0.0022, Extracellular: 0.0002 |
| | Q7JY00_DROME | Golgi apparatus, Membrane | 27,9 | 1,33 | 0 | QWAAILILSGLLLFVIILFII | 21 | L | 1 | https://www.uniprot.org/uniprot/Q7JY00 | SubName: Full=HL02043p {ECO:0000313| | Golgi apparatus: 0.2715, Endoplasmic reticulum: 0.2264, Cell membrane: 0.211, Lysosome/Vacuole: 0.1143, Plastid: 0.1087, Nucleus: 0.0371, Mitochondrion: 0.0231, Peroxisome: 0.0033, Cytoplasm: 0.003, Extracellular: 0.0017 |
| | M9PG08_DROME | Endoplasmic reticulum, Membrane | 28,2 | 1,08 | 0 | LRFIVILPVLICFICTILSILILFQI | 26 | HT | 2 | https://www.uniprot.org/uniprot/M9PG08 | SubName: Full=Rho-related BTB domain c | Endoplasmic reticulum: 0.3353, Cell membrane: 0.3031, Golgi apparatus: 0.1994, Lysosome/Vacuole: 0.1376, Nucleus: 0.009, Mitochondrion: 0.008, Peroxisome: 0.004, Cytoplasm: 0.0034, Extracellular: 0.0001, Plastid: 0.0001 |
| | B7Z0E5_DROME | Endoplasmic reticulum, Membrane | 28,2 | 1,28 | 0 | YVLWTLLILIVILGLVLFFSNL | 22 | PAKLMTIPLE | 10 | https://www.uniprot.org/uniprot/B7Z0E5 | SubName: Full=Misfire, isoform F {ECO:00 | Endoplasmic reticulum: 0.5903, Cell membrane: 0.1549, Golgi apparatus: 0.1222, Lysosome/Vacuole: 0.0591, Nucleus: 0.043, Peroxisome: 0.0118, Cytoplasm: 0.0117, Mitochondrion: 0.0063, Extracellular: 0.0003, Plastid: 0.0003 |
| | A0A0B4K862_DROME | Cell membrane, Membrane | 28,4 | 1,14 | 3 | ILAAVIGGAVVGLLCAILVVMFIVY | 25 | RMRKKDEGSYALDEPKRSPANNSYAKNA | 35 | https://www.uniprot.org/uniprot/A0A0B4K862 | RecName: Full=Syndecan {ECO:0000256| | Cell membrane: 0.4612, Golgi apparatus: 0.1471, Lysosome/Vacuole: 0.1015, Endoplasmic reticulum: 0.0828, Mitochondrion: 0.0789, Plastid: 0.0706, Extracellular: 0.0499, Cytoplasm: 0.005, Nucleus: 0.0018, Peroxisome: 0.0011 |
| | A0A0B4LG79_DROME | Cell membrane, Membrane | 28,4 | 1,14 | 3 | ILAAVIGGAVVGLLCAILVVMFIVY | 25 | RMRKKDEGSYALDEPKRSPANNSYAKNA | 35 | https://www.uniprot.org/uniprot/A0A0B4LG79 | RecName: Full=Syndecan {ECO:0000256| | Cell membrane: 0.5399, Plastid: 0.1597, Golgi apparatus: 0.1356, Endoplasmic reticulum: 0.105, Mitochondrion: 0.0341, Lysosome/Vacuole: 0.0208, Nucleus: 0.0034, Peroxisome: 0.0013, Cytoplasm: 0.0, Extracellular: 0.0 |
| | SYB_DROME | Golgi apparatus, Membrane | 28,4 | 1,29 | 0 | MMIILGVIAVVLLIIVLVSVWP | 22 | SSSDSGSGGGNKAITQAPPH | 20 | https://www.uniprot.org/uniprot/P18489 | RecName: Full=Synaptobrevin; | Golgi apparatus: 0.5405, Endoplasmic reticulum: 0.2556, Cell membrane: 0.0931, Lysosome/Vacuole: 0.0601, Mitochondrion: 0.0243, Nucleus: 0.0106, Extracellular: 0.0105, Plastid: 0.0028, Cytoplasm: 0.0023, Peroxisome: 0.0003 |
| | Q8SXH7_DROME | Cell membrane, Membrane | 28,9 | 1,20 | 2 | IILVSIIMGITILVLITIALCYIA | 24 | YEKCQKKREYYINA | 14 | https://www.uniprot.org/uniprot/Q8SXH7 | SubName: Full=GEO09915p1 {ECO:000031 | Cell membrane: 0.4046, Golgi apparatus: 0.3036, Endoplasmic reticulum: 0.1163, Lysosome/Vacuole: 0.0717, Extracellular: 0.0632, Mitochondrion: 0.0345, Plastid: 0.0046, Nucleus: 0.0012, Peroxisome: 0.0002, Cytoplasm: 0.0001 |
| | M9PIV2_DROME | Golgi apparatus, Membrane | 28,9 | 1,45 | 0 | IMLIVILAAVLLVLLLVGIY | 20 | L | 1 | https://www.uniprot.org/uniprot/M9PIV2 | SubName: Full=Syntaxin 4, isoform B {ECO | Golgi apparatus: 0.519, Endoplasmic reticulum: 0.4212, Lysosome/Vacuole: 0.0291, Cell membrane: 0.0214, Nucleus: 0.0063, Mitochondrion: 0.0018, Cytoplasm: 0.0006, Peroxisome: 0.0005, Plastid: 0.0002, Extracellular: 0.0 |
| | STX4_DROME | Golgi apparatus, Membrane | 28,9 | 1,45 | 0 | IMLIVILAAVLLVLLLVGIY | 20 | L | 1 | https://www.uniprot.org/uniprot/Q7KVY7 | RecName: Full=Syntaxin-4; | Golgi apparatus: 0.582, Endoplasmic reticulum: 0.3317, Lysosome/Vacuole: 0.0376, Cell membrane: 0.0227, Nucleus: 0.0115, Mitochondrion: 0.01, Plastid: 0.0018, Cytoplasm: 0.0016, Peroxisome: 0.001, Extracellular: 0.0001 |
| | M9NDU9_DROME | Golgi apparatus, Membrane | 29,3 | 1,05 | 1 | LFFWAAVCCCVLTFLICLAICISIYIYF | 28 | IR | 2 | https://www.uniprot.org/uniprot/M9NDU9 | SubName: Full=Uncharacterized protein {E | Golgi apparatus: 0.4282, Endoplasmic reticulum: 0.2821, Cell membrane: 0.2689, Lysosome/Vacuole: 0.0204, Mitochondrion: 0.0003, Nucleus: 0.0, Cytoplasm: 0.0, Extracellular: 0.0, Plastid: 0.0, Peroxisome: 0.0 |
| | Q86P32_DROME | Golgi apparatus, Membrane | 29,7 | 1,49 | 0 | WAAILILSGLLLFVIILFII | 20 | L | 1 | https://www.uniprot.org/uniprot/Q86P32 | SubName: Full=RE35194p {ECO:0000313| | Golgi apparatus: 0.416, Endoplasmic reticulum: 0.3162, Lysosome/Vacuole: 0.1153, Nucleus: 0.062, Cell membrane: 0.054, Mitochondrion: 0.0172, Cytoplasm: 0.013, Plastid: 0.0035, Peroxisome: 0.0025, Extracellular: 0.0003 |
| | X2JGJ5_DROME | Endoplasmic reticulum, Membrane | 29,8 | 1,10 | 0 | IIGSFSMFVIFSYLQLFYILVAIYLAL | 27 | HHAHH | 5 | https://www.uniprot.org/uniprot/X2JGJ5 | SubName: Full=Uncharacterized protein {E | Endoplasmic reticulum: 0.8367, Golgi apparatus: 0.0857, Lysosome/Vacuole: 0.0431, Cell membrane: 0.023, Plastid: 0.0081, Mitochondrion: 0.0017, Peroxisome: 0.0009, Nucleus: 0.0006, Cytoplasm: 0.0002, Extracellular: 0.0001 |
| | H1UUB1_DROME | Golgi apparatus, Membrane | 29,9 | 1,42 | 0 | MMIILGVIAVVLLIIVLVSVW | 21 | PSSSDSGSGGGNKAITQAPPH | 21 | https://www.uniprot.org/uniprot/H1UUB1 | SubName: Full=GEO12465p1 {ECO:000031 | Golgi apparatus: 0.5324, Endoplasmic reticulum: 0.2632, Cell membrane: 0.076, Lysosome/Vacuole: 0.0389, Mitochondrion: 0.0374, Nucleus: 0.0254, Extracellular: 0.0218, Plastid: 0.0025, Cytoplasm: 0.0021, Peroxisome: 0.0003 |
| | Q9VGT2_DROME | Endoplasmic reticulum, Membrane | 30,3 | 1,17 | 9 | HSLDVLGTVLLVILLVIIAILLIIVF | 26 | SVCKISKNISTRLVVKKQRKQKVR | 24 | https://www.uniprot.org/uniprot/Q9VGT2 | RecName: Full=UDP-glucuronosyltransfera | Endoplasmic reticulum: 0.8515, Cell membrane: 0.0711, Golgi apparatus: 0.0684, Mitochondrion: 0.0041, Lysosome/Vacuole: 0.003, Peroxisome: 0.0011, Plastid: 0.0004, Nucleus: 0.0002, Cytoplasm: 0.0001, Extracellular: 0.0001 |
| | A4V0S1_DROME | Golgi apparatus, Membrane | 30,3 | 1,44 | 0 | WLMIKIFGVLIFFFLFFVVFM | 21 | S | 1 | https://www.uniprot.org/uniprot/A4V0S1 | SubName: Full=Syntaxin 5, isoform B {ECO | Golgi apparatus: 0.8341, Endoplasmic reticulum: 0.1194, Nucleus: 0.0271, Lysosome/Vacuole: 0.0079, Cell membrane: 0.0066, Mitochondrion: 0.0028, Cytoplasm: 0.0018, Peroxisome: 0.0002, Plastid: 0.0001, Extracellular: 0.0 |
| | STX5_DROME | Golgi apparatus, Membrane | 30,3 | 1,44 | 0 | WLMIKIFGVLIFFFLFFVVFM | 21 | S | 1 | https://www.uniprot.org/uniprot/Q24509 | RecName: Full=Syntaxin-5; | Golgi apparatus: 0.8341, Endoplasmic reticulum: 0.1194, Nucleus: 0.0271, Lysosome/Vacuole: 0.0079, Cell membrane: 0.0066, Mitochondrion: 0.0028, Cytoplasm: 0.0018, Peroxisome: 0.0002, Plastid: 0.0001, Extracellular: 0.0 |

| | | | | | | | | | | | |
|---|---|---|---|---|---|---|---|---|---|---|---|
| Q7K4J7_DROME | Endoplasmic reticulum, Membrane | 31,1 | 1,20 | -1 | LSWICLLIFILALCVVIPLVYVIYLA | 26 | EHPHHNHSAQ | 10 | https://www.uniprot.org/uniprot/Q7K4J7 | SubName: Full=LD36653p {ECO:0000313\| | Endoplasmic reticulum: 0.374, Golgi apparatus: 0.17, Nucleus: 0.1585, Cell membrane: 0.1559, Lysosome/Vacuole: 0.11, Cytoplasm: 0.0147, Mitochondrion: 0.0089, Peroxisome: 0.0047, Extracellular: 0.002, Plastid: 0.0012 |
| M9PCB0_DROME | Cell membrane, Membrane | 32,2 | 1,19 | 0 | FVAFIAAVACLGVIIGLILGIVTLVWF | 27 | KMTRDEETKKTYTGEGMPAEVKKND | 25 | https://www.uniprot.org/uniprot/M9PCB0 | SubName: Full=Uncharacterized protein {E | Cell membrane: 0.3673, Endoplasmic reticulum: 0.3247, Mitochondrion: 0.1007, Golgi apparatus: 0.0958, Plastid: 0.0687, Lysosome/Vacuole: 0.0389, Extracellular: 0.0024, Nucleus: 0.0008, Peroxisome: 0.0004, Cytoplasm: 0.0003 |
| C0PV11_DROME | Cell membrane, Membrane | 32,3 | 1,24 | 2 | MLLSIVFSALLMVVVIGLIVYFTVFY | 26 | HKDKNTDEVQKQVAQLTPIVKRSIRDYFN | 32 | https://www.uniprot.org/uniprot/C0PV11 | SubName: Full=MIP07124p {ECO:0000313 | Cell membrane: 0.3899, Golgi apparatus: 0.2958, Lysosome/Vacuole: 0.1065, Extracellular: 0.1045, Endoplasmic reticulum: 0.0535, Mitochondrion: 0.0436, Nucleus: 0.0032, Cytoplasm: 0.0017, Peroxisome: 0.0008, Plastid: 0.0006 |
| M9PFC0_DROME | Cell membrane, Membrane | 32,3 | 1,24 | 2 | MLLSIVFSALLMVVVIGLIVYFTVFY | 26 | HKDKNTDEVQKQVAQLTPIVKRSIRDYFN | 32 | https://www.uniprot.org/uniprot/M9PFC0 | SubName: Full=GEO09663p1 {ECO:000031 | Cell membrane: 0.3783, Golgi apparatus: 0.3257, Lysosome/Vacuole: 0.1059, Extracellular: 0.0827, Endoplasmic reticulum: 0.0524, Mitochondrion: 0.049, Nucleus: 0.003, Cytoplasm: 0.0015, Peroxisome: 0.0008, Plastid: 0.0007 |
| MEX1_DROME | Cell membrane, Membrane | 32,3 | 1,24 | 2 | MLLSIVFSALLMVVVIGLIVYFTVFY | 26 | HKDKNTDEVQKQVAQLTPIVKRSIRDYFN | 32 | https://www.uniprot.org/uniprot/P23487 | RecName: Full=Protein midgut expression | Cell membrane: 0.3783, Golgi apparatus: 0.3257, Lysosome/Vacuole: 0.1059, Extracellular: 0.0827, Endoplasmic reticulum: 0.0524, Mitochondrion: 0.049, Nucleus: 0.003, Cytoplasm: 0.0015, Peroxisome: 0.0008, Plastid: 0.0007 |
| A0A1Z1CH10_DROME | Lysosome/Vacuole, Membrane | 32,8 | 1,17 | -4 | CIVCSVVVCVLLIIAGFLIFILSVFAFA | 28 | PLVDKMQLRLIEDINVTTTERDPILDYDYG | 30 | https://www.uniprot.org/uniprot/A0A1Z1CH10 | SubName: Full=Uncharacterized protein, is | Lysosome/Vacuole: 0.3996, Cell membrane: 0.2091, Extracellular: 0.1054, Endoplasmic reticulum: 0.0925, Golgi apparatus: 0.0922, Plastid: 0.0709, Mitochondrion: 0.0261, Nucleus: 0.0027, Cytoplasm: 0.0014, Peroxisome: 0.0 |
| M9PJN0_DROME | Golgi apparatus, Membrane | 32,8 | 1,56 | 2 | MLLFIGLIILIVFFLYYILYI | 21 | SGINVIFKSRSPIVY | 15 | https://www.uniprot.org/uniprot/M9PJN0 | SubName: Full=Mushroom body defect, is | Golgi apparatus: 0.8299, Endoplasmic reticulum: 0.1599, Cell membrane: 0.0034, Lysosome/Vacuole: 0.0033, Nucleus: 0.0014, Cytoplasm: 0.0013, Mitochondrion: 0.0007, Extracellular: 0.0, Plastid: 0.0, Peroxisome: 0.0 |
| Q8IR54_DROME | Golgi apparatus, Membrane | 32,8 | 1,56 | 2 | MLLFIGLIILIVFFLYYILYI | 21 | SGINVIFKSRSPIVY | 15 | https://www.uniprot.org/uniprot/Q8IR54 | SubName: Full=Mushroom body defect, is | Golgi apparatus: 0.8344, Endoplasmic reticulum: 0.1542, Cell membrane: 0.0041, Lysosome/Vacuole: 0.0036, Nucleus: 0.0015, Cytoplasm: 0.0014, Mitochondrion: 0.0008, Extracellular: 0.0, Plastid: 0.0, Peroxisome: 0.0 |

**Predicted TA proteins** 254

Tested TA proteins
